# Neural representational geometry of a joint code for stimulus category and category-independent features

**DOI:** 10.64898/2026.03.23.713692

**Authors:** Lorenzo Tiberi, Haim Sompolinsky

**Affiliations:** Center for Brain Science, Harvard University, Cambridge, MA 02138; Edmond and Lily Safra Center for Brain Sciences, Hebrew University, Jerusalem 9190401, Israel

**Keywords:** multivariable population coding, representational geometry, deep neural networks, ventral visual stream

## Abstract

A central question in neuroscience and machine learning is how a single neural representation can support linear access to multiple kinds of information about the same stimulus. While this ability is widespread in biological and artificial systems, the representational principles that make such joint coding possible remain poorly understood. We address this question for an important class of representations: those jointly encoding discrete stimulus categories and continuous features that vary independently of category. Using the framework of category manifolds—the sets of neural representations elicited by stimuli from the same category—we extend existing theories of manifold classification to the equally essential task of regressing category-independent features, determining which aspects of manifold geometry govern regression performance. This provides a unified framework for understanding how classification- and regression-relevant geometry can be jointly optimized to implement an effective joint code. Applying this framework to convolutional neural networks (CNNs), we find that regression-relevant geometry can be optimized through subtle changes that largely preserve classification-relevant geometry. This explains why common representational-similarity measures previously failed to distinguish joint codes from codes optimized exclusively for classification. Motivated by prior work in visual neuroscience suggesting that macaque inferotemporal cortex may jointly encode object category and category-independent features such as object position and size, we use our framework to identify principled geometric signatures that distinguish joint codes from classification-only codes in CNNs and can be tested in future neural recordings. Finally, we characterize how these signatures are affected by common experimental constraints: limited stimulus categories and neural-population subsampling.

---

A hallmark of natural intelligence is the ability to extract multiple kinds of information from the same sensory stimulus. In everyday vision, for example, animals routinely extract from the same visual input both object identity and identity-independent attributes such as object position, size, pose, and texture. A central question in both neuroscience and machine learning is how neural networks may implement this ability.

A widespread solution appears to be the joint encoding of multiple stimulus-related variables in a shared high-level representation, from which these variables can be accessed by simple readouts. In machine learning, this principle is reflected in the growing pursuit of general-purpose representations that serve as a common feature space for multiple downstream tasks, with each task supported by a simple, often linear, readout head (1–4). In neuroscience, neural recordings in higher-order brain areas, such as macaque prefrontal cortex, anterior cingulate cortex, and hippocampus (5, 6), show that multiple task-relevant variables can be linearly decoded from the same population activity.

These observations have raised the question of which geometric properties of neural representations allow multiple variables to be jointly encoded in a linearly accessible form (7). Existing theoretical accounts have mainly addressed highly controlled regimes with low-dimensional stimulus spaces and task variables (5, 6, 8–11), or have focused on binary classification tasks (12). Much less is understood in regimes with many stimulus categories exhibiting high-dimensional within-category variability, and in tasks in which continuous variables must be encoded alongside discrete variables such as stimulus category.

A particularly natural setting for this regime is vision, where single-object images can span many object categories and, within each category, vary across a wide range of category-preserving transformations. In artificial vision, modern object detectors such as DETR (DEtection with TRansformers) (13) provide a concrete example in which a shared representation supports prediction, through simple readout heads, of both an object’s category and continuous object features such as its bounding-box coordinates. In biological vision, it has long been suggested that neural representations in primate higher-order visual areas encode not only object category, but also a variety of object features (14–16), including size (17, 18), texture and material properties (19–21), shape properties (22, 23), and facial properties (24, 25).

A seminal study (26) systematically probed this joint-coding hypothesis by showing that, along the macaque ventral stream (V1→V4→inferior temporal cortex, IT), linear readout performance improves not only for object category, consistent with the well-established role of the ventral stream in object recognition, but also for category-independent features such as object position, size, and pose within the image. These features are traditionally considered nuisance variables to be discarded in the service of invariant object recognition. This suggests that the same population in IT may in fact implement a joint code for category and category-independent information. Yet, the current empirical evidence raises several questions that call for a more principled understanding of joint codes of this kind. For example, both previous work (26) and our analysis show that linear regression performance for category-independent features can improve even across the layers of convolutional neural networks (CNNs) trained exclusively for object classification, despite performance remaining limited even in the top layers. Thus, improvement in regression performance along a visual hierarchy does not by itself distinguish a genuine joint code from a representation optimized only for object classification. The macaque recordings (26, 27) present the same ambiguity: although regression performance improves along the ventral stream, it remains limited in the recorded IT population, raising the question of whether this simply reflects experimental constraints, such as neural population subsampling and limited stimulus sets, or a genuine limitation of the underlying representation in implementing a joint code. Even with access to the full neural population, however, characterizing joint codes—particularly how they differ from classification-only codes—appears nontrivial: recent CNN results (28) show that networks trained either only for object classification or jointly for object classification and regression of category-independent features remain highly similar under common representational similarity measures such as centered kernel alignment (CKA), as well as in their alignment to neural data.

Motivated by these observations, in this work we ask the following fundamental questions: What geometric properties of a representation determine linear classification performance for object category and linear regression performance for category-independent features? How can regression-relevant geometry be optimized alongside classification-relevant geometry to implement an effective joint code? What geometric signatures characterize such a joint code, and how do they differ from those of a classification-only code? And how do these signatures behave under common experimental constraints?

For object classification, theories already exist that relate the geometry of population representations to proxy measures of classification performance, such as storage capacity (29) and few-shot generalization (30). Examples of such geometric measures include the radius and dimensionality of category manifolds, defined as the sets of population responses elicited by images from the same category. What is missing is an analogous theory that identifies the manifold geometry relevant for regression of category-independent features encoded within each category manifold. Here we develop such a theory and test it in CNNs trained for either object classification, regression of category-independent features, or a combination of these two objectives.

In what follows, we first identify two distinct contributions to the regression error of category-independent features: a local error, reflecting the quality of linear feature encoding within individual category manifolds, and a local–global error gap, reflecting how consistently these encodings are organized across categories to support category-independent regression. Applying this regression error decomposition to CNNs, we find that, while even CNNs trained exclusively for classification can reduce the local error along the visual hierarchy, only regression-optimized CNNs strongly suppress the local–global gap, making a small local–global gap a key geometric signature of joint coding in these networks. We then theoretically identify the manifold-geometric properties that determine this error gap and characterize how these properties are reorganized in joint codes relative to classification-only codes implemented by our CNNs. We further show how this regression-relevant geometry can be optimized with only minor changes to coarser, classification-relevant manifold properties such as radius and dimensionality. Thus, substantial changes in regression-relevant geometry can coexist with high global representational similarity: joint and classification-only codes can appear similar under CKA yet remain distinguishable by the finer measures identified by our theory. Finally, because a small local– global error gap appears as a key signature distinguishing joint from classification-only codes in our CNNs, we characterize how reliably this gap can be estimated under common experimental constraints. We show that a limited number of stimulus categories can lead to a strong underestimation of the local–global gap, an effect that our theory can correct by extrapolating to the many-category limit. By contrast, subsampling neural units causes the error gap in joint codes to approach that of classification-only codes, erasing the distinction between them under this key metric and suggesting that future experiments probing joint coding in the ventral stream may require larger-scale neural recordings.

## Results

### 1. Linear decoding framework

To assess how well neural representations encode both an image’s object category and features that vary independently of category (“category-independent” features), we adopt the linear-decoding framework of (26) (**Fig. 1**). We consider image datasets consisting of single-object images labeled by category and by four continuous bounding-box parameters describing object position and size: center coordinates *C*_*h*_, *C*_*v*_ and side lengths *L*_*h*_, *L*_*v*_ along the image’s horizontal (h) and vertical (v) axes (**Fig. 1a**). For each image, an *N* -dimensional neural representation vector is extracted from the activations of neurons in the brain visual hierarchy or from the activations of a CNN that mimics this hierarchy (**Fig. 1b**). Finally, we evaluate the performance of linear decoders in extracting category and category-independent information from these representations, treating it as a proxy for the decoding performance achieved by hypothetical downstream neurons (**Fig. 1c**). For category, we train one-vs-rest classifiers and evaluate their balanced classification accuracy, averaged across categories. For the bounding box parameters, we train one linear regressor per parameter and evaluate it by normalized mean squared error (nMSE), i.e., MSE divided by the parameter variance. All decoding metrics are cross-validated over multiple train–test splits.

**Fig. 1.**
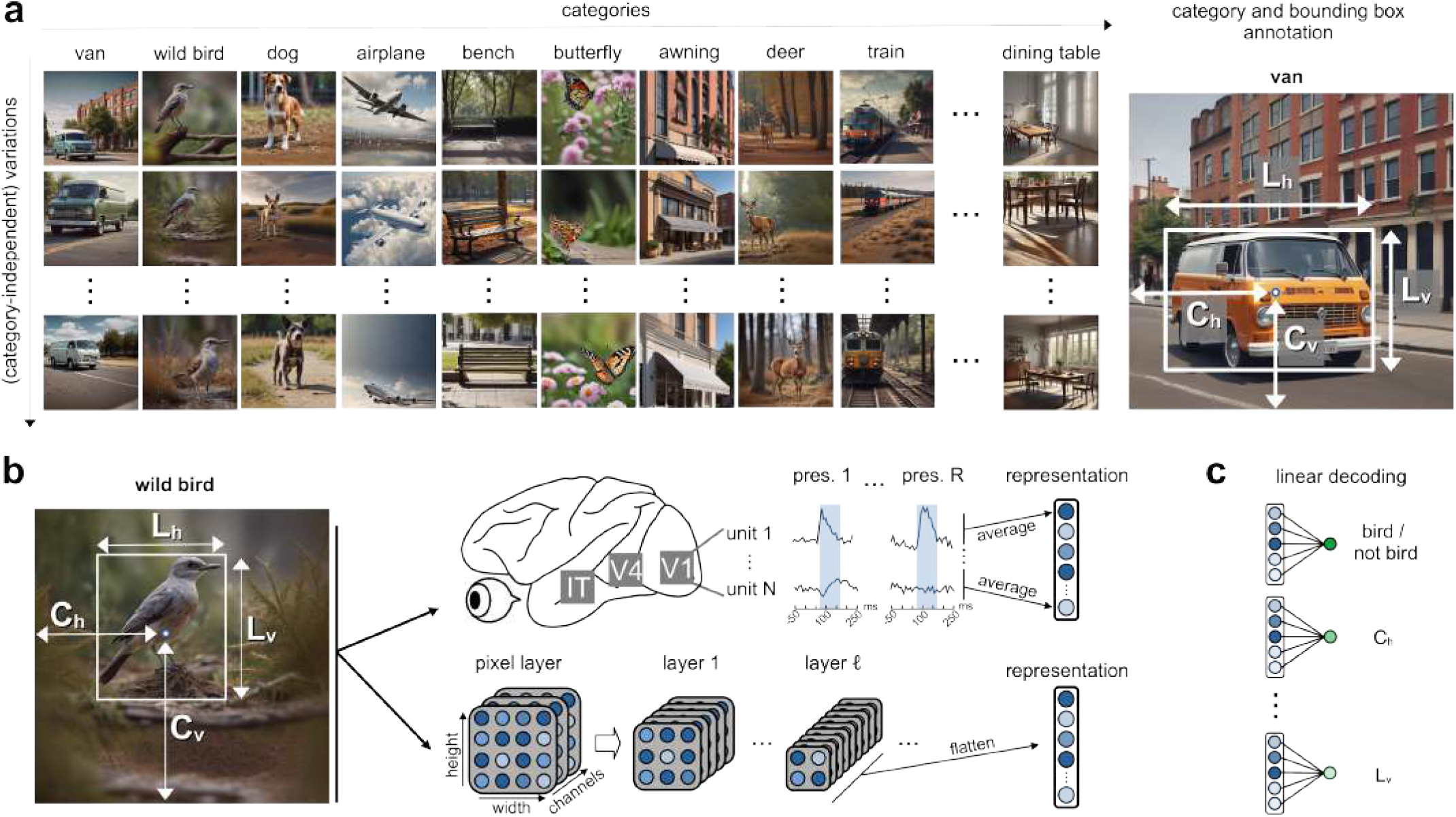
Framework to quantify how well neural representations encode category and category-independent information. (**a**) Overview of the main image dataset used in this work. The dataset contains single-object images annotated with object category and four bounding box parameters *C*_*h*_, *C*_*v*_, *L*_*h*_, *L*_*v*_. (**b**) From image to neural representation. An image is presented to the ventral visual stream (top), modeled in this work by a convolutional neural network (CNN; bottom), and the resulting neural representations are analyzed. Top: Representations from neural recordings are obtained by averaging spike counts over a post-timulus time window (shaded region) and across stimulus repetitions. Bottom: Representations from a given CNN layer are obtained by flattening that layer’s activations into a one-dimensional vector. (**c**) We evaluate the ability of linear decoders to extract category and bounding-box parameters from these representations (see Materials and Methods for details).

### 2. One code supports both classification and regression

We have applied this decoding framework to the publicly available dataset of macaque recordings (27) studied in (26), reproducing their qualitative results (see SI): decoding performance of both category and category-independent features improves along the visual hierarchy. However, we also find that absolute regression performance remains limited: root nMSE remains above 0.5, meaning that the root-mean-square prediction error is still larger than half the across-image standard deviation of the decoded feature.

To investigate what it takes to accurately perform both categorization and regression from a single population representation, we apply the above decoding framework to CNNs trained on a newly constructed large-scale dataset of single-object images with category and bounding-box annotations (**Fig. 1a**). This setup provides a controlled empirical testbed for our theory that is largely free of the experimental constraints of neural recordings: all units in the artificial networks are accessible, while the large number of stimulus categories in our dataset greatly reduces limitations due to category subsampling. These constraints can then be reintroduced in a controlled manner later in our analysis. Although the theory developed below is general and can be applied to any population representation, we chose to test it specifically on a CNN architecture because of its direct relevance to one of the empirical motivations for our study: the hypothesis of a joint code in the macaque ventral stream (26, 28). Indeed, CNN representations have been extensively compared with those of the primate ventral stream and shown substantial alignment with ventral-stream neural responses (31, 32).

Our dataset spans 265 object categories, with 20,000 images per category: 10,000 for training the CNN and 10,000 for generating neural representations and evaluating linear decoders. Images were generated with a multi-stage pipeline that uses diffusion models (33, 34) to produce photorealistic single-object images with a controlled distribution of bounding-box parameters (see Materials and Methods). Compared with previous approaches based on rendered 3D object models in random environments (27, 28), our method produces both more object categories and more photorealistic images of objects embedded in contextually relevant environments, albeit at the cost of losing explicit control over object pose.

On this dataset, we fine-tuned an ImageNet-1k–pretrained ResNet-50 (**Fig. 2a**). We treated the final feature layer as the shared representation and attached two types of readout: (i) the standard linear classification head for predicting object category, and (ii) four additional linear regression heads, each predicting one bounding-box parameter (*C*_*h*_, *C*_*v*_, *L*_*h*_, *L*_*v*_). The network was optimized end-to-end using stochastic gradient descent on a combined loss: cross-entropy loss on the classification head and Smooth L1 loss on each regression head (see Materials and Methods). In addition to this model, which we call network CR, we also trained the same architecture exclusively on the classification task (network C) and exclusively on the regression task (network R). The C and R networks provide empirical ceilings on the performance achievable on this dataset by this architecture, when it is allowed to specialize in a single task.

**Fig. 2.**
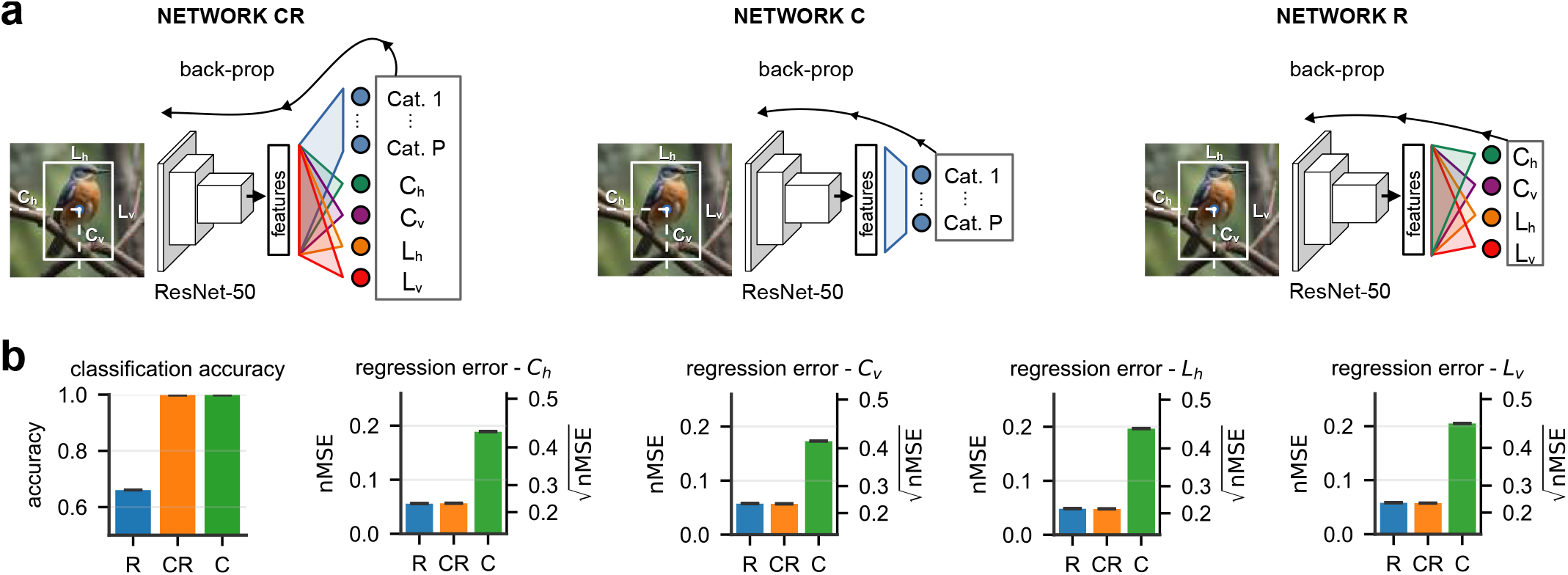
One code supports both object classification and regression of category-independent features. (**a**) The three ResNet-50 variants trained end-to-end on object classification and bounding-box regression jointly (CR), classification alone (C), or regression alone (R). (**b**) Linear decoding of object category and bounding-box parameters from the feature-layer representations of networks R, CR, and C. Error bars indicate SEM across cross-validation (CV) splits. Regression performance is reported as 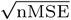 (left axis) and nMSE (right axis).

**Fig. 2b** shows the performance of linear decoders trained on the feature-layer representations of these three networks. Network CR matches the performance of network C on classification and that of network R on regression. We thus conclude that network CR implements an effective joint code that supports linear readout of both object category and category-independent features.

### 3. Theory

What properties of the population code of network CR, or of any population code in general, allow this level of performance in both object classification and regression of category-independent features? For classification, theories already exist that relate neural representation geometry to proxy measures of linear classification performance, such as storage capacity (29) or few-shot learning accuracy (30). Such theories are formulated within the framework of category manifolds, defined as the collection of all neural representations of images belonging to the same category (**Fig. 3a**). It has been shown that along the hierarchy of both classification-trained CNNs (35) and the ventral visual stream (30), such object manifolds change their geometric properties, such as their dimensionality, radius, and centroid separation, so as to become more and more disentangled, thereby improving linear classification performance (**Fig. 3a**). In a system optimized also for regression, one could imagine a similar scenario whereby, along the visual hierarchy, the encoding of category-independent features becomes both more linear within each manifold and more consistently aligned across manifolds, such that it can be regressed accurately by a single category-independent regressor (**Fig. 3a**). Unlike for classification, however, a theory identifying and linking the relevant category-manifold geometry to linear regression performance is currently lacking. We now develop such a theory.

**Fig. 3.**
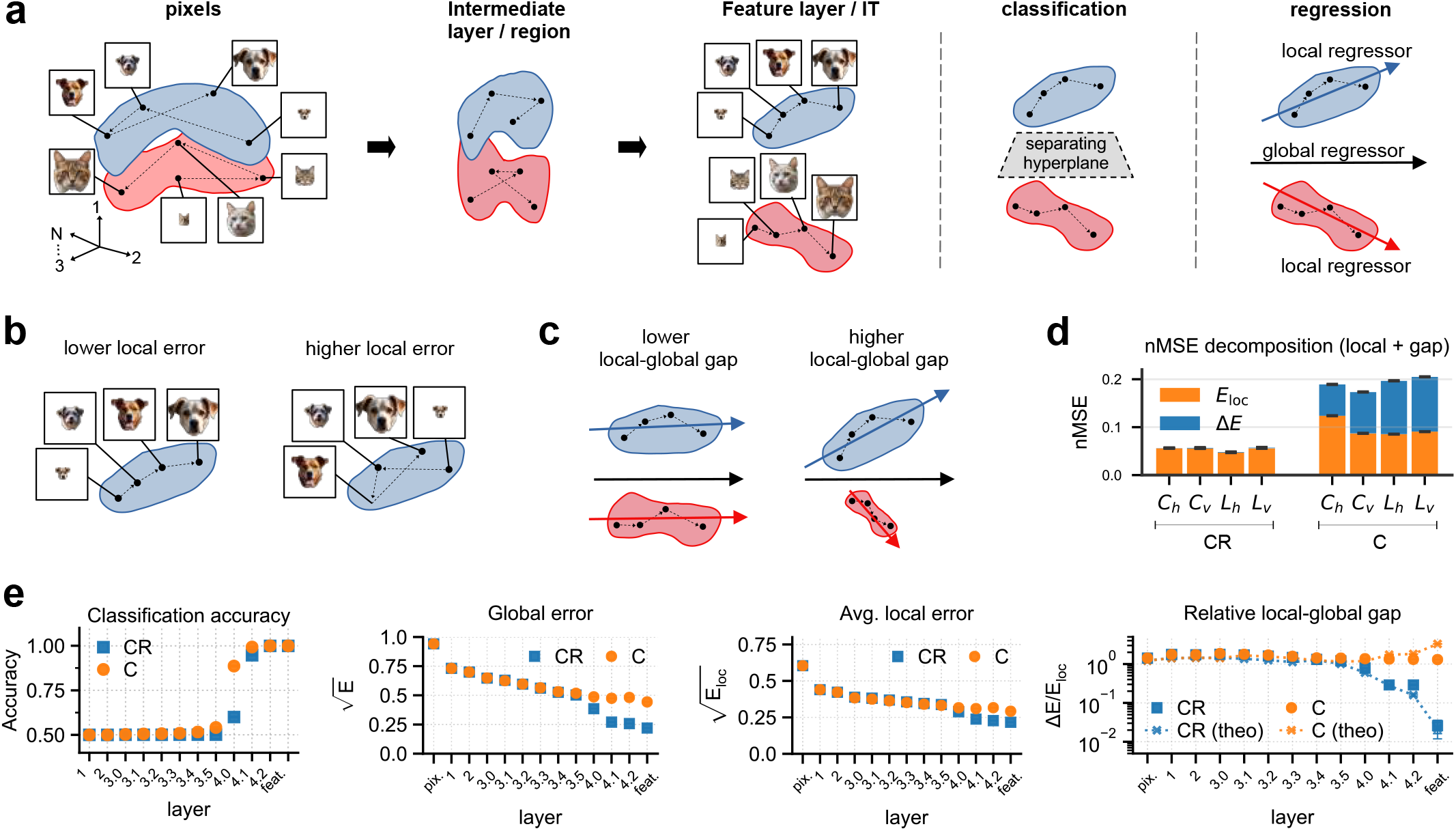
Theoretical framework and regression error decomposition. (**a**) Category manifolds comprise the neural representations elicited by images from the same category (dog, blue; cat, red). Dots denote individual images, and arrows order representations by increasing object size. Joint optimization is schematized as greater manifold disentanglement for classification, together with a feature encoding that is both more linear and more consistent cross-categories for regression. (**b**) The local error *E*_loc_ reflects the linearity of feature encoding within individual manifolds. (**c**) The local–global gap Δ*E* is the additional error incurred by using a single category-independent regressor rather than category-specific regressors; greater consistency in scale and orientation of the feature encoding across manifolds reduces this gap.(**d**) Decomposition of the nMSE into local error *E*_loc_ (orange) and local–global gap Δ*E* (blue) for regression of the four bounding box parameters from the feature layer of networks C and CR. Note that Δ*E* is very small and barely visible for network CR. (**e**) Object-classification accuracy and regression of *L*_*h*_ across network layers. Regression is shown as global error *E*, local error *E*_loc_, and relative gap Δ*E/E*_loc_. For the relative gap, we additionally overlay the theoretical prediction developed in the next section (crosses and dotted lines). Readouts from pixels and intermediate layers used random projection to 2,048 dimensions, matching the feature layer; results without random projection and for the other bounding-box features are analogous (see SI). For panels (d-e), Error bars indicate SEM across CV splits and, where applicable, five projection seeds.

### 3.1. Two sources of regression error

We begin by distinguishing two additive contributions, of very different nature, that determine linear regression performance for category-independent features (**Fig. 3b-c**). Let *E* denote the test MSE obtained when linearly regressing a given category-independent feature with a single category-independent readout. We decompose it as

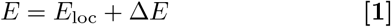

where *E*_loc_ = Avg_*µ*_ [*E*^*µ*^], with *E*^*µ*^ the test MSE achieved by a regressor trained and tested exclusively on the *µ*-th category. The “local error” *E*_loc_ quantifies the quality of the feature’s linear encoding at the *local* level of a single category, i.e. within a single object manifold (**Fig. 3b**). It provides a lower bound on the global error *E*, corresponding to the regression error of an oracle that is allowed to use separate regressors for each category. The residual Δ*E*, termed the “local-global error gap”, captures the additional error incurred by requiring a single category-independent readout, and reflects the global geometric arrangement of object manifolds across categories in the *N* -dimensional space of neural units (**Fig. 3c**).

**Fig. 3d** shows the decomposition *E* = *E*_loc_ + Δ*E* for the MSE of regressing object bounding box parameters from the feature layer of networks C and CR. Relative to network C, network CR improves both the local error *E*_loc_ and the local–global gap Δ*E*. The dominant improvement, however, is in Δ*E*, which drops by orders of magnitude in network CR. A small relative local–global gap Δ*E/E*_loc_ therefore emerges as a particularly distinctive signature of a joint code in these networks. Notably, we find that a small Δ*E/E*_loc_ emerges even in a network, termed CRloc, trained for classification and category-specific regression, without the additional constraint of a shared category-independent regressor (see SI).

This decomposition of the regression error becomes even more informative when examined across the layers of the two networks (**Fig. 3e**). As also observed in previous work (26), total regression error *E* improves even across the layers of network C. A key piece of evidence for a joint code in macaque IT has been the observation that linear readout performance for both object category and category-independent features improves along the ventral stream, although regression performance remained limited even in IT, potentially because of experimental constraints (26). Our results show, however, that while improved regression performance along a visual hierarchy is consistent with a joint code, it is not by itself diagnostic of one, since the same improvement can arise in a network optimized exclusively for object classification. Crucially, our regression error decomposition reveals a key difference between the two networks: in network C, the improvement in regression performance is driven exclusively by a reduction in the local error *E*_loc_, whereas an orders-of-magnitude reduction in the relative gap Δ*E/E*_loc_ emerges only in network CR. This further establishes a small relative local–global gap as a particularly distinctive signature of joint coding in these networks.

The origin of the local-error improvement in network C, despite the absence of explicit regression optimization, is beyond the scope of the present work. One possible explanation is that estimating an object’s bounding box depends on first recognizing the object, such that linearization of bounding-box features may emerge as a by-product of improved object classification. This idea would also be consistent with the observation that, in network CR, regression performance improves sharply only in the final layers, concurrently with a sharp rise in classification accuracy. Spatial inductive biases from convolution may also contribute, as we do not observe similarly clear across-layer improvements in the regression of additional features such as mean luminance, local contrast, and color saturation (see SI). However, other explanations are also possible (36).

### 3.2. Geometry of feature organization across categories

To gain insights into the manifold-geometry reorganization reflected by Δ*E*, we develop a theory that predicts Δ*E* from readily interpretable geometric measures. Under fairly general assumptions on manifold statistics, in the limit of infinitely many categories, and neglecting higher-order corrections in 1*/N* where *N* is the number of neurons, we show (see Materials and Methods and SI) that

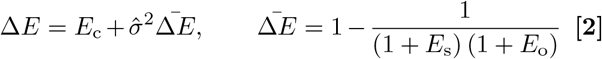

where *E*_s_, *E*_o_ and *E*_c_ represent three distinct sources of error:

#### 1. Centroid error (*E*_c_)

The term *E*_c_ reflects the error in fitting the manifold centroids (**Fig. 4a**, left). Each category *µ* has a centroid (mean representation) 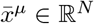 and a mean feature value 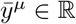. A single category-independent regressor must be able to fit the mapping 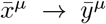. Operationally, *E*_c_ is the part of Δ*E* that would be eliminated by centering each category manifold (and the corresponding feature labels) around its category mean before fitting and evaluating the category-independent regressor (Materials and Methods). The remaining term 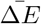 represents the error due to fitting manifold variability around the centroids.

**Fig. 4.**
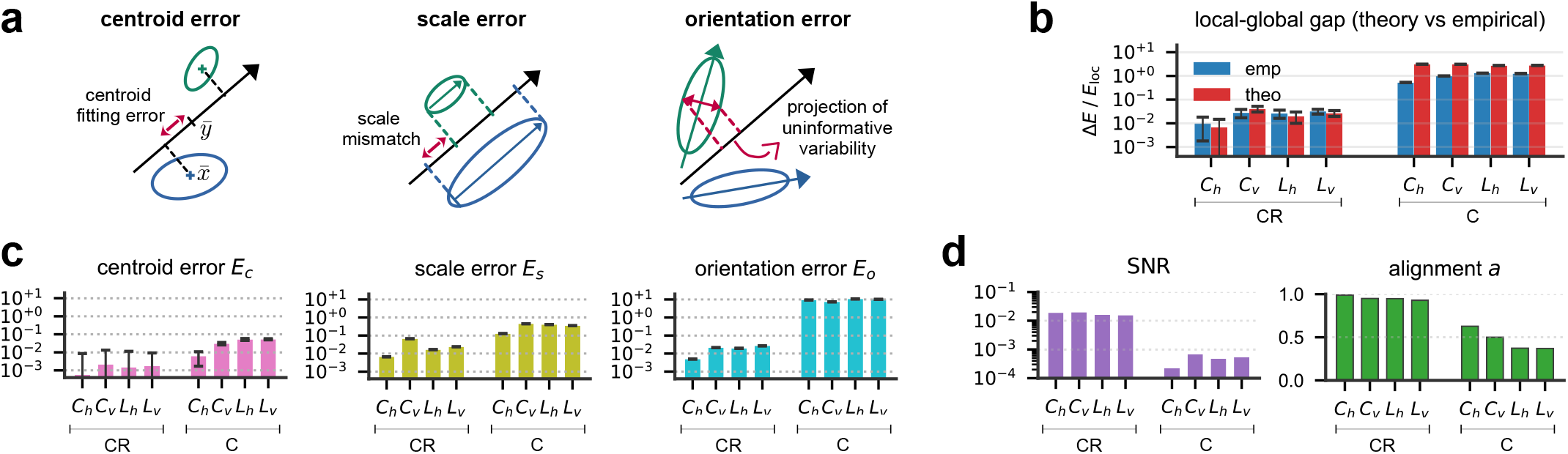
Theory for the local-global error gap. (**a**) Schematic illustration of the three sources of error (*E*_c_, *E*_s_, *E*_o_) contributing to Δ*E*. Left: The centroid error *E*_c_ reflects the error in fitting the manifold centroids. Center: The scale error *E*_s_ arises from variability in the scale of the local feature encoding across categories. Right: The orientation error *E*_o_ arises from misalignment of the local feature-encoding direction across categories, causing the projection of uninformative manifold variability onto the global regressor. (**b**) Measured relative local-global gap Δ*E/E*_loc_ (blue) compared to our theory prediction (red). (**c**) Decomposition of the relative gap Δ*E/E*_loc_ into the contributions from the centroid error *E*_c_ (pink), scale error *E*_s_ (olive), and orientation error *E*_o_ (cyan). All three errors are normalized by dividing by *E*_loc_, and scale and orientation errors are further multiplied by 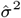. This normalization allows for a more direct comparison of the three errors, since, to linear order, 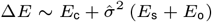. In panels (b-c), error bars denote propagated SEM across CV splits. (**d**) The two elements contributing to the orientation error: SNR, and alignment *a* of the local feature-encoding directions.

#### 2. Scale error (*E*_s_)

The scale error *E*_s_ reflects an across-category mismatch in the scale at which the feature is encoded within each manifold (**Fig. 4a**, center). Let *W*^*µ*^ ∈ ℝ^*N*^ denote the optimal local readout vector regressing the feature in category *µ*, and define the scale of the local encoding as ∥*W*^*µ*^∥. We then define

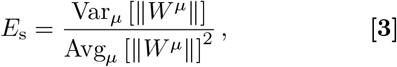

which is the squared coefficient of variation of ∥*W*^*µ*^∥ across categories (and vanishes only when all categories encode the feature with the same scale).

#### 3. Orientation error (*E*_s_)

The orientation error *E*_s_ reflects across-category misalignment of the local encoding directions *ŵ*^*µ*^ = *W*^*µ*^*/* ∥*W*^*µ*^∥. Misalignment causes within-manifold variability that is uninformative for the feature to project onto the global category-independent regressor, increasing error (**Fig. 4a**, right). We obtain

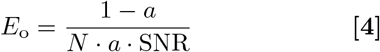

where *a* = ‖Avg_*µ*_ [*ŵ*^µ^] ‖^2^ quantifies the alignment of the local encoding directions (*a* = 0: no net alignment; *a* = 1: perfect alignment) and SNR = Avg_*µ*_ [*ŵ*^*µ*⊤^Σ^*µ*^*ŵ*^*µ*^ */*Avg_*µ*_ [TrΣ^*µ*^] measures the signal-to-noise ratio between the (informative) variance along the local encoding direction and the total (uninformative) manifold variance. Here Σ^*µ*^ ∈ ℝ^*N*×*N*^ is the centered covariance matrix of representations belonging to the *µ*-th manifold. The *N* ^−1^ scaling arises because the expected squared projection of uninformative within-manifold variability onto the global regressor scales as *N* ^−1^. Note that this represents a high-dimensional advantage: lower values of alignment *a* or SNR are sufficient to obtain the same *E*_o_ as the dimensionality *N* of the neural space increases. Therefore, in high dimensions, the geometric constraints for achieving accurate category-independent regression are more relaxed. In particular, accurate regression need not satisfy the naive expectation that the encoding of a feature should align tightly along a specific direction across categories. In a joint code, this flexibility may allow the degree of alignment to be more freely adjusted, for example to leave room for the optimization of other geometric properties that may serve other tasks.

Finally, 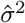 is the variance of the feature explained by the optimal local linear readout, averaged across categories, and converts the dimensionless gap 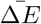 into MSE units.

**Fig. 4b** compares our theoretical prediction with the measured local–global error gap Δ*E* at the feature layer of networks C and CR. The agreement is excellent for network CR and more approximate for network C. Nevertheless, the theory correctly captures the orders-of-magnitude reduction in Δ*E* from network C to CR. Moreover, as shown in **Fig. 3e**, the theoretical prediction closely tracks the measured gap across all other layers of both networks, with good quantitative agreement throughout. **Fig. 4c** further decomposes the theoretical prediction of Δ*E* into contributions from the centroid error *E*_c_, scale error *E*_*s*_, and orientation error *E*_o_. Compared with network C, all three error terms are reduced in network CR, with *E*_s_ and *E*_o_ in particular reaching comparably small values, with neither term dominating. **Fig. 4d** shows the two factors contributing to the orientation error *E*_o_: the SNR and the alignment index *a*. Both are improved in network CR compared with network C. Analogous results for the behavior of *E*_c_, *E*_*s*_, *E*_o_, *a* and SNR across the layers of networks C and CR are provided in the SI.

### 4. Manifold geometry reveals optimization strategy

In the previous section, we identified the manifold geometry relevant to regression performance and showed how network CR optimizes this geometry relative to network C. To understand the joint coding implemented by network CR, we next ask two questions. How can this regression-relevant geometry be optimized alongside classification-relevant geometry to implement an effective joint code? And how does this joint optimization affect the geometry of the representations, particularly compared with that of a code optimized only for classification? The latter question is particularly important in light of previous findings (28) that representations in joint and classification-only codes are nearly indistinguishable under standard measures of representational similarity such as centered kernel alignment (CKA). Our theoretical framework provides insight into both questions. We find that, because regression-relevant geometry is mostly associated with a specific direction within each manifold—the feature-encoding direction—rather than with the manifold as a whole, it can be optimized *within* a manifold while largely preserving overall manifold shape and, consequently, classification-relevant properties such as manifold radius and dimensionality. As a result, differences in manifold geometry between joint and classification-only codes can be subtle enough to be largely missed by coarse measures of representational similarity such as CKA.

To see how this can occur, consider network C, which is optimized only for classification and is represented schematically on the left of **Fig. 5a**. In this network, category-independent features are encoded along relatively low-variance directions within the manifolds, resulting in low SNR, and their encodings are poorly matched across categories in both scale and direction, resulting in high scale error *E*_*s*_ and low alignment *a*. One might intuitively think that a network implementing a joint code would optimize this regression-relevant geometry through the following strategy (**Fig. 5a**, strategy a): increasing variance along the feature-encoding directions and suppressing variance along other manifold directions, thereby increasing SNR; and aligning these encodings in both scale and direction across manifolds, thereby reducing *E*_*s*_ and increasing *a*. Under this strategy, the regressed features would be encoded along the dominant principal components (PCs) of each manifold, and the alignment of these dominant directions across categories would cause the manifolds themselves to become strongly aligned. As we show below, this is indeed the strategy adopted by network R, which is optimized exclusively for regression.

**Fig. 5.**
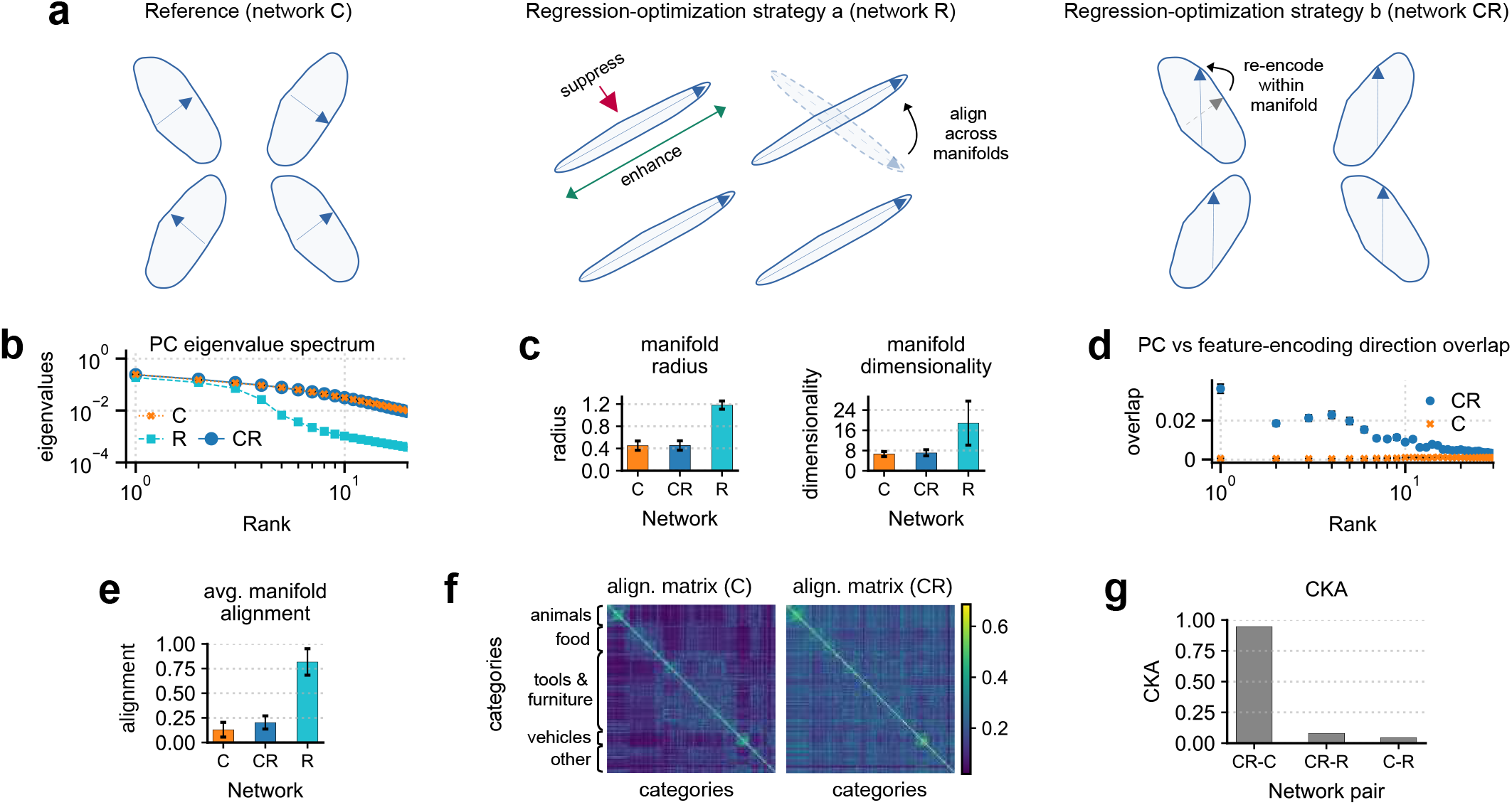
Geometry optimization strategy. (a) Schematic depiction of the two strategies to optimize regression-relevant geometry. (b) PC eigenvalue spectrum (eigenvalue vs rank) for category manifolds in network C (orange), CR (blue), and R (cyan), averaged across categories (error bars are SEM). For each manifold, the eigenvalues were normalized by the manifold total variance averaged across networks. Analogous per-category comparisons of the PC spectra are provided in the SI. (c) Manifold radius and dimensionality, as defined by manifold-capacity theory (29, 35), averaged across categories (error bars are SD). Under this definition, lower dimensionality corresponds to greater category-storage capacity. Dimensionality defined as the participation ratio of the PC eigenvalues is reported in the SI; it is likewise similar in networks C and CR but higher than in R, a trend associated with better few-shot generalization (30). Thus, both lower manifold-capacity dimensionality and higher participation-ratio dimensionality favor different aspects of classification. (d) Squared cosine of the angle between the feature-encoding direction *ŵ*^*µ*^ and the PC directions of manifold *µ* (ordered by rank) for networks C (orange) and CR (blue), averaged across categories (error bars are SEM). We show the case of feature *L*_h_. For analogous results for the other features, see the SI. (e) Manifold alignment, as quantified by the RV coefficient between pairs of manifold covariance matrices, averaged across all pairs of categories (error bars are SD). Alignment can range from 0 to 1, with 1 meaning perfect alignment. For a review of the RV coefficient definition, see the SI. (f) Matrix of pairwise manifold alignments, for network C (left) and network CR (right). Categories are ordered (and grouped) by semantic similarity; macro-groups are indicated on the left. (g) Linear CKA between representations in pairs of networks, computed over the full set of images across all categories as in (28). CKA can range from 0 to 1, with 1 meaning perfect similarity. For a review of the CKA definition, see the SI. Analogous results for CKA performed on single categories are reported in the SI.

However, this strategy substantially alters manifold shapes relative to those in network C, which has presumably optimized these shapes to support classification. Indeed, we show below that network CR, which implements a joint code supporting both regression and classification, adopts a very different strategy (**Fig. 5a**, strategy b): it re-encodes the features *within* each manifold while leaving both manifold shapes and their relative orientations largely unchanged. Specifically, the features are re-encoded along higher-variance directions within each manifold, thereby increasing SNR without substantially reshaping the manifolds; and this within-manifold re-encoding also ensures that features become more aligned across categories in both scale and direction, thereby improving both *E*_s_ and *a* without requiring the manifolds themselves to rotate into strong alignment.

The remaining panels of **Fig. 5** show several measures of manifold geometry that support this picture. **Fig. 5b** shows that the manifold PC spectra—used here to characterize manifold shape—are essentially unchanged in network CR relative to network C. By contrast, network R drastically reshapes the manifolds relative to networks C and CR, concentrating most of their variance along approximately four principal directions—matching the number of regressed category-independent features. Consequently, manifold radius and dimensionality—two key geometric properties relevant to classification performance—are virtually unchanged between networks C and CR, but differ markedly in network R (**Fig. 5c**). Even though manifold shapes remain largely unchanged in network CR compared to C, the feature-encoding direction shows increased overlap with the leading manifold PCs, indicating that the feature is re-encoded along directions of greater manifold variability (**Fig. 5d**). Notably, however, even in network CR, the feature-encoding direction remains broadly distributed across many manifold PCs rather than concentrating along a single dominant PC. In network CR, the greater across-category consistency of the feature encodings in scale and direction also appears to arise predominantly through this within-manifold re-encoding, rather than through an overall rotation of the manifolds. Indeed, overall manifold alignment, as quantified by the RV coefficient between manifold covariance matrices, increases only mildly in network CR relative to network C (**Fig. 5e**). By contrast, network R exhibits much stronger manifold alignment. The source of the weak increase in alignment from network C to CR can be understood by inspecting the matrix of pairwise manifold alignments (**Fig. 5f**). In network C, mild manifold alignment is largely confined to pairs within the same semantic group, as reflected by the block structure of the alignment matrix. In network CR, alignment remains similarly mild in magnitude but is distributed more broadly across manifold pairs, likely reflecting the increased alignment of the shared category-independent feature encodings. In the SI, we also show that manifold centroids—the other key geometric property relevant to classification performance—change only modestly in norm and relative distance in network CR compared with network C.

In sum, in a joint code, the optimization of regression-relevant geometry can be very subtle: feature encodings can be optimized through a reorganization *within* manifolds that only mildly affects manifold shape, centroid geometry, and overall alignment. Moreover, the reencoded features do not become concentrated along a single dominant PC; instead, their encoding directions remain broadly distributed across many PCs. Consequently, substantial changes in regression-relevant geometry can coexist with high global representational similarity: joint and classification-only codes can appear similar under CKA (**Fig. 5g**), consistent with findings reported in a previous study (28). Our results therefore highlight the importance of finer-grained, principled geometric measures—such as those identified by our theory—for characterizing the representational organization of joint codes and revealing how it differs from that of classification-only codes.

### 5. Subsampling effects

We next characterize how typical experimental constraints—namely the subsampling of neural units, and a limited number of categories in the image dataset—affect the estimation of regression performance. We have shown that the most distinctive feature of network CR compared to C is a small relative local–global gap (Δ*E/E*_loc_ ≪ 1); we therefore focus on this quantity here, leaving an analysis of the local error *E*_loc_ to the SI.

We first examine the effect of subsampling the number *P* of categories in our image dataset (**Fig. 6a**). In network C, the relative gap Δ*E/E*_loc_ is strongly underestimated when regressing from few categories (blue curve), potentially making a classification-only population code appear consistent with a joint code. We extended our theory (see Materials and Methods and SI) to capture this finite-*P* behavior (green curve), which can be intuitively understood as a form of overfitting. Crucially, our theory allows one to avoid mistaken conclusions due to this finite-*P* overfitting, as it allows us to reliably extrapolate the local-global gap to the *P* → ∞ limit (red curve). In particular, the manifold-geometric measures entering the theory can be estimated robustly from as few as *P* = 8 categories (the same number of stimulus categories used, for example, in the macaque recordings (26, 27)) yielding a predicted local–global gap consistent with that obtained using all *P* = 265 categories. This can be seen in the relative flatness of the red curve as the number of categories used to estimate the manifold-geometric measures varies from *P* = 8 to *P* = 265.

**Fig. 6.**
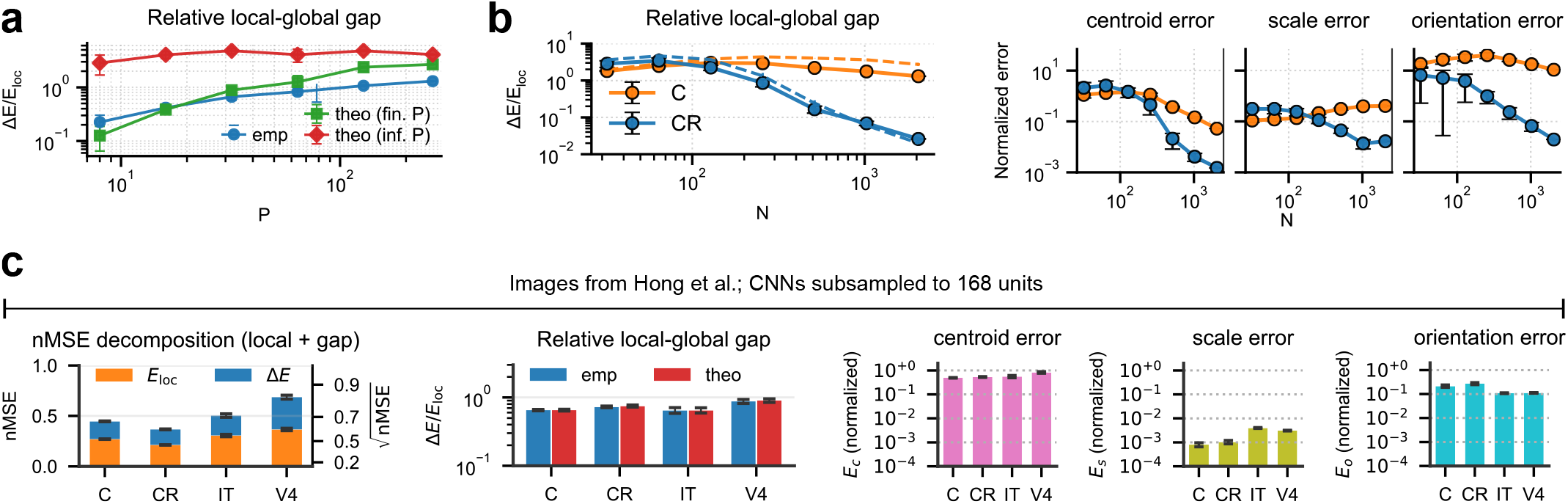
Subsampling effects. (**a**) Relative gap Δ*E/E*_loc_ for regressing the bounding-box feature *L*_*h*_ from network C, as a function of the number *P* of randomly subsampled categories (out of the 265 in our dataset). Blue dots: measured gap; green squares: theoretical prediction for finite *P* ; red diamonds: theoretical prediction in the *P* → ∞ limit, with the manifold-geometric measures entering the theory estimated using only the corresponding *P* subsampled categories; Error bars denote standard deviation across 4 independent category subsamples. (**b**) Δ*E/E*_loc_ (left) and associated error sources (right) for regressing the bounding-box feature *L*_*h*_ from networks C (orange) and CR (blue) on our dataset, as a function of the number *N* of randomly subsampled units (out of 2048 feature-layer units). Dashed lines in the left panel show the theoretical gap prediction. (**c**) Analysis of the regression performance of feature *L*_*h*_, using the images and corresponding neural recordings found in (27). We report the analysis of regression from the IT and V4 units recorded in (27), as well as 168 randomly subsampled units from the feature layer of networks C and CR. Error bars denote SEM across cross-validation splits and, where applicable, random unit subsamples.

Second, we examine the effect of neural-unit subsampling (**Fig. 6b**). For sufficiently small number of neural units, subsampling becomes particularly destructive: in network CR, Δ*E/E*_loc_ increases to the same level as in network C, rendering the two networks indistinguishable by this key metric. Unfortunately, none of the three theoretical error sources remains discriminative under this collapse: all converge to similar values in networks C and CR, with the orientation error *E*_o_ and centroid error *E*_c_ becoming the dominant error contributions. We find a similar phenomenology when analyzing the images and corresponding neural recordings in (27) (**Fig. 6c**): IT, V4, and networks C and CR subsampled to 168 units—the number recorded in IT—all show comparable values of Δ*E/E*_loc_, as well as comparable contributions from each of the three error sources, with *E*_o_ and *E*_c_ emerging as the dominant terms. The similar phenomenology observed in the neural recordings (**Fig. 6c**) and in our controlled CNN analysis (**Fig. 6b**) suggests that this collapse of the local-global gap may, also in the neural data, be largely driven by the limited number of recorded units, and that future experiments may benefit from larger-scale recordings.

## Discussion

In this work, we have identified the representational principles underlying joint codes for stimulus category and continuous category-independent features, and provided a theoretically grounded framework for characterizing such joint codes in terms of representational-geometry measures. By deriving a theory of regression of category-independent features from category manifolds, we have extended existing theories of manifold classification to the missing task of regression, allowing joint codes supporting both classification and regression to be studied within a common framework. Crucially, this common framework is possible because our regression theory is defined over the same category manifolds that underlie previous theories of manifold classification, unlike a recent theory of manifold regression (37), in which manifolds are defined by discretized values of the regressed feature rather than by category.

Our theory is general and can, in principle, be applied to representations generated in any modality, under any training scheme, and by any network architecture, whether biological or artificial. Beyond vision, the framework could be applied to language representations, for example to investigate how the geometry of feature encoding relates to steering vectors (38). Here, we used supervised learning to provide a controlled setting in which joint and classification-only codes could be directly contrasted. A broad representation-learning literature has instead asked how latent factors emerge or become identifiable under unsupervised learning (39–43). Our framework could complement this work by providing principled measures to characterize how such factors are geometrically organized in the representations for linear access. Within vision, it could also be applied to other architectures, such as modern object detectors that predict both object category and bounding-box parameters from the same feature space (13), or vision transformers that learn general-purpose representations (3). Here, we focused on a CNN architecture as a first testbed for our theory because of the direct relevance of CNNs to the hypothesis of a joint code in the macaque ventral stream (26, 28), one of the empirical motivations for this study.

A key piece of evidence for a joint code in macaque IT has been the observation that linear readout performance for both object category and category-independent features improves along the ventral stream, although regression performance remained limited even in IT, potentially because of experimental constraints (26). However, while such improvement along the visual hierarchy is consistent with a joint code, it is not by itself diagnostic of one: regression performance on category-independent features can also improve across the layers of CNNs trained exclusively for object classification, an effect that we reproduce here. Our theory makes a crucial distinction between two contributions through which this improvement in regression performance can arise: a reduction in local error, which quantifies the quality of linear feature encoding within each category manifold, and a reduction in the local–global error gap, which quantifies how consistently these encodings are organized across manifolds to support accurate readout by a single category-independent regressor. Across the visual hierarchy, we find that CNNs optimized exclusively for object classification reduce only the local error, whereas a reduction in the local–global error gap is a prerogative of the CNNs explicitly optimized for regression. Curiously, we find that a small local–global gap can emerge even when the regression objective is category-specific, without the additional constraint of a shared category-independent regressor. In sum, we propose a small relative local–global error gap as a distinctive signature of joint coding.

We have characterized how estimation of this distinctive local-global gap, and in general of regression error, is affected by common experimental constraints, such as a limited number of stimulus categories and neural population subsampling.

A limited number of stimulus categories can lead to a large underestimation of the local-global gap. Crucially, our theory allows this limitation to be overcome through a principled extrapolation to the many-category limit. This extrapolation is enabled by two features of our theory. First, the theory predicts the local–global gap in terms of manifold-geometric measures whose estimation we find to remain robust even with as few as eight categories. Second, our theory provides a theoretical account of the local–global gap in the regime most relevant for category-independent regression, namely the limit of infinitely many categories, while also providing finite-category corrections that connect empirical measurements to this limit. In contrast, previous work has studied the local–global gap only empirically, and only for pairs of category manifolds (44).

Regarding neural-unit subsampling, in our CNNs we find that it causes the local–global gap of joint codes to approach that of classification-only codes, erasing the distinction between them under this key metric. Unfortunately, none of the manifold-geometric measures underlying the local-global gap remains discriminative under subsampling: all progressively converge between joint and classification-only codes. We observe a suggestively similar phenomenology in the macaque recordings (26, 27), consistent with the possibility that limited population sampling may likewise obscure joint-code signatures in IT. This suggests that larger-scale neural recordings may be required for future experiments to test the joint-code hypothesis more decisively. Future experiments may also benefit from our novel image-generation process, which, compared with previous approaches that place objects in random environments (27, 28), produces more photorealistic images with contextually relevant environments, albeit at the cost of losing explicit control over object pose.

Another open problem in characterizing joint codes has been the observation that, in CNNs, these representations can appear virtually indistinguishable from classification-only representations under common representational-similarity measures such as CKA (28). Our theory identifies principled measures of manifold geometry that can instead characterize joint codes and, moreover, help explain this previously observed high similarity under CKA. We find that, because regression-relevant geometry pertains primarily to specific directions within each manifold—namely, the encoding directions of the regressed features—it can be optimized *within* manifolds while largely preserving coarser measures of manifold geometry relevant for classification, such as manifold radius and dimensionality. Thus, substantial changes in regression-relevant geometry can coexist with high global representational similarity.

An open question is whether this picture persists when representations must encode many more category-independent variables. A limitation of our theory is that it considers regression of one feature at a time, neglecting potential effects arising from the concurrent organized encoding of other features. To assess this limitation, in the SI we replicated our analysis on CNNs trained to regress up to eight features: the four bounding-box parameters together with four additional features—bounding-box area, mean luminance, local contrast, and color saturation. We find that all of our main results continue to hold with up to these eight jointly encoded features. Because the expanded feature set includes nonspatial image statistics, these findings also serve to illustrate the generality of our theory, showing that its agreement with empirical results extends beyond spatial features such as object position and size. A challenge for future work will be to devise large-scale image datasets annotated with many more continuous features, allowing substantially richer joint codes to be systematically investigated and the limits of our theory to be tested further.

## Materials and Methods

### Linear decoding

To measure linear-readout performance for object classification and category-independent regression, we used the following procedures. We decoded object category using linear one-vs-rest ridge classifiers. For each category, we fit a separate readout with binary labels *y* = 1 for images in that category and *y* = 0 otherwise, and thresholded readout scores at 0.5. Performance was measured by balanced accuracy, 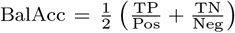, where TP and TN are the numbers of true positives and negatives and Pos and Neg the corresponding numbers of positive and negative test samples. We report balanced accuracy averaged across categories. For global regression, we fit a linear ridge regressor to each scalar bounding-box feature using data pooled across categories. Performance was measured by normalized MSE (nMSE), obtained by averaging test MSE across outer folds and dividing by the variance of the regressed feature. For local regression, we jointly fit category-specific regressors using a shared regularizer across categories rather than a separate *L*_2_ regularizer for each category, and computed *E*_loc_ as the average test MSE across categories. This yielded *E*_loc_ values statistically compatible with those obtained from independent per-category ridge regressors, while reducing noise from regularization selection and providing more stable estimates of the geometric quantities entering our theory (see SI for details). Regularization strengths were selected by nested k-fold cross-validation, with 10 outer train–test folds and 10 inner folds for regularization selection. For across-layer analyses, the same decoding procedures were applied at every layer. Pixels and intermediate activations were flattened across channels and spatial positions and projected to 2048 dimensions using a fast Johnson–Lindenstrauss transform. The 2048-dimensional feature layer was used directly. Layer labels follow the torchvision ResNet-50 module hierarchy: “1” and “2” denote the outputs of layer1 and layer2, while “3.x” and “4.x” denote the outputs of individual residual blocks within layer3 and layer4.

### Dataset generation

Our dataset consists of photorealistic images of a single main object placed in a contextually relevant scene. Each image is labeled with the object identity (one of 265 categories) and the parameters of its bounding box: center coordinates *C*_*h*_, *C*_*v*_ and side lengths *L*_*h*_, *L*_*v*_ along the horizontal (h) and vertical (v) axes. The 265 categories are a subset of the 365-category Objects365 taxonomy, selected because they could be generated reliably with our pipeline. Each category contains 10,000 training images used to train the CNNs and a disjoint set of 10,000 test images, unseen during CNN training, used to train and evaluate linear decoders. Images were generated using the following procedure (**Fig**. 7). First, an image of an object from the desired category (e.g., “wild bird”) was generated from a text prompt using Stable Diffusion XL (34, 45), with categories drawn from the Objects365 taxonomy (46). The object’s bounding box was then estimated using CerberusDet (47), an Objects365-trained detector. The image was isotropically rescaled and pasted onto a white canvas so that the detected box matched a target box whose parameters (*C*_*h*_, *C*_*v*_, *L*_*h*_, *L*_*v*_) were drawn uniformly, subject to preserving aspect ratio and remaining within the canvas (see also the SI). The resulting canvas was used to condition a second generation step with a Stable Diffusion 1.5 inpainting model (33, 48, 49), yielding the final image. This procedure produces images with a controlled distribution of bounding boxes. Further details on the generation procedure and additional example images are provided in the SI.

**Fig. 7.**
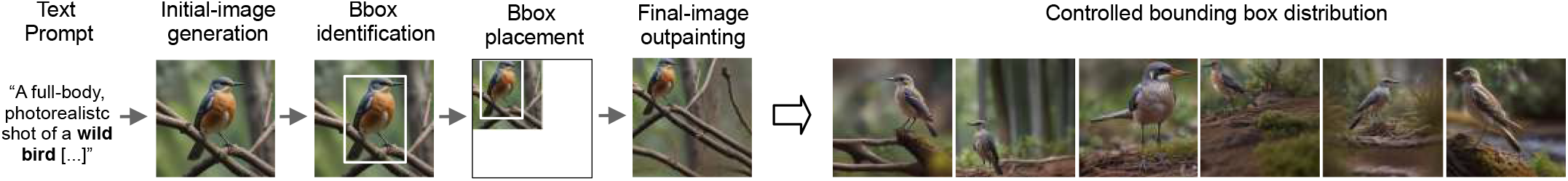
Scheme of the generation process for an image in our dataset.

### CNN architecture and training

All CNNs used an ImageNet-pretrained ResNet-50, with the original classification head removed and the final 2048-dimensional feature layer used as the shared representation. We attached a 265-class linear classification head and/or four linear regression heads predicting (*C*_*h*_, *C*_*v*_, *L*_*h*_, *L*_*v*_), initialized with Xavier-uniform weights and zero biases. We trained three main variants: C (classification only), R (regression only), and CR (classification and regression). Images were resized to 224 × 224 and normalized using standard ImageNet channel statistics; bounding-box parameters were standardized using the training-set mean and standard deviation. Models were trained on shuffled mini-batches from 8,000 images per category, with 2,000 images per category held out for validation. We used stochastic gradient descent (batch size 208; learning rate 0.02; momentum 0.9; weight decay 10^−4^; cosine annealing over 40 epochs), Smooth L1 loss for each regression coordinate, and cross-entropy loss for classification. The total loss summed the applicable regression and classification terms, omitting those corresponding to absent heads. Models were trained until validation performance plateaued.

### Theory definitions and results

#### Definitions of regression errors

Our theory is derived for the training regression error (when applied to real data, however, we test its predictions against the corresponding test error; see ‘Fitting theory to data’). Here we define the training regression errors entering the theory. Consider a set *C* consisting of *P* categories. Call *x*^*µ*^ ∈ R^*N*^ the neural representation of an image (or, more generally, an input stimulus) in the *µ*-th category, where *N* is the number of neural units. Let *y*^*µ*^ be a continuous label associated with the image (e.g. one of the bounding-box parameters). Without loss of generality, we consider globally centered data and labels, that is ⟨⟨*x*^*µ*^⟩_*µ*_⟩_*C*_ = 0 and ⟨⟨*y*^*µ*^⟩_*µ*_⟩_*C*_ = 0, where ⟨…⟩_*µ*_ denotes averaging over all images within the *µ*-th category, and ⟨… ⟩ _*C*_ = Avg_*µ*_ […] denotes averaging over categories.

Define the global regression MSE as

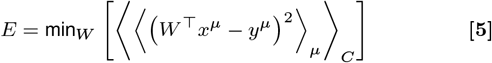

where *W* ∈ ℝ^*N*^ is the global regression vector. We decompose the global regression MSE as *E* = *E*_loc_ + Δ*E*. Here, *E*_loc_ = ⟨*E*^*µ*^⟩_*C*_ and *E*^*µ*^ is the local regression MSE for the *µ*-th category, defined as

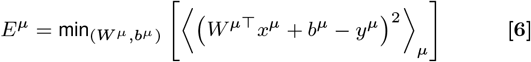

where *W*^*µ*^ ∈ ℝ^N^, *b*^*µ*^ ∈ ℝ are the local regression vector and bias for category *µ*. It can be shown (see SI) that Δ*E* = *E* − *E*_loc_ is given by

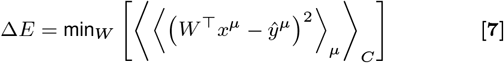

where 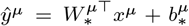 is the label predicted by the local regressor for category *µ*, with the local regressor defined as 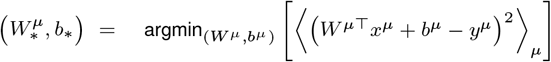. In other words, the local-global error gap is itself a global regression error in which the target is the locally linearized label *ŷ*^*µ*^ rather than the original label *y*^*µ*^.

#### Generative model

We derive Δ*E* for the following generative model of manifolds. We first consider the case in which each manifold and associated labels are locally centered, that is ⟨*x*^*µ*^⟩_*µ*_ = 0 and ⟨*y*^*µ*^⟩_*µ*_ = 0, ∀*µ*. Without loss of generality, we parameterize a point in the *µ*-th manifold as

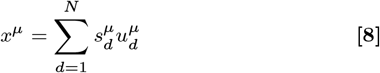

Where 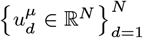 is an orthonormal basis and 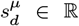 is the coordinate of the point along direction *d*. We choose 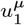 to correspond to the feature-encoding direction, that is 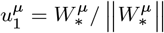. We choose the remaining directions 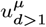 as the principal-component directions of the manifold in the subspace orthogonal to 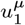, such that

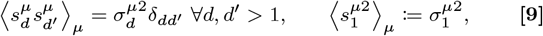

while there can be correlations between 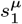 and 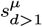. Note that local centering of the manifolds implies 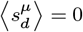. This choice of parameterization can always be made without loss of generality. We then make the following assumptions on the across-manifold statistics:

1. The feature-encoding directions 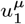 share a common mean direction and isotropic second-order fluctuations across categories

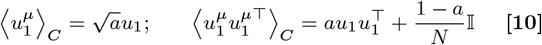

where *a* ∈ [0, 1] is the alignment strength, and *u*_1_ ∈ R^*N*^ is a unit-norm vector representing the common mean direction.
2. All manifold directions with *d >* 1 are isotropically distributed up to their second order statistics, conditionally on being orthogonal to 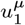 (as by their definition as the elements of an orthonormal basis). Concretely, 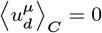 and

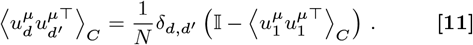
3. Aside from the above correlations, all manifold parameters are otherwise uncorrelated.

Note that no assumptions are made nor need to be made about higher order statistics of 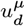, since regression by MSE minimization is a quadratic problem and therefore depends only on statistics up to second order.

#### Theory results

Our theoretical results are derived in the asymptotic regime *P*→ ∞, with *N* large but finite. This is the regime most relevant to category-independent regression, because a category-independent regressor should apply to images from arbitrary categories drawn from the same statistical ensemble. This differs from the asymptotic regime relevant to manifold capacity (29), in which *P, N* → ∞ at fixed category load *α* = *P/N*. We also derive an extension of our theory to this regime; see “Theory for finite number of categories” below. Under the generative model described above, and neglecting higher order corrections in 1*/N*, we obtain (see SI for the derivation).

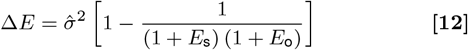

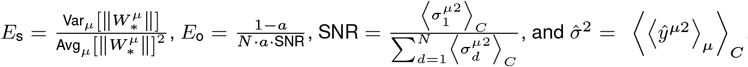. In the case in which manifolds and labels are not locally centered, we quantify the contribution of category centroids through the centroid error *E*_c_, defined as

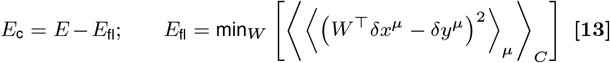

with *δx*^*µ*^ = *x*^*µ*^ − ⟨*x*^*µ*^⟩_*µ*_ and *δy*^*µ*^ = *y*^*µ*^ − ⟨*y*^*µ*^⟩_*µ*_ denoting the locally centered representations and labels. That is, *E*_c_ measures the additional regression error induced by the presence of category-dependent centroids. The remainder of the theory is then derived exactly as above, with *E*_fl_ taking the role of *E*.

#### Theory for finite number of categories

We also derive the local-global gap Δ*E* for finite number of categories. Formally, the result is derived in the regime of finite category density, by studying the thermodynamic limit *P, N* → ∞ at fixed 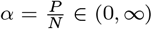. The result for infinitely many categories presented above (*P*→ ∞ with *N* large but finite) is recovered as the limiting case *α* → ∞. The expression for generic *α* is lengthy and is provided in the SI, alongside its derivation.

#### Fitting theory to data

The theory derived above applies to the *training* regression error. For a validation of the theory in this setting, using the generative model described above, see the SI. In the main text, instead, we test its predictions against the *test* regression error measured from the empirical data, and find good agreement in this case as well. This agreement assumes that the number of images is sufficiently large and that the regularization of the fitted regressor is appropriate, so that the train-test generalization gap is small and the theoretical prediction remains informative for the test error. For the purpose of estimating the theory parameters from data, the local regressors are taken to be the regularized empirical solutions used in practice, as determined by the procedure described in the ‘Linear Decoding’ section of the Methods, rather than the unregularized minimizers of the local MSE. Accordingly, all quantities entering the theory that depend on the local regressors are evaluated using these regularized estimates.

## Data, Materials, and Software Availability

The code to reproduce the results of this study is publicly available at https://github.com/tiberilor/manifolds-joint-code.

## ACKNOWLEDGMENTS

We thank Xu Pan and David Klindt for helpful discussions. This research is supported by the Swartz Foundation, the Gatsby Charitable Foundation, the Kempner Institute for the Study of Natural and Artificial Intelligence at Harvard University, the NSF grant PHY-2309135 to the Kavli Institute for Theoretical Physics and Office of Naval Research grant No. N0014-23-1-2051.

