## Appendix SI for "Neural representational geometry of a joint code for stimulus category and category-independent features"

#### This PDF file includes:

Supporting text
Figs. S1 to S28
SI References

### Supporting Information Text

### 1. Supplementary results

**A. Linear decoding performance in macaque recordings.** We have applied the linear decoding framework presented in the main text to the publicly available dataset of macaque recordings (1) studied in (2), as well as to an ImageNet-pretrained ResNet-50 CNN. We reproduce their qualitative results (**Fig. S1a**): decoding performance of both category and category-independent features improves along the visual hierarchy. However, we also find that absolute regression performance remains limited (**Fig. S1b**): in macaque IT, as well as in the CNN for most features, root nMSE remains above 0.5, meaning that the root-mean-square prediction error is still larger than half the across-image standard deviation of the decoded feature.

We note that this last point may be less apparent in the original presentation of (2), where regression performance was reported using the Pearson correlation coefficient. Here we instead report $\sqrt{\text{nMSE}}$, which we find more directly interpretable, as it measures the typical prediction error (root-mean-squared error) in units of the standard deviation of the decoded feature. In particular, because Pearson correlation is a nonlinear function of nMSE, it tends to compress even appreciable differences in nMSE into a narrow interval near 1. Indeed, for ordinary least-squares (no ridge) evaluated on the training set, Pearson correlation is related to nMSE by $\text{Pears} = \sqrt{1 - \text{nMSE}}$. For ridge regression on held-out data, this relation is only approximate but still useful as an intuition. We can see that Pearson correlation compresses a broad range of $\sqrt{\text{nMSE}}$ values into a narrow range near 1. For example, $\sqrt{\text{nMSE}} \in [0, 0.5]$—from perfect prediction to errors on the order of half a standard deviation—maps to $\text{Pears} \in [0.87, 1]$, making differences in regression accuracy harder to visualize and interpret.

**Fig. S1b** additionally reports regression performance using the Pearson correlation coefficient, alongside nMSE, to facilitate a more direct comparison with (2). The Pearson values we obtain are comparable to those reported in that work (cf. Supplementary Fig. 3 and Supplementary Fig. 9 of (2)). This comparison is not exact, because the original analysis was performed on the full private dataset, whereas we here use the publicly available subset. Relative to the public dataset, the full dataset contained 320 additional images per category, as well as 98 additional IT units recorded from a third macaque. Even so, the Pearson-based regression performance reported here remains of similar magnitude to that reported in (2). In particular, the IT Pearson values in (2) are still sufficiently below 1 to be consistent with a root nMSE well above 0.5, matching the conclusions we draw here from the public data.

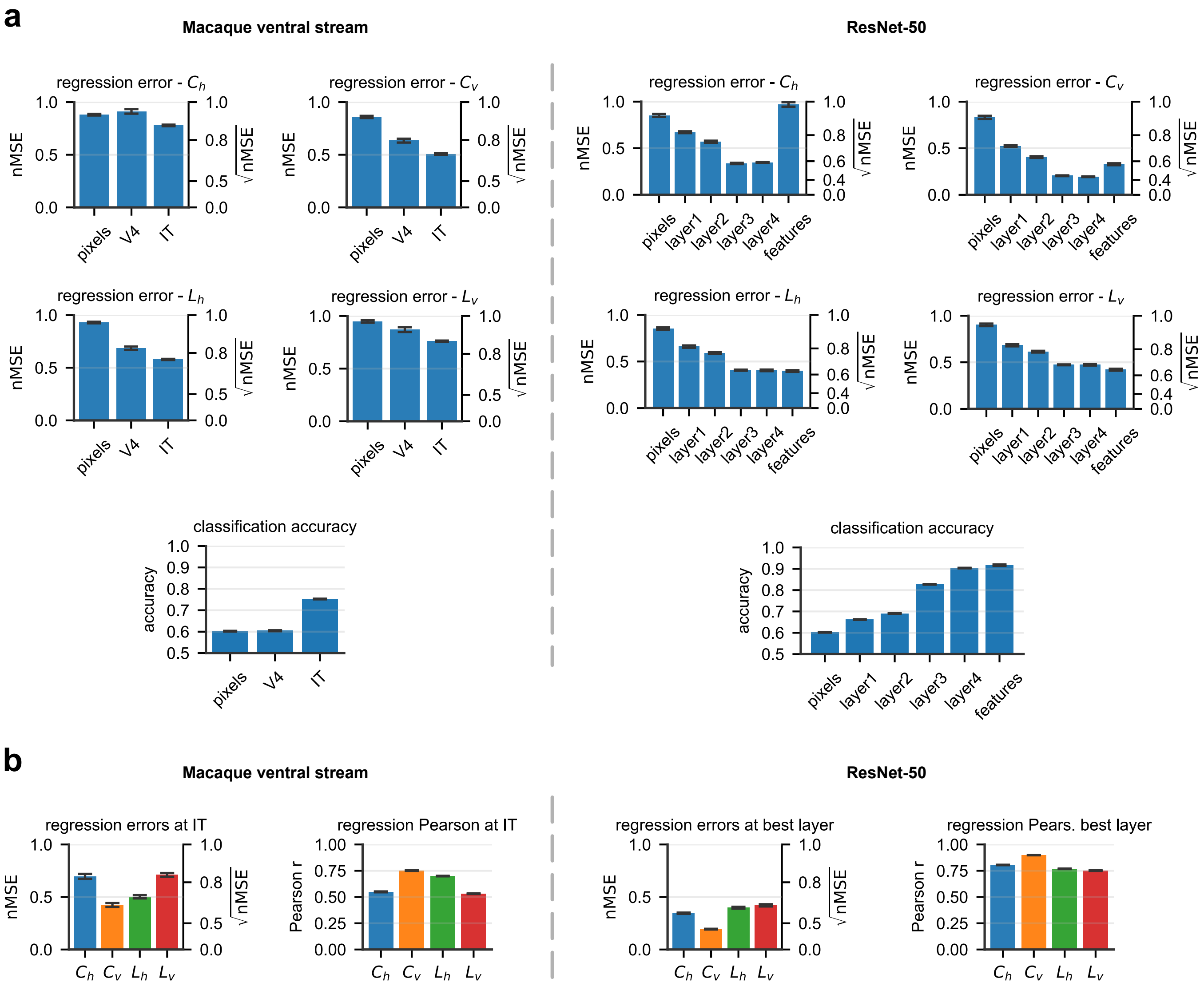

**Fig. S1.** Linear decoding performance for macaque neural recordings and ResNet-50 activations. (**a**) Linear decoding performance along the visual hierarchy, for macaque neural recordings (left column) and ResNet-50 activations (right column). For a fair comparison across regions, the 168 IT units are randomly subsampled to match the 88 units in V4. Similarly, units in layers 1–4 of ResNet-50, as well as the pixel layer, are randomly projected to match the 2048 units in the final feature layer. (**b**) The regression performance at IT using all 168 units (left column), and error at the best-performing layer of ResNet-50 (right column). We report performance both in terms of the nMSE (left) and the Pearson coefficient (right). Error bars indicate the standard error of the mean (SEM) across cross-validation splits and, where applicable, across random subsamples of units.

**B. Network optimized for classification and category-specific regression (network CRloc).** Here we show that optimization for regression alone is sufficient to substantially reduce the local-global gap, even in the absence of the additional constraint imposed by a shared readout across categories. Indeed, we find that $\Delta E$ is largely reduced not only in network CR, which is optimized for *category-independent* regression and classification, but also in a network that we term CRloc, which we optimized for *category-specific* regression and classification (**Fig. S2a-b**). In CRloc, category-specific regression is implemented by attaching multiple regression heads to the feature layer—one for each category—rather than a single head shared across categories. We observed that, in network CRloc, $\Delta E$ is largely reduced compared to network C, down to values only slightly higher than those for network CR. The same observation holds for the three sources of error $E_{\mathrm{c}}$, $E_{\mathrm{s}}$, $E_{\mathrm{o}}$ (**Fig. S2c**), and the two factors contributing to $E_{\mathrm{o}}$, namely alignment $a$ of the feature-encoding directions, and SNR (**Fig. S2d**).

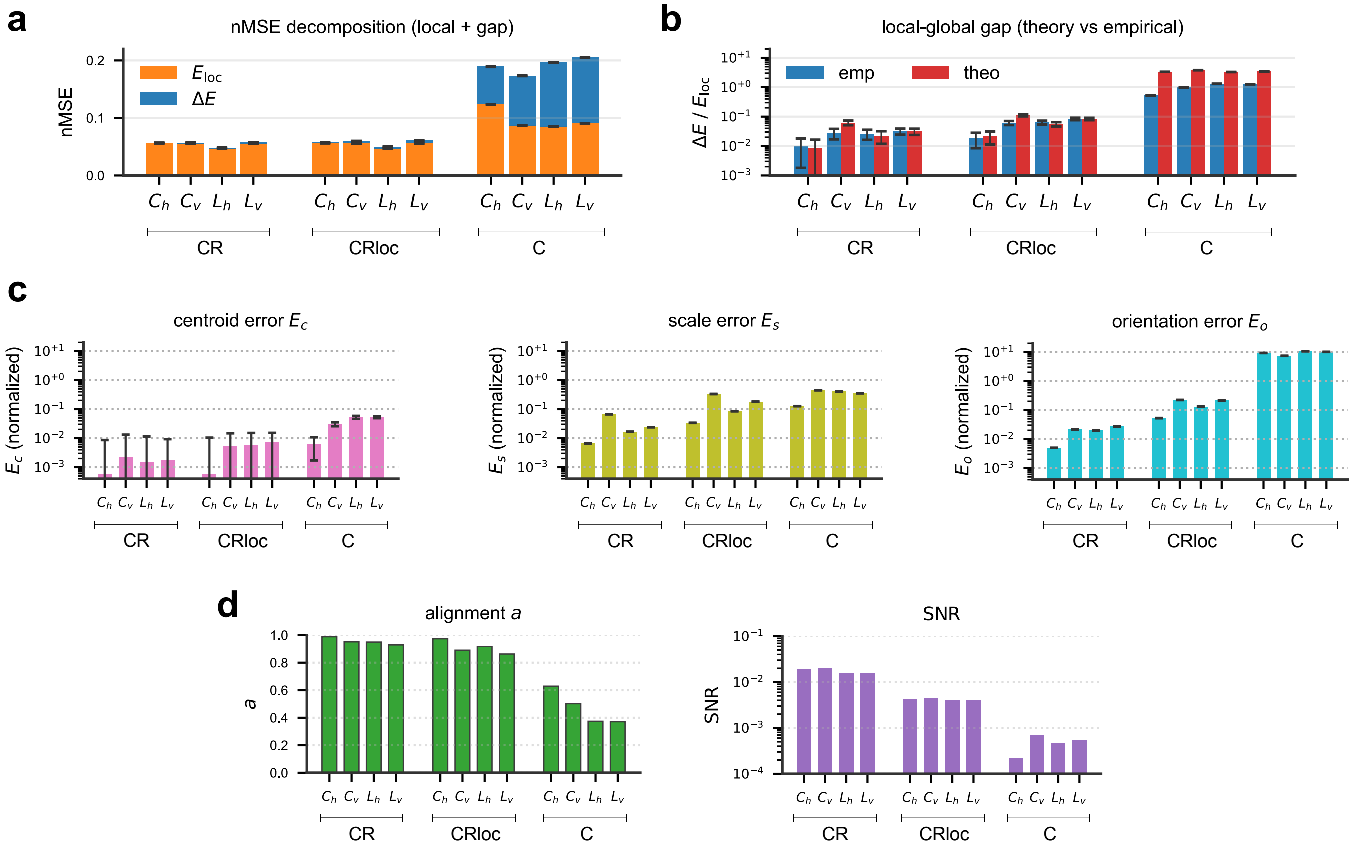

**Fig. S2.** Regression performance in networks CR, CRloc and C. (**a**) Decomposition of the nMSE into local error $E_{\text{loc}}$ (orange) and local–global gap $\Delta E$ (blue) for regression of the four bounding box coordinates. Note that $\Delta E$ is very small and barely visible for networks CR and CRloc. (**b**) Measured relative local-global gap $\Delta E / E_{\text{loc}}$ (blue) compared to our theory prediction (red). (**c**) Decomposition of the relative gap $\Delta E / E_{\text{loc}}$ into the contributions from the centroid error $E_{\text{c}}$ (pink) scale error $E_{\text{s}}$ (olive) and orientation error $E_{\text{o}}$ (cyan). The three errors are normalized as explained in the caption of **Fig. 4**. (**d**) The two elements contributing to the orientation error: SNR, and alignment $a$ of the local feature-encoding directions.

### C. Additional results for section "Manifold geometry reveals optimization strategy".

***C.1. Per-category changes in manifold shapes.*** In the main text, we showed that, on average across categories, manifold shapes are only weakly altered in network CR relative to network C (cf. **Fig. 5b**). Here we examine this effect on a category-by-category basis. For a given category $\mu$, let $\{\lambda_d^\mu\}_{d=1}^N$ and $\{\gamma_d^\mu\}_{d=1}^N$ denote the PC eigenvalues of the $\mu$-th category manifold in networks CR and C, respectively. We quantify the spectral difference between these two manifolds as

$$d^\mu = \mathcal{N}^{-1}\left[\sum_{d=1}^{N}\left(\sqrt{\lambda_d^\mu} - \sqrt{\gamma_d^\mu}\right)^2\right]^{1/2}, \qquad \mathcal{N} = \left[\sum_{d=1}^{N}(\lambda_d^\mu + \gamma_d^\mu)\right]^{1/2}. \qquad [1]$$

Without the normalization factor $\mathcal{N}$, this quantity corresponds to the Wasserstein-2 distance between the covariance matrices of category $\mu$ in networks C and CR, after aligning their principal axes, which is appropriate here since our goal is to compare manifold shapes independent of their orientations. Intuitively, the Wasserstein-2 distance measures how much one distribution must be rearranged to match the other. Here we normalize this distance by the combined root variance of the two manifolds, $\mathcal{N}$, so that the resulting quantity is dimensionless, bounded in the range $d^\mu \in [0, 1]$, and comparable across categories. **Fig. S3** reports the histogram of $d^\mu$ across categories, together with example PC spectra for selected categories in networks C and CR. These results show that manifold shapes undergo only small relative changes between networks C and CR, with $\text{Avg}_\mu[d^\mu] \sim 0.083$.

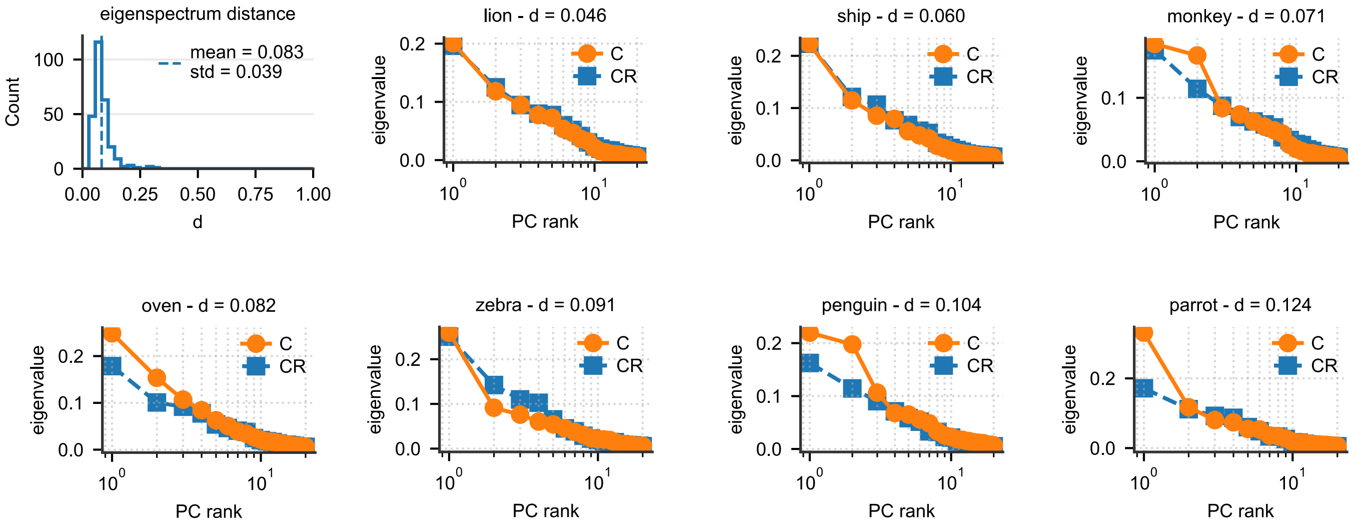

**Fig. S3.** Change in manifold shapes between networks C and CR. Top left: histogram across categories of the spectral distance $d^{\mu}$. Other panels: PC spectra for selected categories in networks C and CR. Categories were sampled randomly, with sampling probability weighted by a Gaussian distribution with mean and standard deviation equal to those of the empirical distribution of $d^{\mu}$. For each category, the PC spectra are normalized by the category total variance, averaged between networks C and CR.

***C.2. Changes in centroid norm and separation.*** For each category manifold $\mu$, we denote by $\bar{x}_\mu \in \mathbb{R}^N$ its centroid. We compare two centroid measures across networks C and CR: the centroid norm $\|\bar{x}_\mu\|$, and the pairwise relative centroid distance $\|\bar{x}_\mu - \bar{x}_\nu\|/\langle\|\bar{x}_\rho\|\rangle_\rho$ between two manifolds $\mu$ and $\nu$. Centroid norms see only modest changes in network CR relative to network C, while centroid distances are virtually unchanged in distribution (**Fig. S4**).

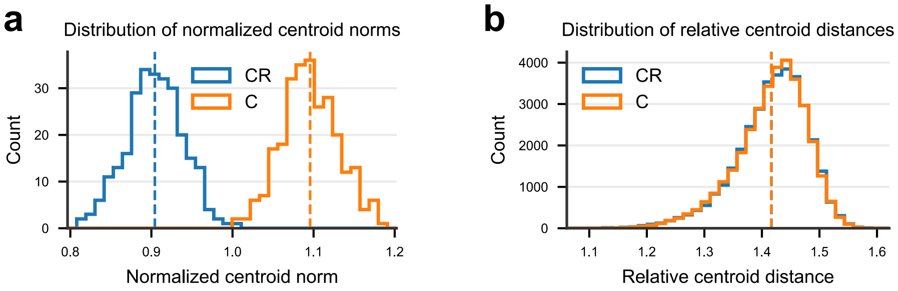

**Fig. S4.** Manifold centroid norm and separation, in networks C (orange) and CR (blue). (a) Histogram of centroid norms $\|\bar{x}_\mu\|$. Centroid norms are normalized by their average across categories and across the two networks. (b) Histogram of relative centroid distances $\|\bar{x}_\mu - \bar{x}_\nu\|/\langle\|\bar{x}_\rho\|\rangle_\rho$. Vertical dashed lines indicate the corresponding averages.

***C.3. Additional information on manifold dimensionality and CKA.*** As shown in the main-text analysis of the geometry optimization strategy (**Fig. 5**), the PC spectrum of network R is dominated by only four components; we reproduce this result here for convenience (**Fig. S5a**, left), together with its dimensionality according to manifold-capacity theory (Fig. **S5a**, center). One might then wonder why the latter is substantially larger for network R than for networks C and CR. This apparent discrepancy arises because the dimensionality defined by manifold-capacity theory need not coincide with conventional covariance-based measures of manifold dimensionality, such as the participation ratio. Denoting by $\lambda_d^\mu$ the covariance eigenvalues of manifold $\mu$, the participation ratio is defined as $D_{\mathrm{PR}}^\mu = (\sum_d \lambda_d^\mu)^2 / \sum_d (\lambda_d^\mu)^2$ and is shown in **Fig. S5a**, right. As expected from this definition, network R has a participation-ratio dimensionality close to four. By contrast, dimensionality in manifold-capacity theory captures geometric properties beyond the second-order statistics summarized by the PC spectrum. In particular, it is defined from the anchor points that constrain linear separability and therefore determine manifold storage capacity (3). Lower anchor dimensionality generally corresponds to greater storage capacity and is therefore preferable for this task. Instead, the opposite is true for participation-ratio dimensionality, which appears in the theory of few-shot learning accuracy (4): higher dimensionality improves generalization from a limited number of examples. Thus, the lower anchor dimensionality and higher participation-ratio dimensionality of networks C and CR are both favorable, but for two distinct proxy measures of classification performance: storing many familiar categories and generalizing from a few examples of novel categories, respectively.

In the main text, we compared the representations of networks C, CR, and R using centered kernel alignment (CKA). Following (5), we computed CKA over the full set of representations elicited by all images from all categories. We reproduce this result here for convenience (**Fig. S5b**, left). Networks C and CR are nearly indistinguishable under this metric, with a CKA value close to one. This high similarity is consistent with our finding that joint optimization in network CR modifies regression-relevant manifold geometry subtly, largely preserving the manifold shapes and centroid geometry that support classification. To better understand this, we also computed CKA separately for each category and then averaged the resulting values across categories (**Fig. S5b**, right). Specifically, for each category, we centered its representations and computed CKA between the corresponding manifolds in two networks. Under this per-manifold analysis, network CR becomes more distinguishable from network C and is approximately equally similar to networks C and R. Together, these results indicate that the high full-data CKA between networks C and CR is driven primarily by their nearly unchanged arrangement of manifold centroids. The relatively mild differences in manifold shape between the two networks have little effect on full-data CKA compared with this dominant centroid similarity. At the same time, per-manifold CKA shows that network CR retains substantial similarity to both networks C and R, consistent with regression optimization modifying within-manifold geometry without completely restructuring it. Nevertheless, although per-manifold CKA reveals the intermediate character of network CR, it remains less informative for identifying joint codes than the quantities identified by our theory such as the local-global error gap and its underlying geometric components. CKA requires one or more reference representations against which the representation of interest can be compared, in this case networks C and R, and it has no direct quantitative relationship to regression performance. By contrast, the measures identified by our theory provide reference-independent signatures of joint coding that are directly linked to performance. In particular, a joint code is expected to exhibit a small relative local-global gap, independently of any comparison with another network.

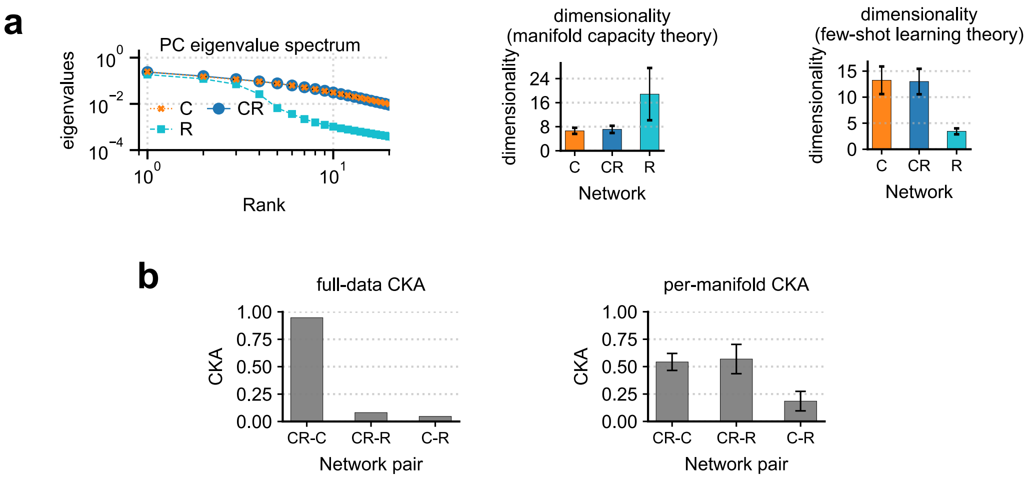

**Fig. S5.** Additional measures of manifold dimensionality and representational similarity. (a) Left: category-averaged PC spectra, normalized by total variance (mean ± SEM). Center: dimensionality from manifold-capacity theory. Right: participation-ratio dimensionality. Bars show mean ± SD across categories. (b) Debiased linear CKA between network pairs, computed using either the full dataset (left) or each centered category manifold separately and then averaged across categories (right; mean ± SD). Colors denote networks C (orange), CR (blue), and R (cyan).

***C.4. Alignment with neural data.*** **Fig. S6** reports the neural-data alignment of networks C, CR, and R, together with an ImageNet-pretrained ResNet-50, as evaluated using the public Brain-Score benchmarks (6). Networks C and CR exhibit highly similar alignment scores across layers and brain regions, consistent with their similarity under coarse manifold-geometry measures (manifold shapes, centroid norms and separation, and manifold alignment) and CKA. Importantly, however, network R also achieves broadly similar Brain-Score values across many regions and layers, despite diverging from C and CR under manifold-geometry measures and CKA, particularly in the final feature layers. Thus, similar neural-alignment scores cannot be attributed uniquely to similarity under these representational measures. These results admit two non-mutually exclusive interpretations. First, Brain-Score and our geometric analyses may capture partly distinct aspects of the representations. Second, the available neural recordings and associated Brain-Score benchmarks may lack sufficient resolving power to distinguish between models with meaningfully different representational geometries. The latter interpretation is consistent with our conclusion in the main text that the number of recorded units may be insufficient to reliably distinguish a genuine joint code from a representation optimized exclusively for classification.

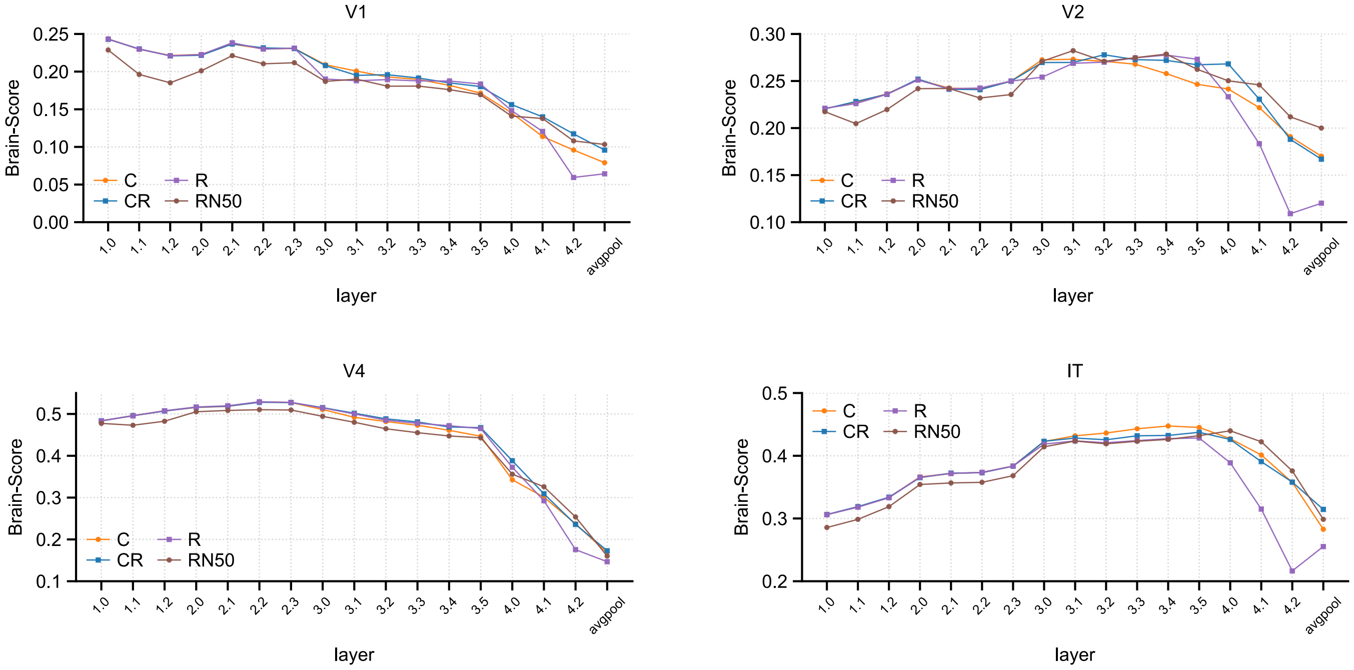

**Fig. S6.** Neural-data alignment on public Brain-Score ventral-stream benchmarks. For each layer of networks C, CR, R, and an ImageNet-pretrained ResNet-50, we report the Brain-Score alignment to four publicly available neural benchmarks (1, 7): FreemanZiemba2013.V1-pls (V1), FreemanZiemba2013.V2-pls (V2), MajajHong2015.V4-pls (V4), and MajajHong2015.IT-pls (IT).

**D. Across-layer behavior of regression-relevant geometry measures.** Here we report the across-layer behavior of the manifold-geometry measures underlying the local-global error gap as identified by our theory (**Fig. S7**). For convenience, we also report the quantities already reported in the main text (**Fig. S7a**): classification accuracy, global regression error, local error and local-global error gap with theory prediction. Additionally, we report the manifold-geometry measures contributing to the local–global gap—namely the centroid, scale, and orientation errors (**Fig. S7b**). In both networks C and CR, the centroid error is only relevant in early layers, while in later layers it becomes negligible compared to the other sources of error. The scale error remains approximately constant in network C, whereas network CR tends to reduce it in the final layers. In contrast, the orientation error increases sharply in network C. This behavior can be understood by inspecting the two components of the orientation error: SNR and alignment $a$ (**Fig. S7c**). In network C, alignment $a$ remains approximately constant, while SNR decreases drastically. This is consistent with the classification-only objective of network C, which might incentivize suppression of class-irrelevant variability such as bounding-box coordinates, thereby reducing their SNR. By contrast, network CR appears to counteract this increase in orientation error by improving both SNR and alignment $a$ across layers.

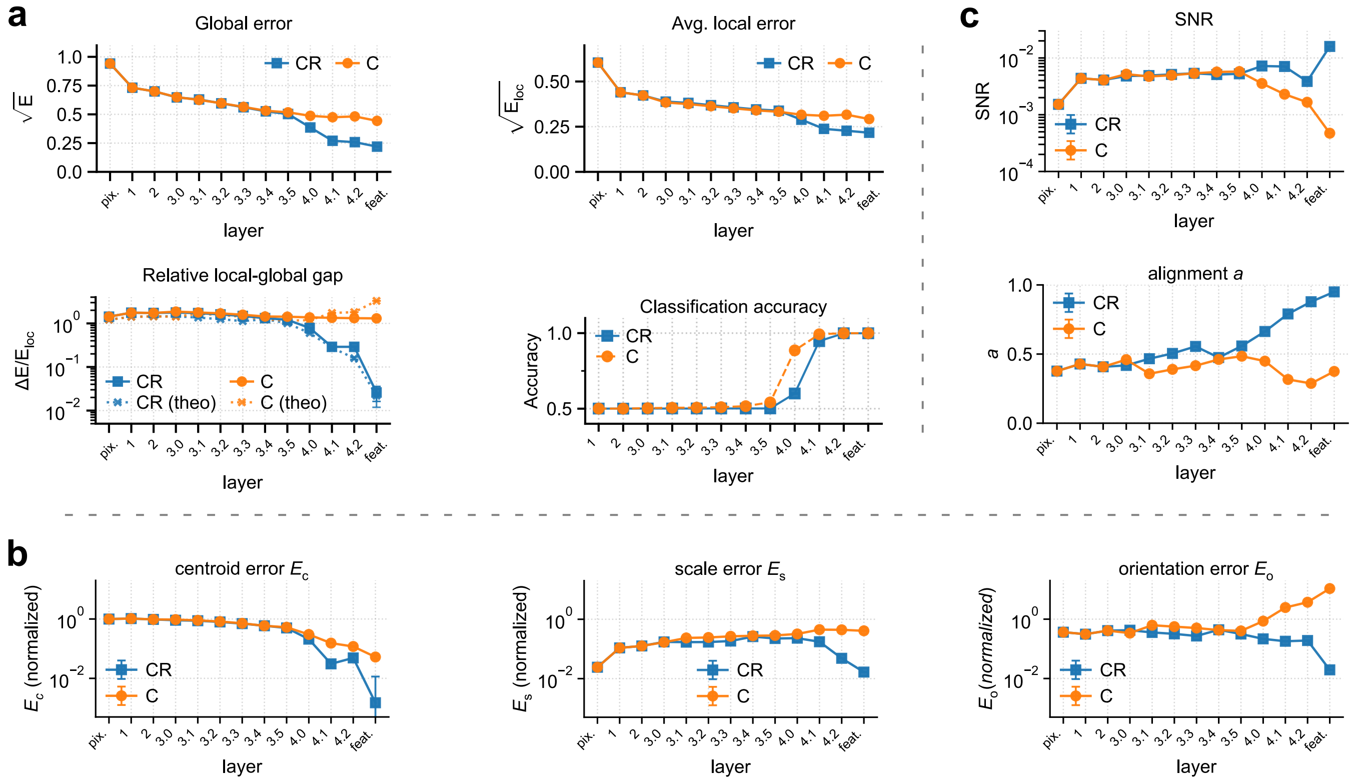

**Fig. S7.** Decoding performance along the visual hierarchy. (**a**) Regression performance for the bounding-box parameter $L_\mathrm{h}$ and classification performance, using linear readouts from progressively deeper layers of networks C (orange circles, solid line) and CR (blue squares, solid line). At each layer, linear readouts were trained on activations randomly projected to 2048 dimensions, to match the dimensionality of the final feature layer. For regression, we report the global regression error, as well as its decomposition into local error $E_\mathrm{loc}$ and relative local-global gap $\Delta E/E_\mathrm{loc}$. For the gap, we also report our theory prediction (x markers, dotted lines). Note that global and local error are reported as root nMSE. (**b**) The three contributions to the relative local-global gap, as identified by our theory: centroid error $E_\mathrm{c}$, orientation error $E_\mathrm{o}$, scale error $E_\mathrm{s}$. These quantities are normalized in the same way as explained in **Fig. 4**. (**c**) The two elements contributing to the orientation error: SNR, and alignment $a$ of the local feature-encoding directions.

**E. Additional category-independent features (bounding-box area, mean luminance, local contrast, and color saturation).** To test whether our main results continue to hold when a network is required to encode more than four bounding-box features alongside object category, we considered eight jointly encoded features: the four bounding-box coordinates together with four additional features, namely bounding-box area, mean luminance, local contrast, and color saturation. These additional features also allow us to test the generality of our theory and establish that its agreement with empirical results does not depend on the particular spatial bounding-box features considered in the main text. We find that our main theoretical and empirical results continue to hold when as many as eight features are jointly encoded.

***E.1. Features definition.*** Bounding-box area was defined as $A = L_h L_v$. The mean luminance, local contrast, and color saturation were computed from the same $224 \times 224$ RGB image provided to the CNN, before ImageNet normalization. Let $p$ index the $N = 224^2$ pixels, and let $R_p$, $G_p$, and $B_p$ denote the red, green, and blue values at pixel $p$, scaled to $[0, 1]$. For any pixel-wise quantity $f_p$, we denote its spatial average by $\langle f_p \rangle_p = \frac{1}{N} \sum_{p=1}^{N} f_p$. Mean luminance was defined as $\overline{Y} = \langle Y_p \rangle_p$, where $Y_p$ is the pixel-wise luminance $Y_p = 0.2126 R_p + 0.7152 G_p + 0.0722 B_p$,. Local contrast was defined as the spatial average of the local luminance standard deviation divided by the local mean luminance: $C_{\text{local}} = \left\langle \frac{\sigma_Y(p)}{\mu_Y(p)+\epsilon} \right\rangle_p$, where $\mu_Y(p) = (G_\sigma * Y)_p$ and $\sigma_Y(p) = \sqrt{(G_\sigma * Y^2)_p - (G_\sigma * Y)_p^2}$. Here, $G_\sigma$ is a normalized two-dimensional Gaussian kernel with standard deviation $\sigma = 3$ pixels, and $*$ denotes spatial convolution. Thus, $G_\sigma * Y$ is a Gaussian-weighted local average of the luminance image. We used reflection padding and $\epsilon = 10^{-6}$ to avoid division by zero. Finally, saturation was computed from the RGB channels as follows. Let $M_p = \max(R_p, G_p, B_p)$ and $m_p = \min(R_p, G_p, B_p)$. The saturation at pixel $p$ was defined as $S_p = \begin{cases} \dfrac{M_p - m_p}{M_p}, & M_p > 0, \\ 0, & M_p = 0, \end{cases}$ and mean saturation as $\overline{S} = \langle S_p \rangle_p$.

***E.2. Network training.*** As in the main text, we trained three CNN variants: a network trained only for classification (network C, identical to that used in the main text), a network trained exclusively to regress the eight features (network R+), and a network trained jointly for classification and regression of all eight features (network CR+).

***E.3. Results.*** **Fig. S8** is analogous to **Fig. 2b** in the main text and shows that network CR+ implements an effective joint code: it matches the classification performance of network C while achieving regression performance comparable to network R+.

**Figs. S9-S10** show the regression error $E$, its decomposition into local error $E_{\text{loc}}$ and the local-global error gap $\Delta E$, and the key theoretical quantities underlying the gap, $E_c$, $E_s$, $E_o$, SNR, and alignment $a$, across the main layers of networks C and CR+. These results support the same conclusions as in the main text. For every feature, network CR+ reduces the local-global gap by several orders of magnitude relative to network C at the feature layer. Importantly, our theory continues to capture the trend of the local-global gap across layers when eight features are jointly encoded, in most cases with good quantitative agreement. The local error can decrease even across the layers of network C. However, for image statistics that do not describe object position or extent, such as contrast and saturation, this decrease is less clearly monotonic than for the bounding-box features. Mean luma shows no monotonic improvement and is decoded most accurately directly from pixels, as might be expected given that it is a simple function of pixel values. This is consistent with the intuition that the improvement in local error for bounding-box features is driven, at least in part, by the spatial inductive bias of the CNN.

**Fig. S11** is analogous to **Fig. 5** in the main text and leads to the same conclusion: regression optimization in network CR+ occurs through a subtle reorganization of feature encoding within manifolds, without substantially altering manifold shapes or coarse classification-relevant properties such as manifold radius and dimensionality. **Fig. S11a** shows that the PC spectrum of network CR+ is virtually unchanged relative to network C. By contrast, network R+ exhibits a much steeper spectrum, with variance concentrated in roughly as many leading PCs as there are encoded features. **Fig. S11b** shows that, for all features examined, network CR+ increases the overlap between the feature-encoding directions and leading manifold PCs relative to network C. Nevertheless, each feature remains distributed across several PCs rather than becoming aligned with a single leading PC. **Fig. S11c** shows that average manifold alignment increases only modestly in network CR+ relative to network C and remains substantially lower than in network R+. The increase from C to CR+ is larger than that from C to CR, consistent with CR+ encoding a larger number of category-independent features. Finally, **Fig. S11d** shows that manifold radius and dimensionality, as defined by manifold-capacity theory (3, 8), are virtually unchanged between networks C and CR+, whereas both change substantially in network R+.

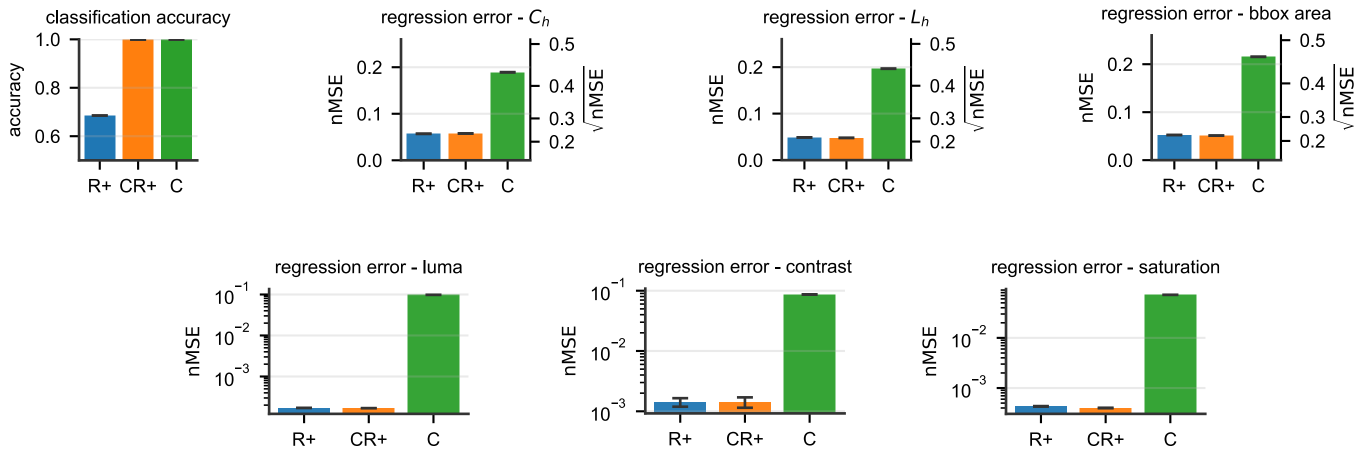

**Fig. S8.** Results of linearly decoding object category and regressing category-independent features from the feature layer of networks R+, CR+ and C. Error bars indicate the SEM across cross-validation (CV) splits. We show one example bounding box center and length parameters ($C_\text{h}$and $L_\text{h}$); results for $C_\text{v}$and $L_v$ are analogous. For $C_\text{h}$, $L_\text{h}$ and area we show nMSE in linear scale, with corresponding $\sqrt{\text{nMSE}}$ shown on the right y-axis for convenience. For luminance, contrast and saturation, we use a logarithmic scale since the discrepancy in performance between network C and networks CR+ or R+ is higher.

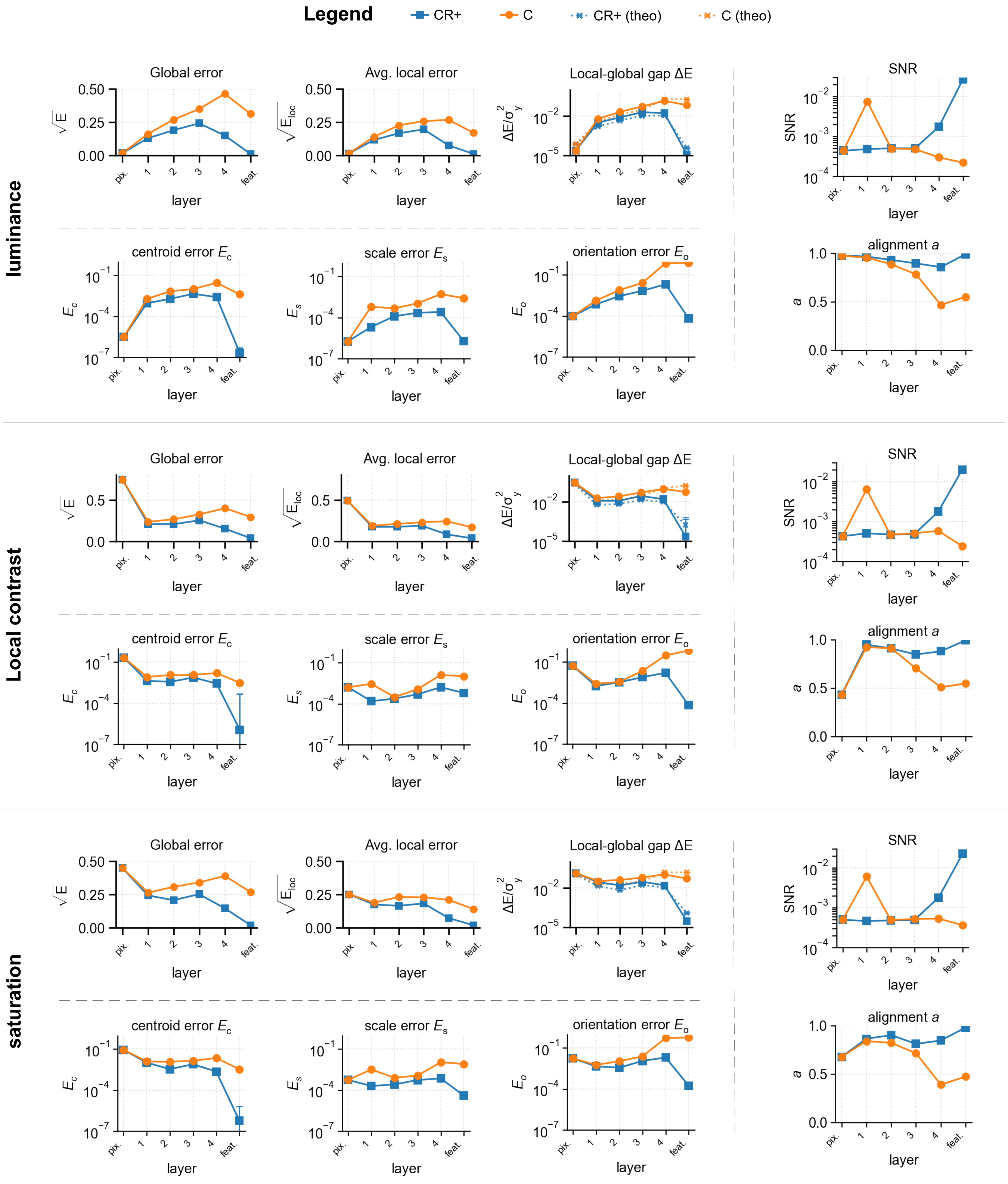

**Fig. S9.** Regression performance of luminance, contrast, and saturation across the layers of networks C (orange circles) and CR+ (blue squares). At each layer, readouts were trained after random projection to 2048 dimensions, matching the feature layer. Regression is shown as normalized global error $E$, local error $E_{\text{loc}}$, and local–global gap $\Delta E$. For the local-global gap, we additionally overlay our theoretical prediction (x markers and dotted lines). We also show the key theoretical quantities underlying the gap, $E_c$, $E_s$, $E_o$, $\text{SNR}$, and alignment $a$. Note that, unlike in the main text, here we plot the variance-normalized local–global gap $\Delta E$ rather than the relative gap $\Delta E/E_{\text{loc}}$. This avoids division by $E_{\text{loc}}$, which for these features can be small and nonmonotonic across layers, particularly in network CR+, and makes changes in $\Delta E$ easier to visualize. Consistently, $E_c$, $E_s$, and $E_o$ are shown on the same normalized-MSE scale rather than as fractions of $E_{\text{loc}}$. They are therefore normalized by the total variance of the target labels, as are $E$, $E_{\text{loc}}$, and $\Delta E$. Following the main text, the scale and orientation contributions, $E_s$ and $E_o$, are additionally multiplied by the linearized label variance $\hat{\sigma}^2$. Error bars are propagated SEM across CV splits and, where applicable, 5 random projection seeds.

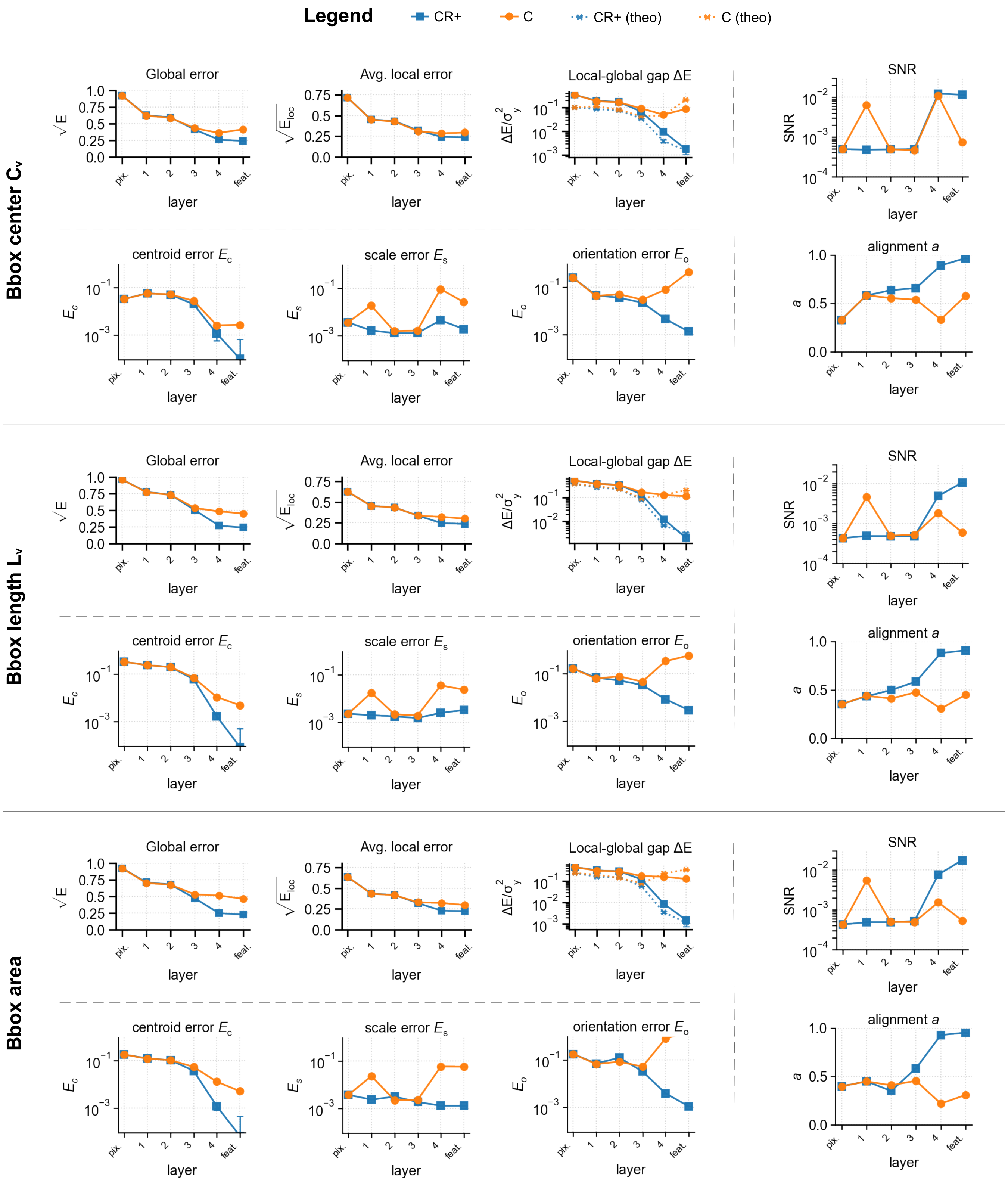

**Fig. S10.** Same as **Fig.** S9, but for bounding box area, and one example bounding box center and length parameters ($C_v$ and $L_v$).

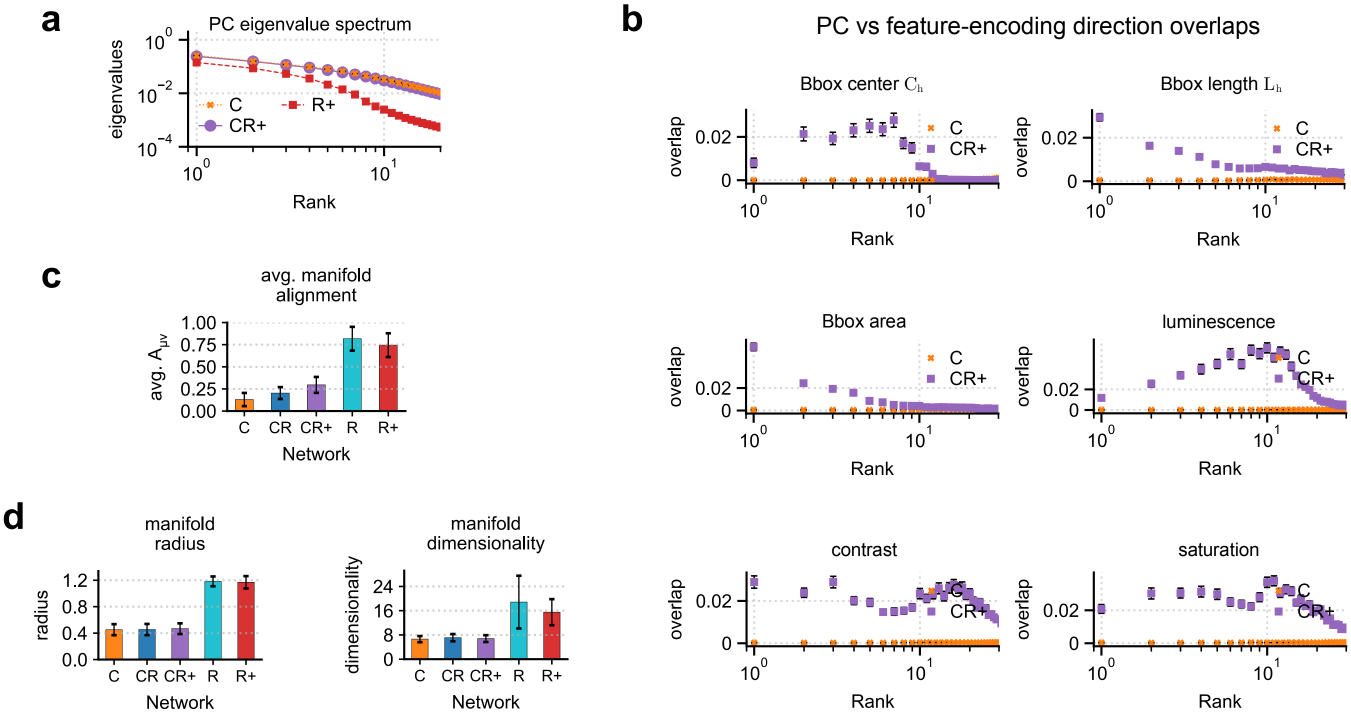

**Fig. S11.** Geometry optimization strategy. (a) PC eigenvalue spectrum (eigenvalue vs rank) for category manifolds in network C (orange), CR+ (purple), and R+ (red), averaged across categories (error bars are SEM). For each manifold, the eigenvalues were normalized by the manifold total variance averaged across networks. (b) Squared cosine of the angle between the feature-encoding direction $\hat{w}^{\mu}$ and the PC directions of manifold $\mu$ (ordered by rank) for networks C (orange) and CR+ (purple), averaged across categories (error bars are SEM). We show one example bounding box center and length parameters ($C_h$and $L_h$), results for $C_v$and $L_v$ are analogous. (c) Manifold alignment, as quantified by the RV coefficient between pairs of manifold covariance matrices, averaged across all pairs of categories (error bars are SD). (d) Manifold radius and dimensionality, as defined by manifold-capacity theory (3, 8), averaged across categories (error bars are STD).

### F. Further details on the effect of subsampling neural units.

***Effect of subsampling on the local error.*** In the main text, we examined the effects of subsampling object categories $P$ and neural units $N$ on the local-global error gap $\Delta E$. Here we report the effect of subsampling on the other component of the global regression error: the local error $E_{\mathrm{loc}}$. Because the local error $E_{\mathrm{loc}}$ measures within-category regression performance, subsampling the number of categories does not systematically affect it, aside from increased estimation noise. We therefore focus on the effect of subsampling neural units, which we report in **Fig. S12**.

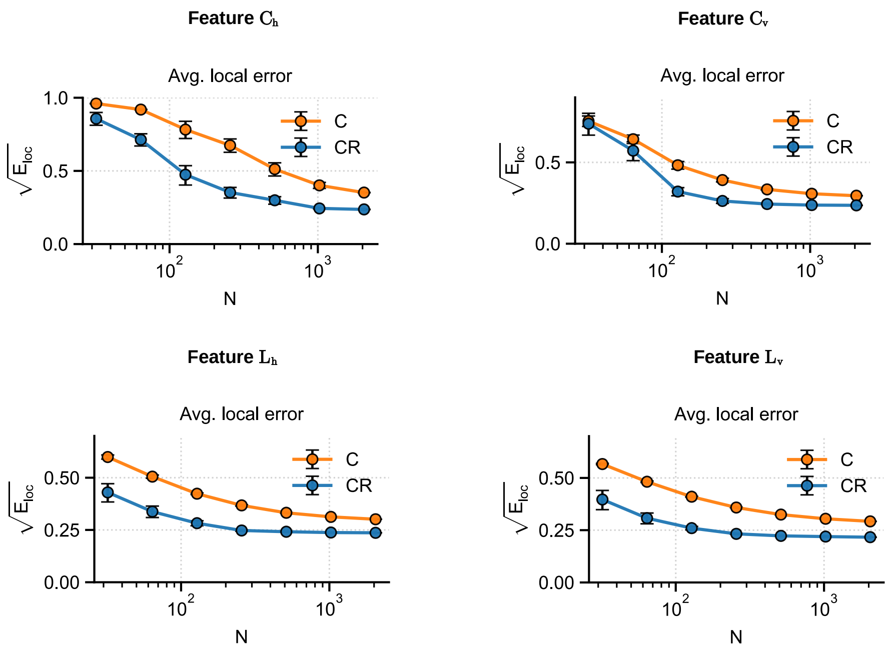

**Fig. S12.** Local regression error $E_{\text{loc}}$ for networks C (orange) and CR (blue) on our image dataset, as a function of the number $N$ of randomly subsampled units (out of 2048 feature-layer units).

***Effect of limited images per category on the estimation of the local-global gap.*** In the main text, we reported $\Delta E/E_{\mathrm{loc}}$ for regression of bounding-box features from the neural representations elicited by the image set used in (1, 2) to obtain macaque ventral-stream recordings (**Fig. 6c**). Specifically, we showed results for the recorded units in IT and V4, as well as for networks C and CR subsampled to 168 units, matching the number recorded in IT. Since all units are available in C and CR, one could in principle also study how $\Delta E/E_{\mathrm{loc}}$ varies with the number $N$ of subsampled units, mirroring the analysis performed in the main text on our image dataset (**Fig. 6b**). However, this analysis is much less informative for the image set used in (1, 2), because that dataset contains only 400 images per category. When $N$ becomes comparable to, or larger than, 400, the local regressors begin to overfit, artificially increasing the estimated local error $E_{\mathrm{loc}}$ and thereby underestimating the local–global gap. This effect is clearly visible in **Fig. S13**: beyond sufficiently large $N$, the local error reverses its expected downward trend and instead increases with $N$. As a consequence, the estimated local–global gap is progressively underestimated and can even become negative. More generally, this analysis highlights an important requirement for future experiments aiming to estimate the local–global error gap reliably: the number of images per category should exceed the number of recorded units, so as to avoid this overfitting regime.

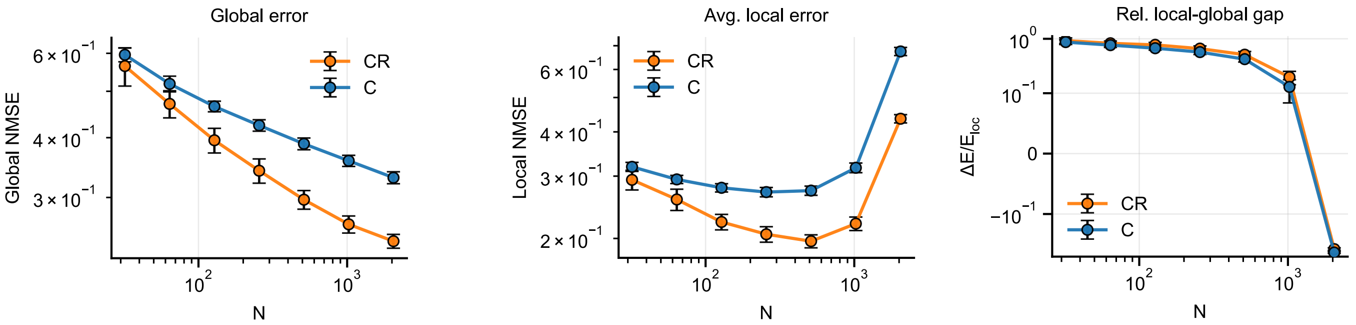

**Fig. S13.** Global error, local error, and relative local–global gap in the regression of bounding-box parameter $L_h$, shown as a function of the number $N$ of subsampled units, for networks C and CR on the image set used in (1, 2).

**G. Main text results for other bounding-box features.** Here we report the same kind of results presented in the main text, for bounding-box features that were not shown in the main text.

***Feature-encoding direction vs PCs overlaps.*** **Fig. S14** reports the overlap of the local feature-encoding direction with the top PCs of the corresponding category-manifold, averaged across categories. The trend is the same as discussed in the main text (cf. **Fig. 5**): in network CR compared to C, the feature-encoding directions are much more aligned with top manifold PCs.

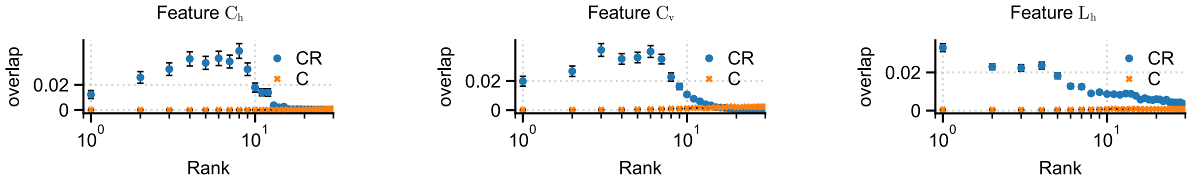

**Fig. S14.** Feature-encoding direction vs PCs overlap. Squared cosine of the angle between the feature-encoding direction $\hat{w}^\mu$ and the PC directions of manifold $\mu$ (ordered by rank) for networks C (orange) and CR (blue), averaged across categories (error bars are SEM).

***Decoding performance along the visual hierarchy.*** **Fig. S15** reports regression performance and associated theoretical measures for the bounding-box feature $C_\mathrm{h}$. We focus on $C_\mathrm{h}$ because $C_\mathrm{v}$ shows the same qualitative behavior; similarly, in the main text and in an SI section above we reported $L_\mathrm{h}$, while $L_\mathrm{v}$ follows an analogous trend. Overall, the across-layer behavior of center coordinates is similar to that of box lengths described in **Fig. S7**, with one minor difference: in network C, the regression error for the center coordinates increases at the final transition from the last convolutional layer to the feature layer. A plausible explanation is that center coordinates rely more directly on the spatial arrangement of activity across the convolutional feature maps, because they encode *where* the object is located in the image. Global average pooling removes much of this spatial information by collapsing each feature map into a single scalar, and therefore can impair linear decoding of object position. By contrast, box lengths are invariant to object translation and thus may be less sensitive to the pooling operation. In network C, which is optimized only for classification, there is no pressure to recode positional information into a form that survives the final pooling step. In network CR, by contrast, explicit regression supervision can promote a reorganization of the last convolutional representation such that center-coordinate information remains linearly accessible after average pooling.

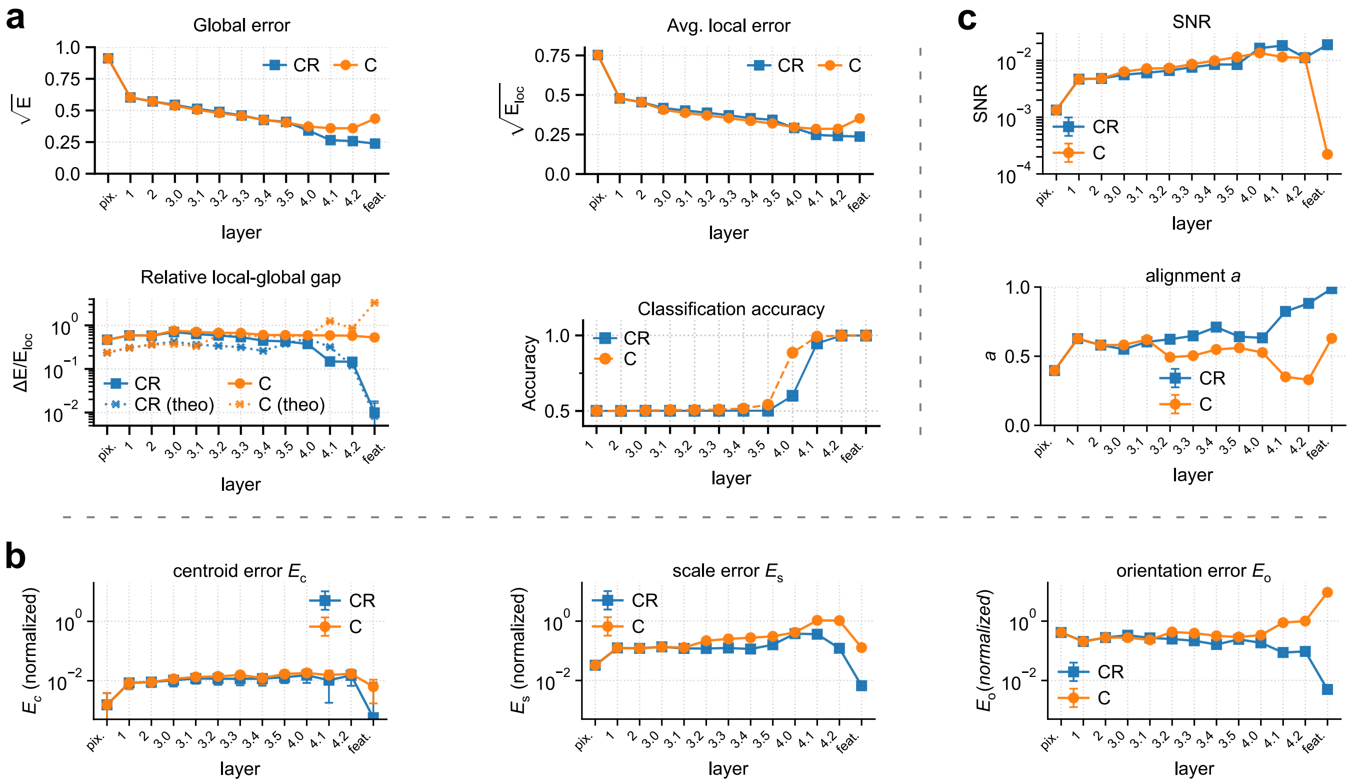

**Fig. S15.** Decoding performance along the visual hierarchy. (**a**) Regression performance for the bounding-box parameter $C_h$ and classification performance, using linear readouts from progressively deeper layers of networks C (orange circles, solid line) and CR (blue squares, solid line). For regression, we report the global regression error, as well as its decomposition into local error $E_{loc}$ and relative local-global gap $\Delta E/E_{loc}$. For the gap, we also report our theory prediction (x markers, dotted lines). Note that global and local error are reported as root nMSE. (**b**) The three contributions to the relative local-global gap, as identified by our theory: centroid error $E_c$, orientation error $E_o$, scale error $E_s$. These quantities are normalized in the same way as explained in **Fig. 4**. (**c**) The two elements contributing to the orientation error: SNR, and alignment $a$ of the local feature-encoding directions.

***Subsampling effects.*** **Fig. S16** reports the effect of subsampling neural units on the two components of global regression error—local error $E_{\rm loc}$ and local-global gap $\Delta E$—for the bounding-box features not shown in the main text. The trend is the same as observed in the main text (cf. **Fig. 6b**): for a sufficiently small number of subsampled units $N$, the local-global gap $\Delta E$ in networks C and CR becomes indistinguishable. **Fig. S17** reports the analysis of regression performance on the images and corresponding neural recordings in (1), for the other bounding-box features not shown in the main text. The trend is the same as observed in the main text (**cf. Fig. 6c**): IT, V4, as well as networks C and CR subsampled to 168 units, matching the number recorded in IT, all show comparable values of $\Delta E/E_{\rm loc}$, as well as comparable contributions from each of the three error sources, with $E_{\rm o}$ and $E_{\rm c}$ emerging as the dominant terms.

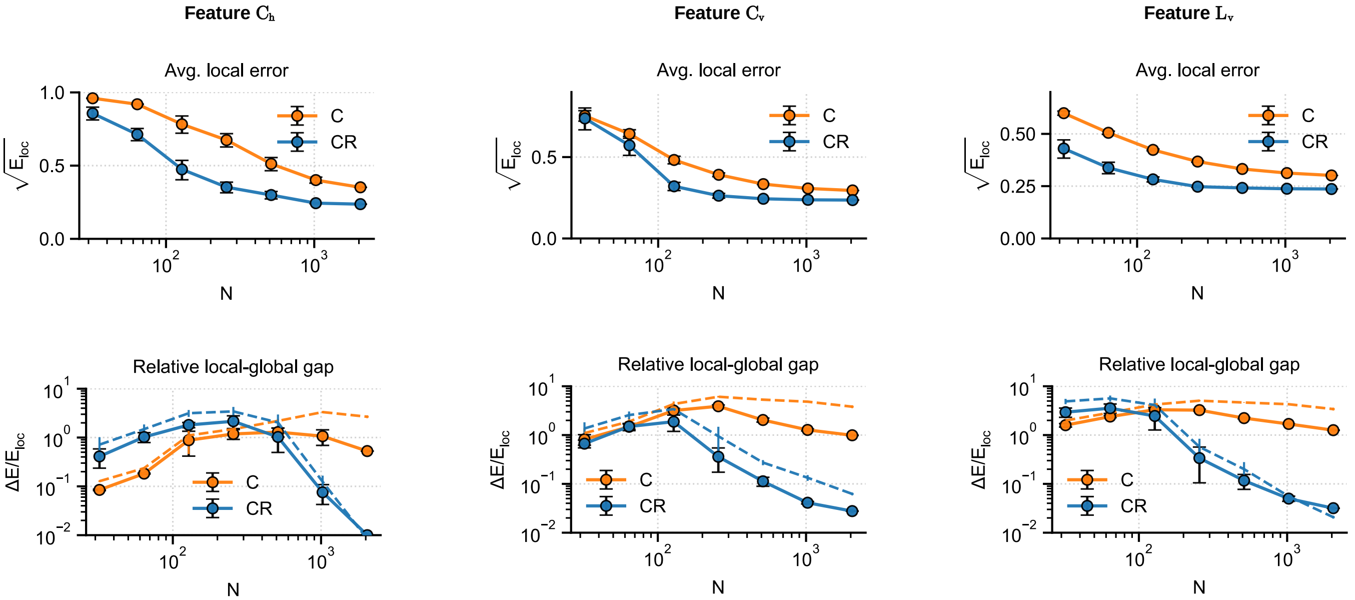

**Fig. S16.** Effect of subsampling neural units. $E_{\text{loc}}$ and $\Delta E/E_{\text{loc}}$ for regressing the bounding-box features $(C_h, C_v, L_v)$ from networks C (orange) and CR (blue) on our dataset, as a function of the number $N$ of randomly subsampled units (out of 2048 feature-layer units). For the gap, we also report our theory prediction (x markers, dotted lines). Error bars denote SEM across cross-validation splits and random unit subsamples.

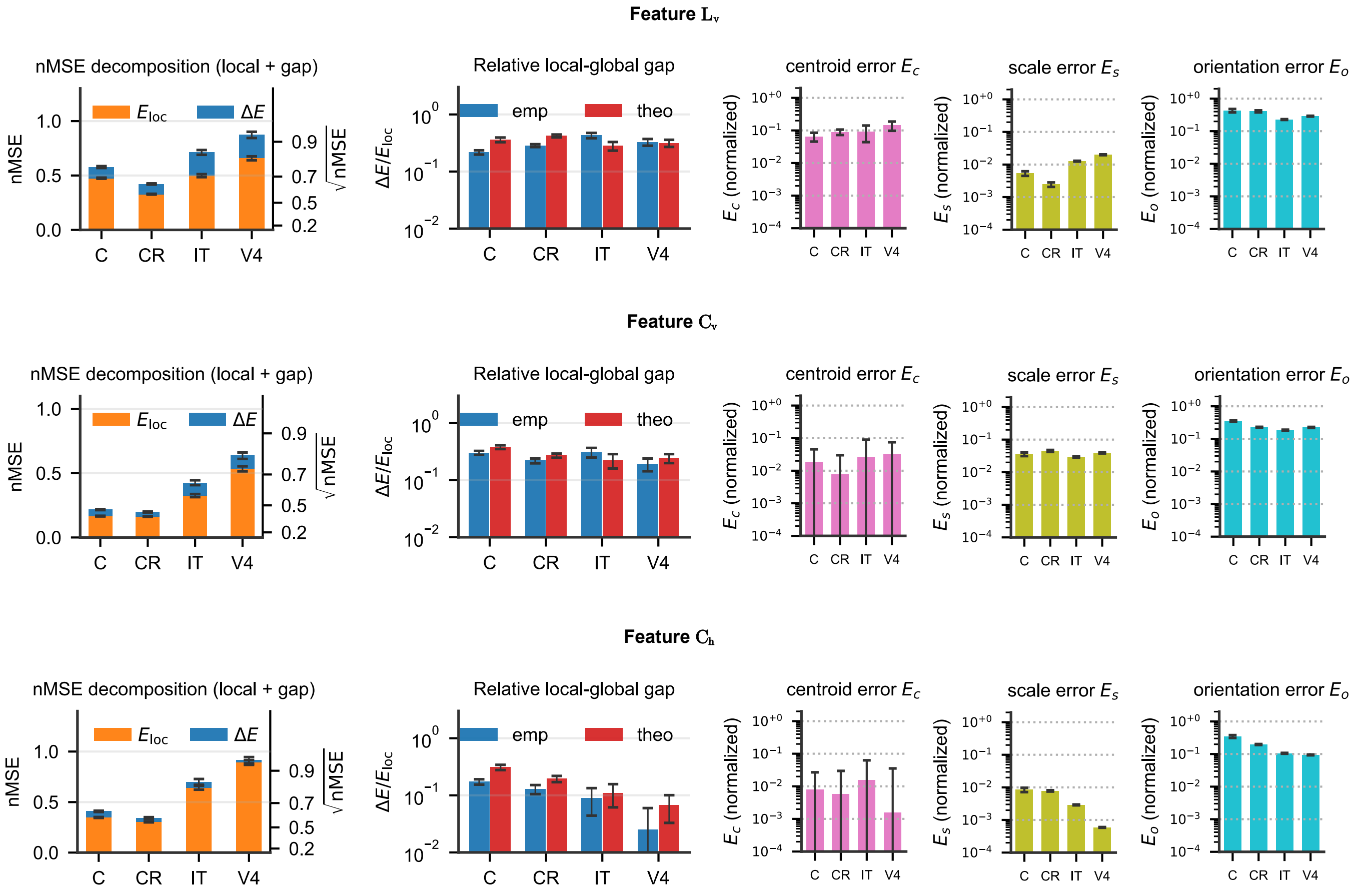

**Fig. S17.** Analysis of regression performance on the images and corresponding neural recordings found in (1). We report the analysis of regression from the IT and V4 units recorded in (1), as well as 168 randomly subsampled units from the best performing layer of networks C and CR. For features $C_h$ and $C_v$, the best performing layer is layer 4, for feature $L_v$ it is the feature layer. Error bars denote SEM across cross-validation splits and, where applicable, random unit subsamples.

**H. Robustness of main-text findings on different training runs.** Here we show that our main-text findings are robust across different training runs of networks C and CR. We report several main-text results for four distinct realizations of each network, corresponding to four random seeds used to initialize the training runs. The runs differed in the random selection of mini-batches during gradient-descent training and in the random initialization of the readout heads. A pretrained ResNet-50 backbone was used in every run, as explained in the main text. **Fig. S18** reports the global regression error, its decomposition into the local error and the local-global error gap, the theoretical prediction of this gap, and the theory-derived quantities that predict it. All these results depend only weakly on the training seed. Likewise, **Fig. S19** reports the other measures of manifold geometry presented in the section "Manifold geometry reveals optimization strategy" and the associated Supplementary Information. Again, we observe only a weak dependence on the training seed.

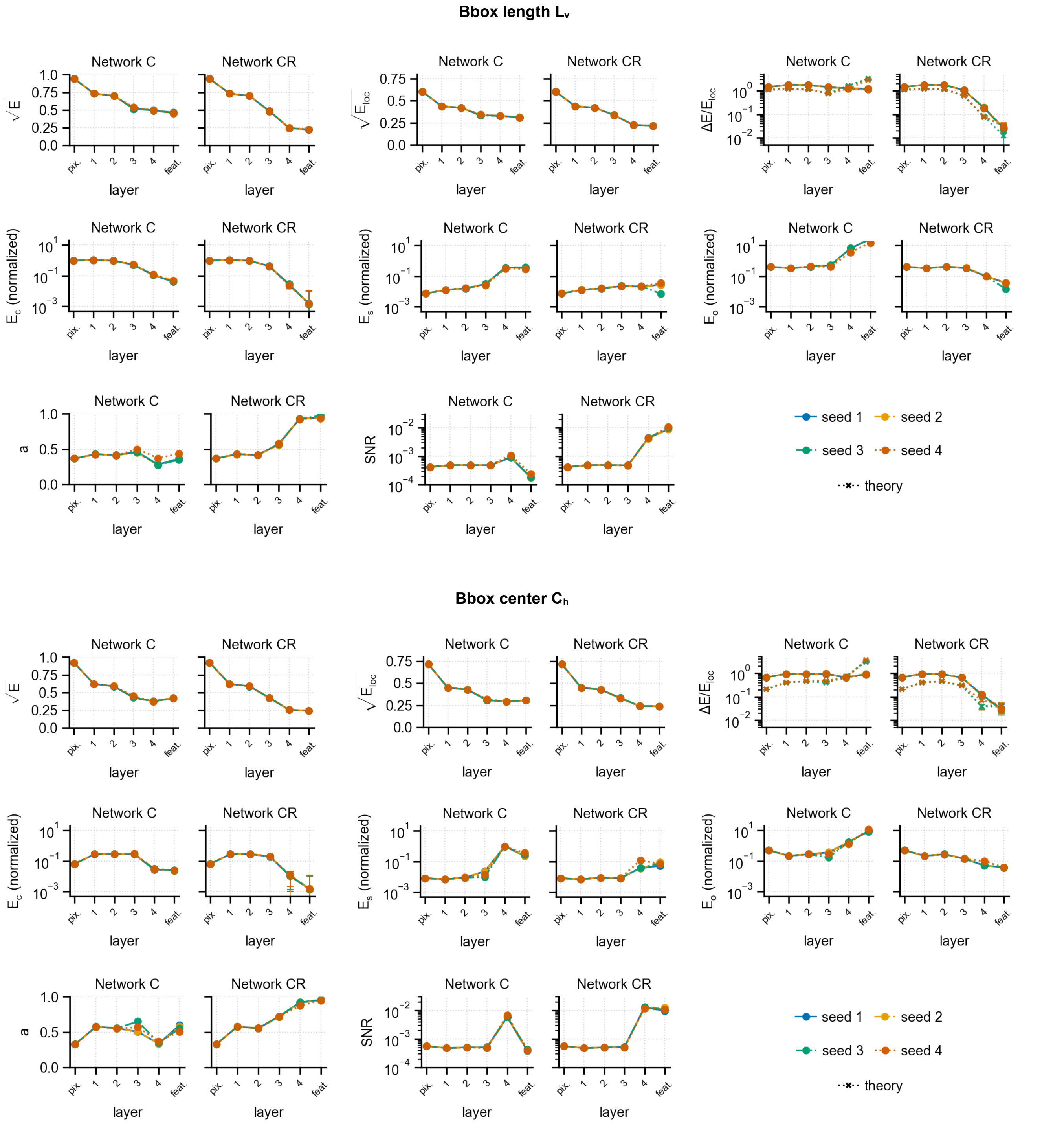

**Fig. S18.** Regression results across four training seeds. Colors and line styles indicate the different seeds (see legend). For the local-global error gap, theoretical predictions are shown using cross markers and dashed lines in the color corresponding to each training seed. We report results for one bounding-box length and one bounding-box center parameters. Results for the other coordinates are analogous.

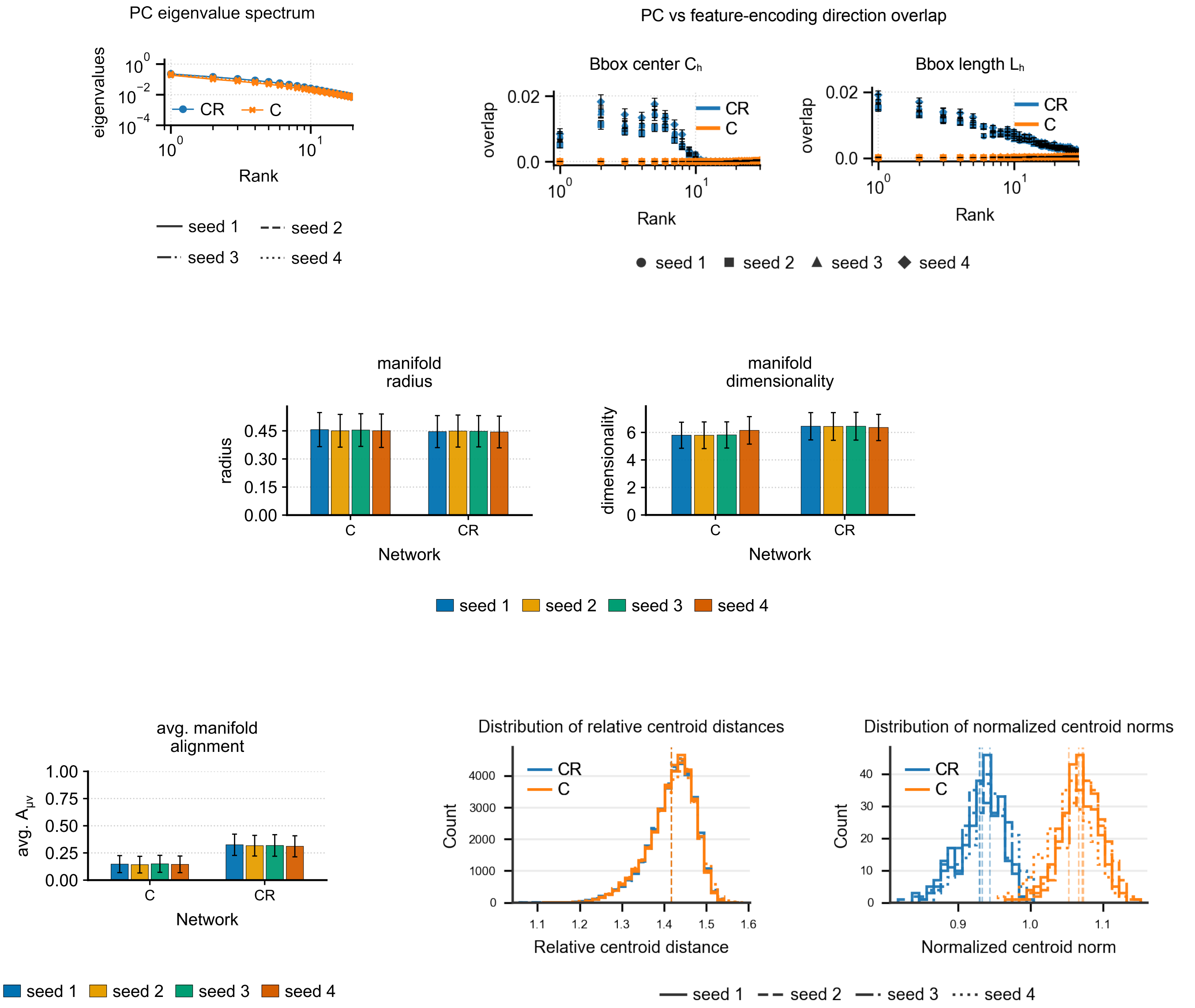

**Fig. S19.** Manifold geometry across four training seeds. Depending on the quantity shown, training seeds are distinguished by different colors, line styles, or markers (see legends).

**I. Across-layer trends in regression error are independent of random-projection dimensionality.** The across-layer trends in regression performance reported in **Fig. 3e** were obtained after randomly projecting each flattened convolutional representation to 2,048 dimensions, matching the dimensionality of the final feature layer. Here, we show that the qualitative improvement in regression performance across layers is not a trivial consequence of this dimensionality reduction. **Fig. S20** reports regression performance for pixels and the four main convolutional layers of networks C and CR as a function of random-projection dimensionality, extending to the full, unprojected representation of each layer. Even at the full dimensionality, regression performance improves monotonically from pixels to layer 4. Thus, the across-layer trend reported in the main text is preserved independently of projection size.

To extend the analysis beyond $N = 8192$, we replaced the exact ridge-regression solver used in the main text, whose computational cost increases rapidly with dimensionality, with minibatch stochastic gradient descent. This procedure optimizes the same ridge-regression objective, with regularization selected through the same nested cross-validation scheme. Regressors were trained for six epochs using batches of 200 images and a cosine learning-rate schedule decreasing from $10^{-3}$ to $10^{-5}$. Input representations were centered and scaled before optimization to improve numerical stability. Solid curves show SGD regression, whereas dotted curves show exact ridge regression over the computationally feasible range. SGD closely matches the exact solution across projection sizes, with only slightly higher error, supporting its use at larger dimensionalities.

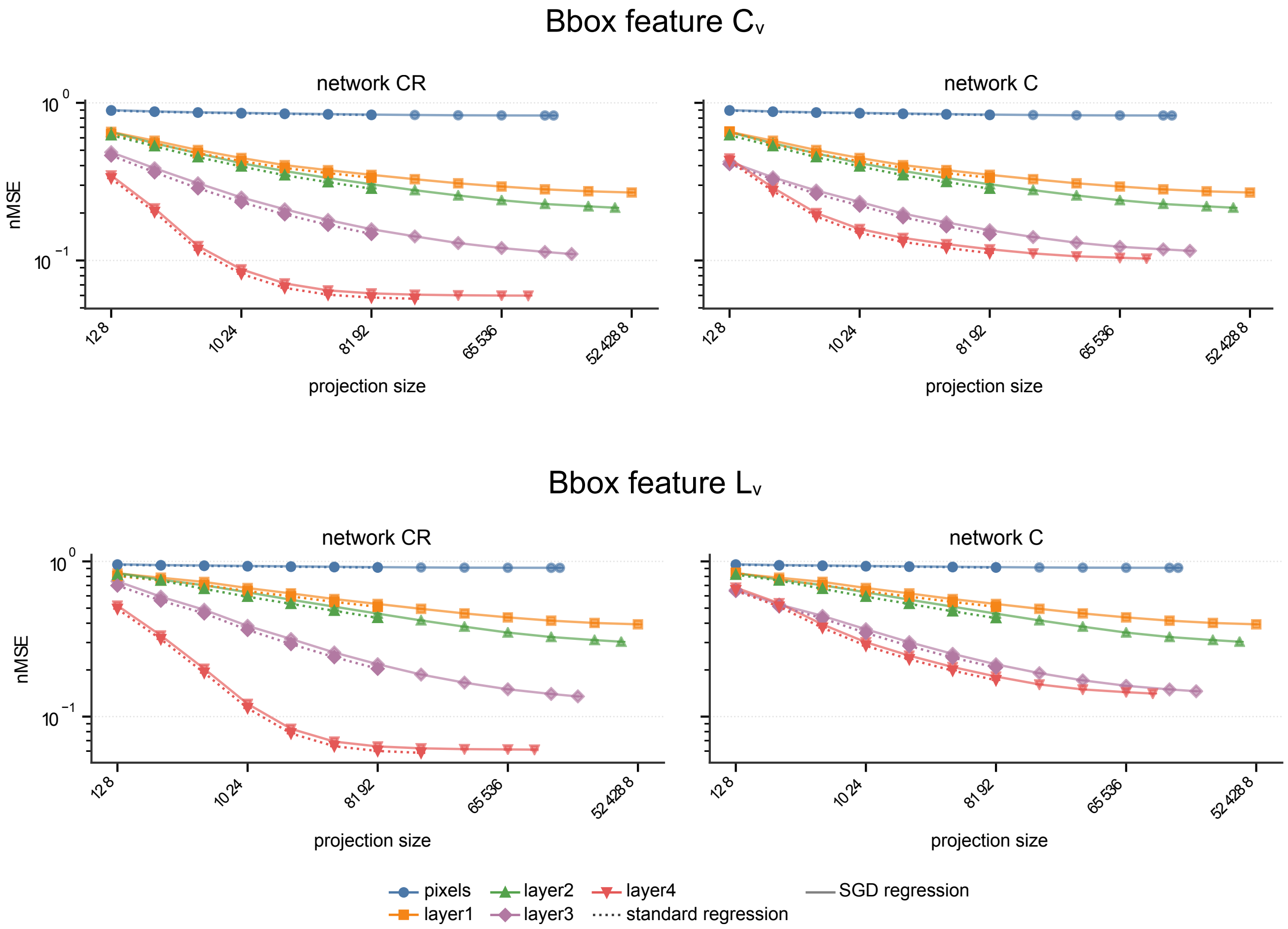

**Fig. S20.** Regression performance across projection sizes. Normalized mean-squared regression error (nMSE) for pixels and convolutional layers 1–4 of networks CR (left) and C (right), as a function of projection size. Solid lines show ridge regression optimized using SGD; dotted lines show exact ridge-regression solutions. Error bars indicate the SEM across outer cross-validation folds and 5 random projection seeds. We show regression for one bounding-box center ($C_v$) and one bounding-box length ($L_v$). The results for the other center and length parameters are analogous.

**J. Across-layer trends do not depend on the layer-freezing training schedule.** In training the CNNs, we found that gradually unfreezing the weights of the ResNet-50 layers through the epochs, from the feature layer downstream to layer 1, yielded models with slightly better regression performance. In particular, the models reported in the main text were trained by unfreezing a new layer each epoch, in sequence: the feature layer, layer 4, and layer 3, while layers 2 and 1 were kept frozen throughout the training. Here in the SI, we show that the trends across layers reported in the main text (cf. **Fig.** S7) are virtually unchanged when network CR is trained end to end with all layers unfrozen from the very first epoch, rather than with the layer-unfreezing schedule described above. We call this network CR*. **Fig. S21** shows regression performance as a function of layer depth for networks CR, C and CR*. We still observe the trends reported in the main text: a sharp rise in regression performance emerges only from layer 4, as well as an appreciable distinction in regression performance between networks C and CR/CR*.

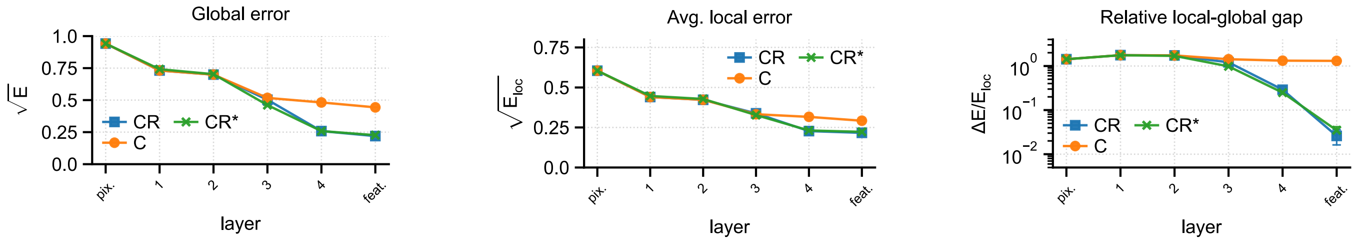

**Fig. S21.** Regression performance along the visual hierarchy. Regression performance for the bounding-box parameter $L_{\mathrm{h}}$ using linear readouts from progressively deeper layers of networks C (orange circles, solid line) and CR (blue squares, solid line) and CR* (green cross, solid line). At each layer, linear readouts were trained on activations randomly projected to 2048 dimensions, to match the dimensionality of the final feature layer. Note that global and local error are reported as root nMSE.

### 2. Local regression with shared regularization

To estimate linear feature encoding within each category, used to compute $E_{\text{loc}}$ and the geometric quantities entering our theory, we jointly fit category-specific regressors ($W^\mu \in \mathbb{R}^N, b^\mu \in \mathbb{R}$) by minimizing

$$E_{\text{common}} = \frac{1}{P}\sum_{\mu=1}^{P}\left\langle \left(W^{\mu\top}x^\mu + b^\mu - y^\mu\right)^2 \right\rangle_\mu + \gamma L(\{W^\mu\}),$$

with

$$L(\{W^\mu\}) = \frac{1}{P}\sum_{\mu=1}^{P}\left\|W^\mu - \bar{W}\right\|^2, \qquad \bar{W} = \frac{1}{P}\sum_{\nu=1}^{P}W^\nu.$$

Here, $x^\mu \in \mathbb{R}^N$and $y^\mu \in \mathbb{R}$are respectively the representation and target feature value for category $\mu$, $P$ is the number of categories, and $\langle\cdot\rangle_\mu$ denotes averaging over training samples from category $\mu$. The regularization strength $\gamma$ was selected by nested cross-validation as described in Materials and Methods. We computed $E_{\text{loc}}$ as the average test nMSE across categories.

This procedure yielded $E_{\text{loc}}$ values statistically compatible with those obtained from independent per-category ridge regressors, while providing more stable estimates of the local regression weights $\{W^\mu\}$, whose across-category variability in norm and direction determines the scale and orientation errors, $E_{\text{s}}$ and $E_{\text{o}}$. With independent fits, broad ranges of ridge penalties produced statistically indistinguishable performance, leading to substantial variability in the selected penalties and hence unnecessary variability in the estimated weights. The shared regularizer reduces this estimation variability by shrinking each category-specific regressor toward the mean regressor $\bar{W}$, allowing more stable estimation of the geometric quantities entering our theory.

### 3. Review of manifold alignment (RV coefficient) and centered kernel alignment

**A. Manifold alignment (RV coefficient).** For each category $\mu$, let $\Sigma^\mu \in \mathbb{R}^{N\times N}$ denote the covariance matrix of its centered representations. We quantify the alignment between manifolds $\mu$ and $\nu$ using the RV coefficient

$$A_{\mu\nu} = \frac{\text{Tr}(\Sigma^\mu\Sigma^\nu)}{\sqrt{\text{Tr}[(\Sigma^\mu)^2]\,\text{Tr}[(\Sigma^\nu)^2]}}.$$

This is essentially the normalized Frobenius inner product between the two covariance matrices. The measure is normalized so that $A_{\mu\nu} \in [0,1]$. Intuitively, $A_{\mu\nu}$ is large when manifolds $\mu$ and $\nu$ are aligned, i.e., their axes of variability (PCs) point in similar directions and carry comparable variance. $A_{\mu\nu} = 1$ when the covariance matrices are identical up to an overall scale.

To remove the overlap expected between randomly oriented manifolds, in the figures we report the baseline-corrected coefficient

$$\widetilde{A}_{\mu\nu} = \frac{A_{\mu\nu} - A^{\text{rand}}_{\mu\nu}}{1 - A^{\text{rand}}_{\mu\nu}}, \qquad A^{\text{rand}}_{\mu\nu} = \frac{\sqrt{D_\mu D_\nu}}{N},$$

where

$$D_\mu = \frac{\text{Tr}(\Sigma^\mu)^2}{\text{Tr}[(\Sigma^\mu)^2]}$$

is the participation-ratio dimensionality of manifold $\mu$. Thus, $\widetilde{A}_{\mu\nu} = 0$ corresponds to the random-alignment baseline and $\widetilde{A}_{\mu\nu} = 1$ to perfect alignment.

**B. Centered kernel alignment.** We use linear centered kernel alignment (CKA) to compare representations from two networks (9). Let $X \in \mathbb{R}^{M\times N_X}$ and $Y \in \mathbb{R}^{M\times N_Y}$ contain the representations of the same $M$ images in the two networks. Their centered Gram matrices are

$$K_{\text{c}} = HXX^\top H, \qquad L_{\text{c}} = HYY^\top H, \qquad H = I - \frac{1}{M}\mathbf{1}\mathbf{1}^\top.$$

Linear CKA is the normalized alignment between these Gram matrices,

$$\text{CKA}(X,Y) = \frac{\text{Tr}(K_{\text{c}}L_{\text{c}})}{\sqrt{\text{Tr}(K_c^2)\,\text{Tr}(L_{\text{c}}^2)}}.$$

It ranges from 0 to 1, with 1 indicating identical representational similarity structure up to transformations to which linear CKA is invariant. We report the debiased version of CKA, obtained by replacing the Hilbert–Schmidt independence criterion terms entering CKA with their unbiased estimators (9, 10). Nevertheless, because of the large number of images used here, biased and debiased CKA agree to several significant digits.

### 4. Image dataset

**A. Image generation process.** Here we provide a more in-depth description of the image-generation procedure for our dataset. Images were generated using a multi-stage pipeline that combines text-to-image generation (Stable Diffusion XL, SDXL (11, 12)), object detection (CerberusDet (13)), and image outpainting (Stable Diffusion v1.5, SD1.5 (14–16)) to produce single-object images with controlled object placement and controlled distribution of bounding-box coordinates. The pipeline to generate one image in our dataset consists of the following three steps (cf. **Fig. S22**).

***Step 1: Object generation.*** We first generate a "seed" image of a single object in the desired category using Stable Diffusion XL. The goal of this step is simply to obtain a clear, unobstructed, single-instance depiction of the target object in a contextually appropriate scene, without yet enforcing any control over object position or scale (i.e., control over the object's bounding-box parameters). To make large-scale generation feasible, we use SDXL-Lightning weights (model release identifier: ByteDance/SDXL-Lightning) with the SDXL base checkpoint (stabilityai/stable-diffusion-xl-base-1.0), enabling generation in only 4 denoising steps. This allowed us to synthesize the full dataset within a few days on a 10×A100 GPU cluster. For a given category named <category>, we use the following prompt: "A full-body, photorealistic shot of a single <category>. The <category> is the clear main subject, fully in the scene, and unobstructed. The background is detailed, contextually appropriate, but does not distract.". We also use the negative prompt: "clutter, multiple objects, occlusion, low quality, blurry, distorted, unrealistic, extra limbs, cropped, artifacts, persons, people, humans, mannequin".

***Step 2: Object placement.*** The goal of this step is to enforce controlled object location and size by mapping the seed image from step 1 onto a new canvas so that the main object occupies a newly sampled bounding box. We first obtain the object's bounding box in the seed image using CerberusDet, and treat this detected box as the object's initial bounding box. If CerberusDet does not detect any instance of the object, we discard the seed image and return to step 1. We then sample a target bounding box on a fixed 512×512 canvas as follows. First, we sample a target maximum side length $L_{\max} \sim \text{Unif}\,[200, 500]$ (in pixel units) and isotropically rescale the seed image such that the detected bounding-box's maximum side length matches the newly sampled one, that is we ensure $\max(L_h, L_v) = L_{\max}$. Next, conditioned on the resulting $(L_h, L_v)$, we sample the target bounding-box center uniformly over all centers that would keep the box inside the canvas: $C_h \sim \text{Unif}\,[L_h/2, 512 - L_h/2]$ and $C_v \sim \text{Unif}\,[L_v/2, 512 - L_v/2]$. In sum, object size (via $L_{\max}$) is sampled uniformly over a prescribed range, and object position is sampled uniformly conditional on size, subject to the newly sampled bounding-box staying fully within the image. To realize this target bounding-box, we paste the rescaled seed image onto a blank 512×512 canvas so that the object bounding-box matches the target one. Some regions of the canvas will remain uncovered by the rescaled image, and are filled in Step 3 via outpainting.

***Step 3: Outpainting.*** The goal of this step is to fill in the uncovered regions introduced in Step 2, producing a natural-looking full 512×512 image while preserving the placed object and its newly sampled bounding box. We achieve this by outpainting the uncovered regions using an outpainting variant of SD1.5. Starting from the 512×512 canvas produced in Step 2, we construct an outpainting mask that marks the uncovered regions. To reduce visible seams at the boundary between original and synthesized content, we blur the mask edges with a Gaussian blur. Before outpainting, we initialize the undefined regions with reflection padding to provide the generative model with plausible color and texture statistics and reduce sharp discontinuities at the boundary between the pasted seed image and the area to be synthesized. Technically, we fill each blank side by repeatedly tiling a mirror-reflected strip of pixels taken only from the portion of the seed image between the pasted-image border and the object's bounding-box edge (never from inside the bounding box), so that the object itself is not reflected—avoiding to encourage the outpainting model to generate additional instances of the object. We finally outpaint the masked regions using SD1.5 inpainting (stable-diffusion-v1-5/stable-diffusion-inpainting) with an LCM scheduler and the latent-consistency/lcm-lora-sdv1-5 LoRA, using 4 denoising steps, guidance scale 0, and strength 0.95. The outpainting prompt is the same category prompt from Step 1 with the suffix: 'The background extends naturally from the existing scene. Seamless transition between regions, no visible boundaries. Consistent lighting and focus throughout the entire image." The negative prompt is identical to Step 1. Finally, we rerun CerberusDet on the outpainted 512×512 image and accept the sample only if the detector returns exactly one instance of the desired category, so that images with multiple instances of the object are not accepted.

***Category selection.*** As mentioned above, we generated images in 265 categories out of the 365 categories in the Objects365 taxonomy, selected among those that could be generated reliably with our pipeline. To select these categories, we ran a diagnostic pass generating 50 images per category and measured how often the pipeline failed due to CerberusDet either not detecting the target object or detecting multiple instances. We excluded categories with high failure rates, since their low yield would have substantially increased the runtime and compute required to produce 20,000 accepted images per category. Furthermore, we visually inspected the 50 images in each category and excluded categories that systematically produced implausible or low-quality images, such as ambiguous or inconsistent object identity, severe artifacts, or the presence of multiple salient objects. For a few categories, SDXL introduced characteristic artifacts that were *consistent* across images within that category (e.g., "knife" images often contained forked or unusually curved blades). Since our goal is a dataset in which images within each category share a consistent semantic label, rather than strict realism, we did not exclude such categories. During this inspection, we also identified categories for which CerberusDet consistently localized the intended object but systematically assigned a different class label. Because we used CerberusDet only to extract bounding boxes, we defined for each category a small set of accepted detector labels and accepted an image if the detector returned exactly one instance with a label in that set.

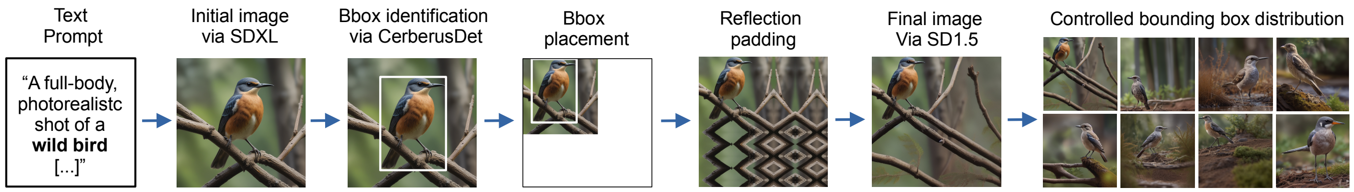

**Fig. S22.** Generation process for an image in our dataset. From left to right: an initial image of an object in the desired category (e.g. "wild bird") is generated from a text prompt using Stable Diffusion XL (11, 12), with categories drawn from the taxonomy of Objects365 (17), an image dataset for object detection. The object's bounding box is then estimated by CerberusDet (13), an Objects365-trained detector. The image is then isotropically rescaled and pasted onto a white canvas so that the detected box matches a target box whose parameters $(C_h, C_v, L_h, L_v)$ have been drawn uniformly (conditional on preserving aspect ratio and remaining inside the canvas). The remaining white area is filled by reflection-padding the border region around the pasted image, and the resulting canvas is used as the conditioning image for a second round of generation with a Stable Diffusion 1.5 inpainting model (14–16), yielding the final sample. The method allows generation of images with a controlled distribution of bounding boxes.

**B. Example images and bounding-box statistics .** Here we report additional example images from our dataset (**Figs. S23-S26**) together with the bounding-box statistics of the full dataset.

**Fig. S27a** shows the global bounding-box statistics across all images and categories. To interpret this figure, recall the bounding-box sampling procedure. First, the maximum side length $L_{\max} = \max(L_h, L_v)$ is sampled uniformly, $L_{\max} \sim \mathrm{Unif}[200, 500]$, in pixel units (the full image is $512 \times 512$ pixels), consistent with the approximately uniform distribution shown in **Fig. S27a**, left. The aspect ratio $L_h/L_v$ is then kept fixed with respect to the original seed image in order to preserve the natural appearance of the object. Across all categories, this yields an overall distribution of aspect ratios that is roughly symmetric around 1 (**Fig. S27a**, middle-left). The center coordinates are then sampled uniformly over all positions that keep the bounding box inside the canvas, namely $C_h \sim \mathrm{Unif}[L_h/2,\, 512 - L_h/2]$ and $C_v \sim \mathrm{Unif}[L_v/2,\, 512 - L_v/2]$. This produces distributions that are symmetric around the image center (**Fig. S27a**, middle-right and right). Note, however, that the center coordinates are more concentrated near the image center overall, because the allowed range of center positions depends on the bounding-box size. In particular, boxes with larger side lengths can occupy only a narrower range of center positions around the image center, since the box must remain fully inside the image. As a result, center coordinates closer to the image center occur more frequently in the dataset.

We also report bounding-box statistics on a per-category basis (**Fig. S27b**). The bounding-box sampling procedure does not depend explicitly on object category, except through the natural range of aspect ratios associated with different categories (for example, cars tend to be more extended along the horizontal axis, whereas penguins tend to be more extended along the vertical axis). This is reflected in the high uniformity across categories in the distributions of maximum side length and center coordinates. By contrast, aspect ratio is much more category-dependent, both in its within-category mean and in its within-category spread (standard deviation).

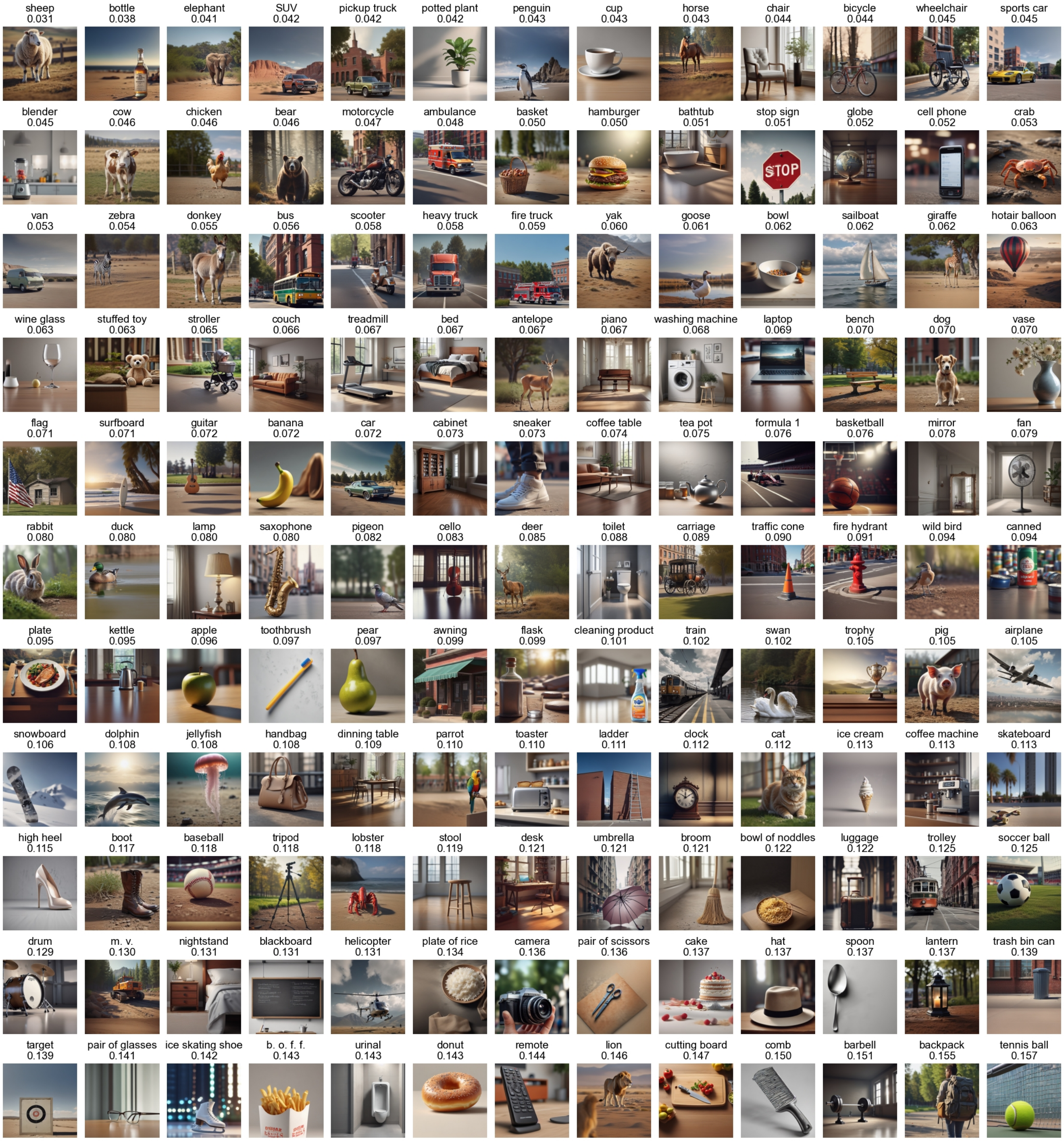

**Fig. S23.** All categories in our dataset (part 1; continues in Fig. S24). One representative image per category is shown. Below the category label, we report the test regression $\sqrt{\text{nMSE}}$ achieved by network CR for that category, averaged across all images in the category and across the four bounding box parameters. Categories are ordered by ascending $\sqrt{\text{nMSE}}$.

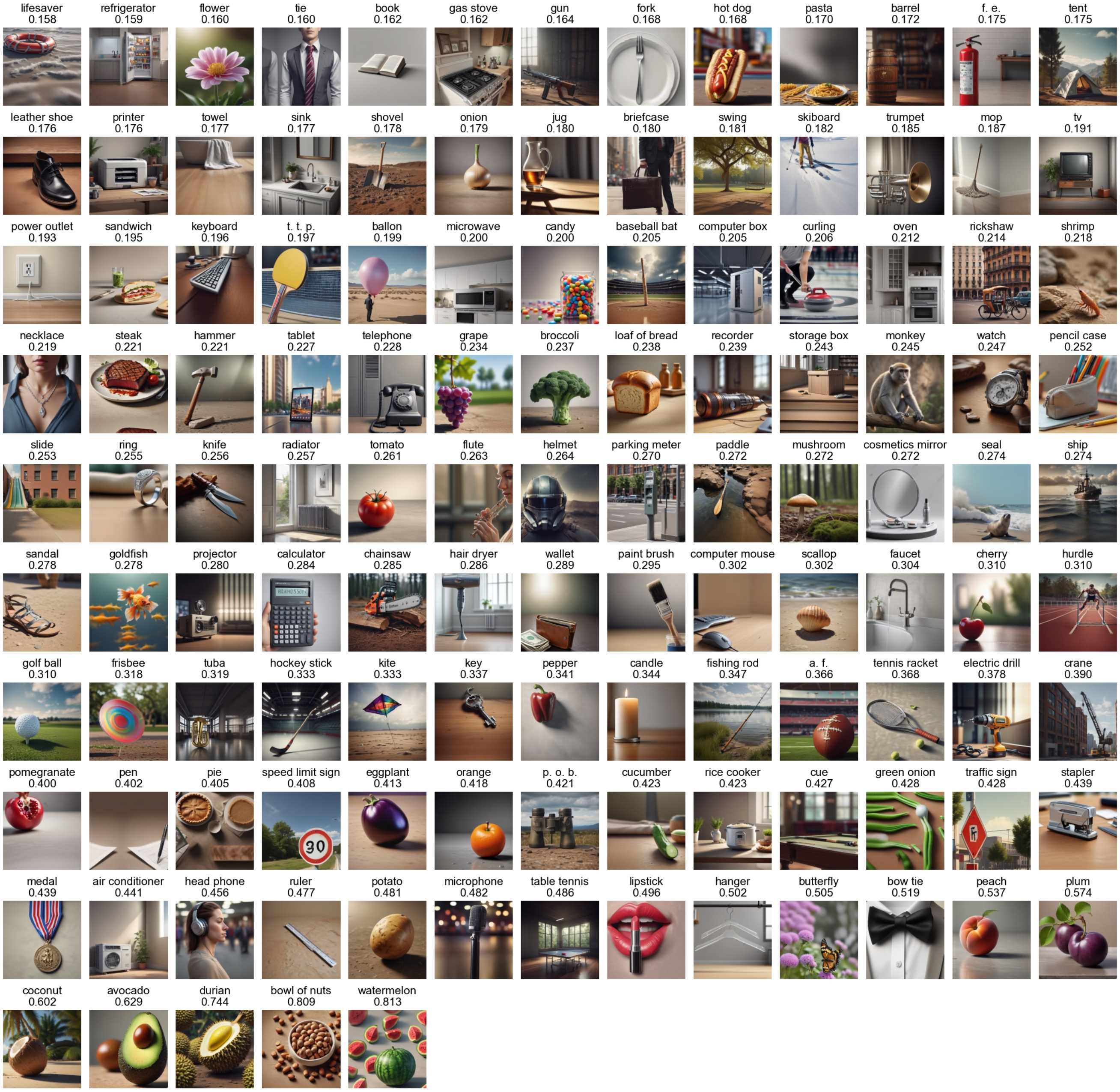

**Fig. S24.** All categories in our dataset (part 2; see Fig. S23).

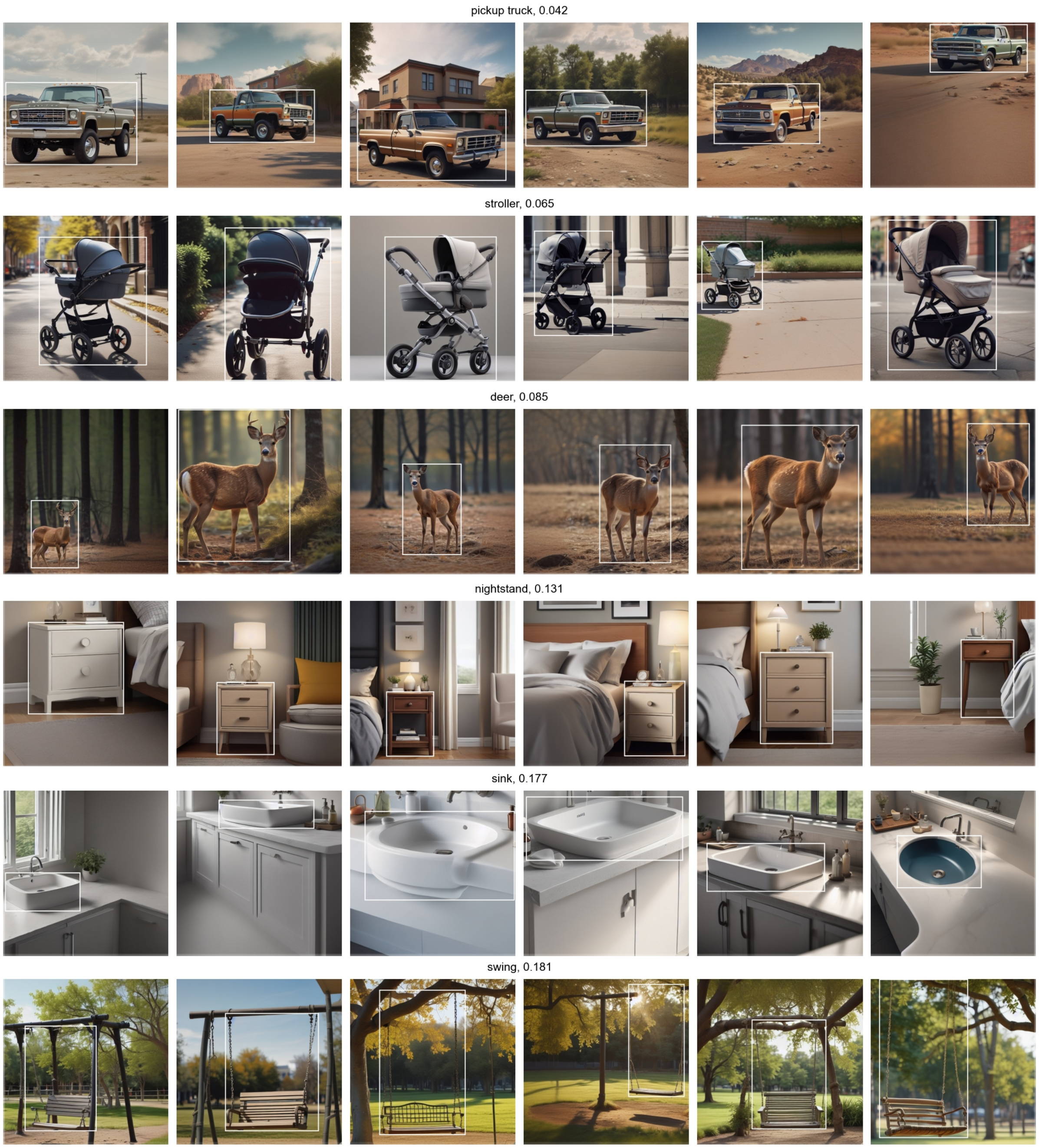

**Fig. S25.** Bounding-box variations (part 1; continues in Fig. S26). For a given category, we show six randomly selected images from our dataset, with overlaid ground-truth bounding-box in white. Next to the category label, we report the test regression $\sqrt{\text{nMSE}}$ achieved by network CR for that category, averaged across all images in the category and across the four bounding box parameters. Categories are ordered by ascending $\sqrt{\text{nMSE}}$.

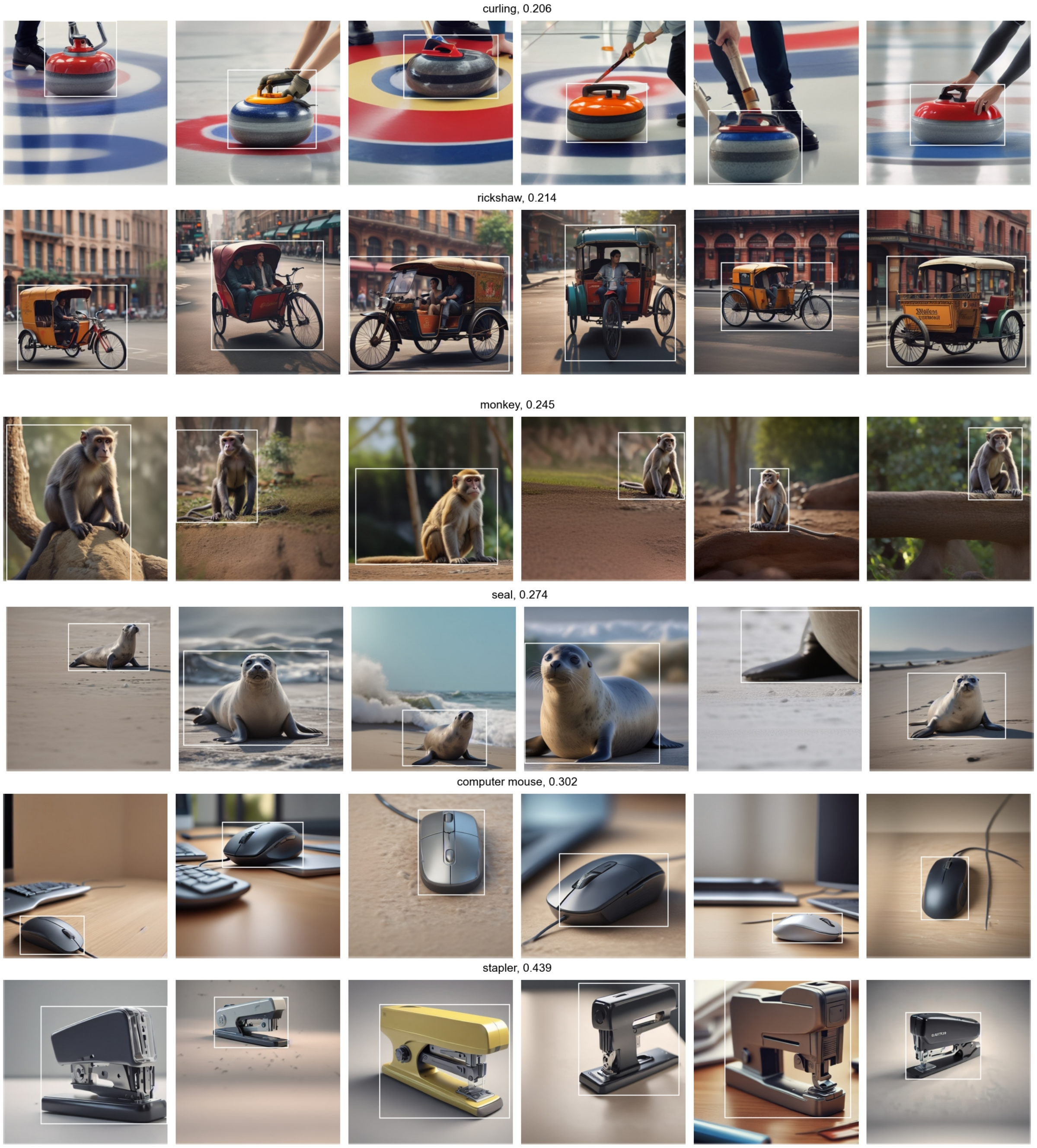

**Fig. S26.** Bounding-box variations (part 2; see Fig. S25).

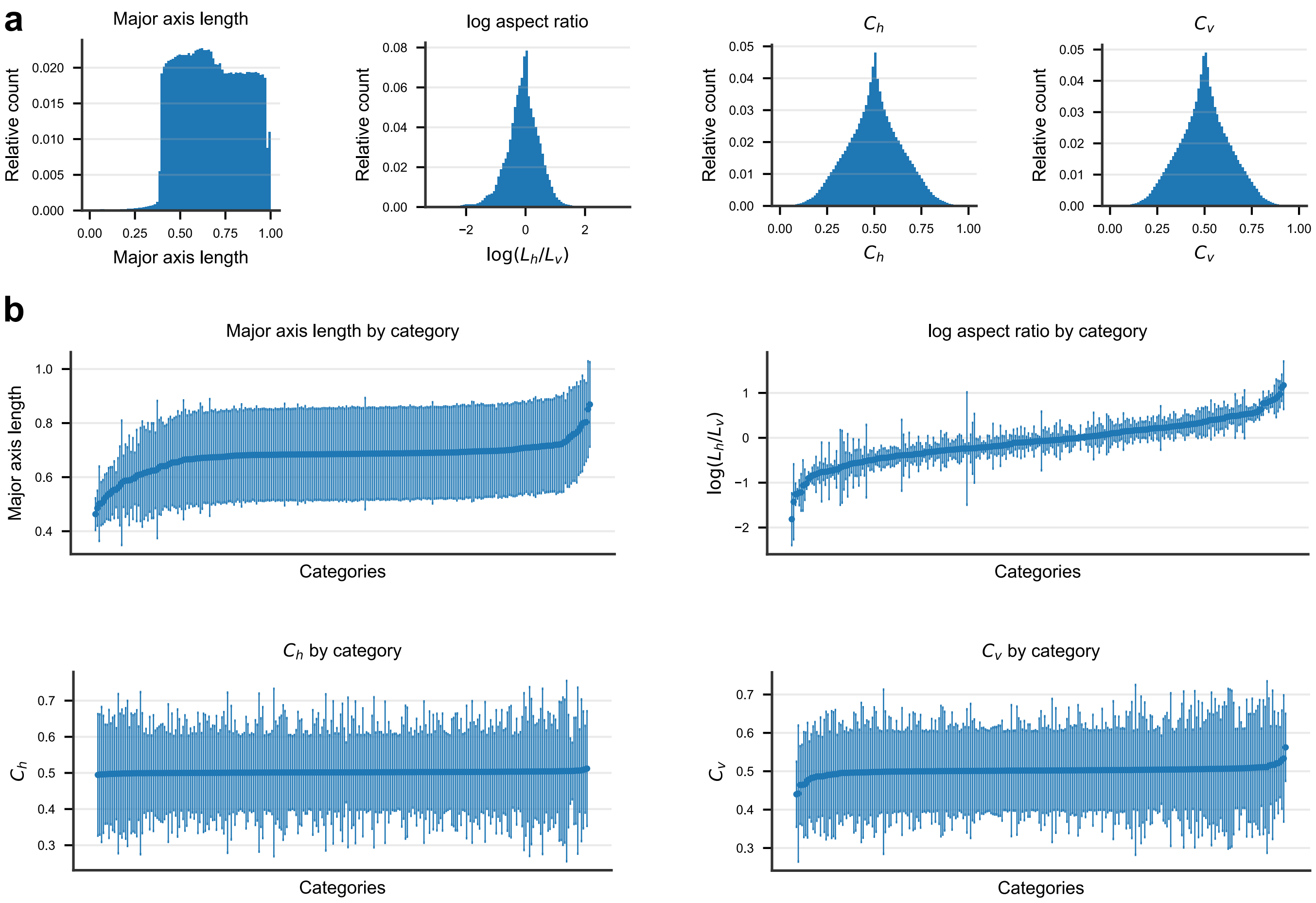

**Fig. S27.** Bounding-box statistics in our image dataset. (**a**) Histogram across all images and all categories of major axis length max $(L_h, L_v)$, log aspect ratio log $(L_h/L_v)$, and center coordinates $C_h$ and $C_v$. Major axis length and center coordinates are given in relative units, i.e. in pixel units divided by $512$, which is the image side-length in pixel units. (**b**) Same quantities shown in (a), but on a per-category basis. Categories are ordered on the x-axis based on the within-category average of the considered quantity. Markers indicate the within-category average and error bars the within-category standard deviation of the considered quantity.

### 5. Theory derivation and numerical validation

**A. Rewriting of the local-global gap.** ***Definitions.*** For convenience, we first recall the definitions of the global and local regression errors introduced in the Methods.

Without loss of generality, we consider globally centered data and labels, that is $\left\langle \langle x^\mu \rangle_\mu \right\rangle_C = 0$ and $\left\langle \langle y^\mu \rangle_\mu \right\rangle_C = 0$, where $\langle \ldots \rangle_\mu$ denotes averaging over all images within the $\mu$-th category, and $\langle \ldots \rangle_C = \text{Avg}_\mu [\ldots]$ denotes averaging over categories.

The global regression MSE is

$$E = \min_W \left\langle \left\langle \left( W^\top x^\mu - y^\mu \right)^2 \right\rangle_\mu \right\rangle_C , \tag{2}$$

while the local regression MSE for category $\mu$ is

$$E^\mu = \min_{(W^\mu, b^\mu)} \left[ \left\langle \left( W^{\mu\top} x^\mu + b^\mu - y^\mu \right)^2 \right\rangle_\mu \right] \tag{3}$$

The average local error is $E_{\text{loc}} = \langle E^\mu \rangle_C$, and the local-global error gap is defined as $\Delta E = E - E_{\text{loc}}$. We denote by $\hat{y}^\mu = W_*^{\mu\top} x^\mu + b_*^\mu$ the prediction of the optimal local regressor for category $\mu$, where $(W_*^\mu, b_*^\mu) = \text{argmin}_{(W^\mu, b^\mu)} \left[ \left\langle \left( W^{\mu\top} x^\mu + b^\mu - y^\mu \right)^2 \right\rangle_\mu \right]$.

***Result.*** We now show that $\Delta E$ can be rewritten as a global regression error on the locally linearized labels $\hat{y}^\mu$, that is

$$\Delta E = \min_W \left[ \left\langle \left\langle \left( W^\top x^\mu - \hat{y}^\mu \right)^2 \right\rangle_\mu \right\rangle_C \right] \tag{4}$$

***Derivation.*** For each category $\mu$, define the local residual

$$\epsilon^\mu = y^\mu - \hat{y}^\mu . \tag{5}$$

We can show that the residual satisfies

$$\langle \epsilon^\mu \rangle_\mu = 0, \qquad \langle x^\mu \epsilon^\mu \rangle_\mu = 0. \tag{6}$$

Indeed, the optimal local regressor is given by

$$W_*^\mu = K^{\mu +} d^\mu, \qquad b_*^\mu = \langle y^\mu \rangle_\mu - W_*^{\mu\top} \langle x^\mu \rangle_\mu , \tag{7}$$

where $K^\mu = \left\langle \delta x^\mu \delta x^{\mu\top} \right\rangle_\mu$, $d^\mu = \langle \delta x^\mu \delta y^\mu \rangle_\mu$, $\delta x^\mu = x^\mu - \langle x^\mu \rangle_\mu$, $\delta y^\mu = y^\mu - \langle y^\mu \rangle_\mu$, and $K^{\mu +}$ denotes the Moore-Penrose pseudoinverse. Therefore,

$$\epsilon^\mu = y^\mu - \hat{y}^\mu = \delta y^\mu - W_*^{\mu\top} \delta x^\mu . \tag{8}$$

Averaging immediately gives $\langle \epsilon^\mu \rangle_\mu = 0$. Moreover,

$$\langle x^\mu \epsilon^\mu \rangle_\mu = \langle \delta x^\mu \epsilon^\mu \rangle_\mu = d^\mu - K^\mu W_*^\mu = d^\mu - K^\mu K^{\mu +} d^\mu = 0, \tag{9}$$

which completes the derivation of Eq. (6).

Since $\hat{y}^\mu$ is itself a linear function of $x^\mu$, from Eq. (6) it follows that

$$\langle \hat{y}^\mu \epsilon^\mu \rangle_\mu = 0. \tag{10}$$

We can then write, for any global regressor $W$,

$$\begin{aligned} E(W) &= \left\langle \left\langle \left( W^\top x^\mu - y^\mu \right)^2 \right\rangle_\mu \right\rangle_C = \left\langle \left\langle \left( W^\top x^\mu - \hat{y}^\mu - \epsilon^\mu \right)^2 \right\rangle_\mu \right\rangle_C \\ &= \left\langle \left\langle \left( W^\top x^\mu - \hat{y}^\mu \right)^2 \right\rangle_\mu \right\rangle_C - 2 \left\langle \left\langle \left( W^\top x^\mu - \hat{y}^\mu \right) \epsilon^\mu \right\rangle_\mu \right\rangle_C + \left\langle \left\langle \epsilon^{\mu 2} \right\rangle_\mu \right\rangle_C \end{aligned} \tag{11}$$

where in the last step we expanded the square. The cross term vanishes, since

$$\left\langle \left( W^\top x^\mu - \hat{y}^\mu \right) \epsilon^\mu \right\rangle_\mu = W^\top \langle x^\mu \epsilon^\mu \rangle_\mu - \langle \hat{y}^\mu \epsilon^\mu \rangle_\mu = 0. \tag{12}$$

By construction, $\left\langle \epsilon^{\mu 2} \right\rangle_\mu = E^\mu$, and therefore $\left\langle \left\langle \epsilon^{\mu 2} \right\rangle_\mu \right\rangle_C = E_{\text{loc}}$. Minimizing $E(W)$ with respect to $W$, we thus obtain

$$E = \min_W \left\langle \left\langle \left( W^\top x^\mu - \hat{y}^\mu \right)^2 \right\rangle_\mu \right\rangle_C + E_{\text{loc}}, \tag{13}$$

which immediately implies Eq. (4).

**B. Derivation of the local-global gap in terms of manifold geometry.** ***Generative model.*** For convenience, we first recall the definitions and assumptions of the manifold generative model introduced in the Methods. As explained in the Methods, we derive the local-global gap in the locally-centered case $\langle x^\mu \rangle_\mu = 0, \langle y^\mu \rangle_\mu = 0, \forall \mu$, and then quantify the effect of manifold centroids by the centroid error. Without loss of generality, each point in manifold $\mu$ is parameterized as

$$x^\mu = \sum_{d=1}^{N} s_d^\mu u_d^\mu, \tag{14}$$

where $\{u_d^\mu\}_{d=1}^N$ is an orthonormal basis of $\mathbb{R}^N$, and $s_d^\mu$ is the coordinate along direction $d$. We choose

$$u_1^\mu = \frac{W_*^\mu}{\|W_*^\mu\|} \tag{15}$$

to be the feature-encoding direction, while the remaining directions $u_{d>1}^\mu$ are chosen to diagonalize the covariance in the subspace orthogonal to $u_1^\mu$. With this choice, the covariance of the manifold coordinates takes the form

$$\left\langle s^\mu s^{\mu\top} \right\rangle_\mu = \Sigma^\mu = \begin{pmatrix} \sigma_1^{\mu 2} & \Sigma_{12}^\mu & \cdots & \Sigma_{1N}^\mu \\ \Sigma_{21}^\mu & \sigma_2^{\mu 2} & 0 & 0 \\ \vdots & 0 & \ddots & 0 \\ \Sigma_{N1}^\mu & 0 & 0 & \sigma_N^{\mu 2} \end{pmatrix}. \tag{16}$$

Note that local centering of the manifolds implies $\langle s_d^\mu \rangle = 0$. Let us also define

$$r^\mu = \|W_*^\mu\| \tag{17}$$

as the scale of the feature encoding, and note that

$$\hat{y}^\mu = r^\mu s_1^\mu \tag{18}$$

We assume the following across-manifold statistics:

1. The feature-encoding directions $u_1^\mu$ share a common mean direction and isotropic second-order fluctuations across categories

$$\langle u_1^\mu \rangle_C = \sqrt{a} u_1 \tag{19}$$

$$\left\langle u_1^\mu u_1^{\mu\top} \right\rangle_C = a u_1 u_1^\top + \frac{1-a}{N}\mathbb{I} \tag{20}$$

   where $a \in [0,1]$ is the alignment strength, and $u_1 \in \mathbb{R}^N$ is a unit-norm vector representing the common mean direction.

2. All manifold directions with $d > 1$ are isotropically distributed up to their second order statistics, conditionally on being orthogonal to $u_1^\mu$ (as by their definition as the elements of an orthonormal basis). Concretely,

$$\langle u_d^\mu \rangle_C = 0 \tag{21}$$

$$\left\langle u_d^\mu u_{d'}^{\mu\top} \right\rangle_C = \frac{1}{N-1}\delta_{d,d'}\left(\mathbb{I} - \left\langle u_1^\mu u_1^{\mu\top} \right\rangle_C\right) = \frac{1}{N}\delta_{d,d'}\left(\mathbb{I} - a\, u_1 u_1^\top\right) + \mathcal{O}\left(\frac{1}{N^2}\right). \tag{22}$$

3. Aside from the above correlations, all manifold parameters are otherwise uncorrelated.

Note that no assumptions are made nor need to be made about higher order statistics of $u_d^\mu$, since regression by MSE minimization is a quadratic problem and therefore depends only on statistics up to second order.

***Asymptotic regime.*** Our main theory is derived in the asymptotic regime $P \to \infty$ with $N$ large but finite. Assuming $N$ is large, our theory neglects correction of order $\mathcal{O}\left(N^{-1}\right)$. In this regard, we assume the $\mathrm{SNR}^{-1} = \sum_d \left\langle \sigma_d^{\mu 2} \right\rangle_C / \left\langle \sigma_1^{\mu 2} \right\rangle_C$ to be extensive in $N$, $\mathrm{SNR}^{-1} = \mathcal{O}(N)$. This is important to keep the orientation error $E_\mathrm{o}$ of $\mathcal{O}(1)$. This choice is motivated by the empirical observation that the orientation and scale errors are of similar magnitude, and therefore the orientation error should be considered as an $\mathcal{O}(1)$ term, rather than neglected as an $\mathcal{O}\left(N^{-1}\right)$ term. We emphasize that this assumption is used as a controlled scaling prescription for our theoretical derivation. It should not be interpreted as a literal claim about the behavior of variance in arbitrarily large biological networks. Rather, it provides a convenient large-$N$ continuation of the empirical finite-$N$ regime in which we find the orientation error to remain finite.

***Result.*** For the generative model and asymptotic regime described above, and neglecting $\mathcal{O}\left(N^{-1}\right)$ corrections, we find

$$\Delta E = \hat{\sigma}^2 \left[1 - \frac{1}{(1+E_\mathrm{s})(1+E_\mathrm{o})}\right] \tag{23}$$

$$E_\mathrm{s} = \frac{\mathrm{Var}_\mu [r^\mu]}{\mathrm{Avg}_\mu [r^\mu]^2} \tag{24}$$

$$E_{\mathrm{o}} = \frac{1-a}{N \cdot a \cdot \mathrm{SNR}} \qquad [25]$$

where $\mathrm{SNR} = \frac{\langle \sigma_1^{\mu 2}\rangle_C}{\sum_{d=1}^{N} \langle \sigma_d^{\mu 2}\rangle_C}$, and $\hat{\sigma}^2 = \left\langle \left\langle \delta\hat{y}^{\mu 2}\right\rangle_\mu \right\rangle_C$ with $\delta\hat{y}^\mu = \hat{y}^\mu - \langle \hat{y}^\mu\rangle_\mu$.

***Derivation.*** Recall that

$$\Delta E = \min{}_W \left[\left\langle \left\langle \left(W^\top x^\mu - \hat{y}^\mu\right)^2 \right\rangle_\mu \right\rangle_C\right] \qquad [26]$$

minimizing w.r.t. $W$ we obtain

$$\Delta E = \hat{\sigma}^2 - d^\top K^+ d \qquad [27]$$

where

$$K = \left\langle \left\langle x^\mu x^{\mu\top}\right\rangle_\mu \right\rangle_C \qquad [28]$$

$$d = \left\langle \langle x^\mu \hat{y}^\mu\rangle_\mu \right\rangle_C \qquad [29]$$

$$\hat{\sigma}^2 = \left\langle \left\langle \hat{y}^{\mu 2}\right\rangle_\mu \right\rangle_C \qquad [30]$$

Under our assumptions of manifold statistics, we have, neglecting $\mathcal{O}\left(N^{-1}\right)$ corrections,

$$K = \left\langle \sigma_1^{\mu 2}\right\rangle_C \left(a u_1 u_1^\top + \frac{1-a}{N}\mathbb{I}\right) + \frac{1}{N}\sum_{d>1} \left\langle \sigma_d^{\mu 2}\right\rangle_C \left(\mathbb{I} - a u_1 u_1^\top\right) \qquad [31]$$

$$d = \langle r^\mu\rangle_C \left\langle \sigma_1^{\mu 2}\right\rangle_C \sqrt{a} u_1 \qquad [32]$$

$$\hat{\sigma}^2 = \left\langle r^{\mu 2}\right\rangle_C \left\langle \sigma_1^{\mu 2}\right\rangle_C \qquad [33]$$

We note that $d$ is an eigenvector of $K$, therefore we can readily derive

$$d^\top K^+ d = \frac{a \langle r^\mu\rangle_C^2 \left\langle \sigma_1^{\mu 2}\right\rangle_C^2}{a\left\langle \sigma_1^{\mu 2}\right\rangle_C + \frac{1-a}{N}\sum_d \left\langle \sigma_d^{\mu 2}\right\rangle_C} \qquad [34]$$

Normalizing by $\hat{\sigma}^2 = \left\langle r^{\mu 2}\right\rangle_C \left\langle \sigma_1^{\mu 2}\right\rangle_C$ we obtain

$$\frac{d^\top K^+ d}{\hat{\sigma}^2} = \frac{\langle r^\mu\rangle_C^2}{\langle r^{\mu 2}\rangle_C} \frac{1}{1 + \frac{1}{N}\frac{1-a}{a}\frac{\sum_d \langle \sigma_d^{\mu 2}\rangle_C}{\langle \sigma_1^{\mu 2}\rangle_C}} \qquad [35]$$

We immediately identify the second term to be $\frac{1}{1+E_{\mathrm{o}}}$. Defining $\delta r^\mu = r^\mu - \langle r^\mu\rangle_C$, we can rewrite the first term as

$$\frac{\langle r^\mu\rangle_C^2}{\langle r^\mu\rangle_C^2 + \langle \delta r^{\mu 2}\rangle_C} = \frac{1}{1 + \frac{\langle \delta r^{\mu 2}\rangle_C}{\langle r^\mu\rangle_C^2}} = \frac{1}{1+E_{\mathrm{s}}} \qquad [36]$$

Therefore

$$\frac{d^\top K^+ d}{\hat{\sigma}^2} = \frac{1}{(1+E_{\mathrm{s}})(1+E_{\mathrm{o}})} \qquad [37]$$

which plugged into Eq. (27) gives exactly Eq. (23), as desired.

**C. Derivation of the local-global gap in the regime of finite number of categories.** Here we derive the local-global gap $\Delta E$ in the regime of finite category density, by studying the thermodynamic limit $P, N \to \infty$ at fixed $\alpha = \frac{P}{N} \in (0, \infty)$ with the replica method. Our previous result for infinitely many categories is recovered as the limiting case $\alpha \to \infty$. We recall that we assume that $\mathrm{SNR}^{-1} = \mathcal{O}(N)$ or, equivalently for $\left\langle \sigma_1^{\mu 2}\right\rangle = \mathcal{O}(1)$, $\sum_d \left\langle \sigma_d^{\mu 2}\right\rangle_C = \mathcal{O}(N)$. For this finite-$P$ replica calculation only, we introduce a Gaussian surrogate ensemble realizing the first and second moments assumed in the manifold generative model, Eqs. (19-22). We write

$$u_1^\mu = \sqrt{a}\, u_1 + \sqrt{1-a}\, \eta^\mu, \qquad \eta^\mu \overset{\text{i.i.d.}}{\sim} \mathcal{N}\left(0, \frac{1}{N} I\right),$$

which reproduces the mean and covariance in Eqs. (19-22), and define the remaining directions $u_{d>1}^\mu$ as jointly Gaussian vectors with zero mean and the covariance specified by Eq. (22). This additional Gaussian assumption enables the quenched-disorder average below and is not required for the $P \to \infty$ result.

**_C.1. Gibbs distribution._** To derive the finite-$P$ corrections to the local-global gap, we use the replica method. We start from the training objective

$$\Delta E(W) = \left\langle \left\langle \left(W^\top x^\mu - \hat{y}^\mu\right)^2 \right\rangle_\mu \right\rangle_C , \quad [38]$$

whose minimum is $\Delta E$. We introduce the Gibbs distribution

$$p(W) = \frac{1}{Z} \exp\left(-\beta \tilde{\Delta E}(W)\right), \quad [39]$$

with $\tilde{\Delta E}(W) = \frac{P}{2}\Delta E(W)$ and inverse temperature $\beta \in [0, \infty)$. The corresponding partition function is

$$Z = \int dW \, \exp\left(-\beta \tilde{\Delta E}(W)\right). \quad [40]$$

The average training error under this measure is

$$\langle \Delta E(W) \rangle_{p(W)} = -\frac{2}{P} \frac{\partial \ln Z}{\partial \beta}. \quad [41]$$

In the zero-temperature limit $\beta \to \infty$, the measure $p(W)$ concentrates on the minimizer $W_*$ of $\Delta E(W)$ and therefore

$$\lim_{\beta \to \infty} \langle \Delta E(W) \rangle_{p(W)} = \Delta E(W_*) = \Delta E. \quad [42]$$

The quenched disorder here is given by the random manifold parameters entering $x^\mu$, namely the manifold directions. We assume that the training error is self-averaging, that is it coincides with its disorder average $\Delta E = \langle \Delta E \rangle_C$. Thus, $\Delta E$ can be obtained from the disorder-averaged partition function $\langle \ln Z \rangle_C$ using the replica method.

**_C.2. Auxiliary fields._** We write explicitly the shift term appearing in $\Delta E(W)$ (Eq. (38)) as

$$W^\top x^\mu - \hat{y}^\mu = s_1^\mu \left(\sqrt{a}\, W^\top u_1 + \sqrt{1-a}\, W^\top \eta^\mu - r^\mu\right) + \sum_{d=2}^{N} s_d^\mu W^\top u_d^\mu. \quad [43]$$

This suggests introducing the order parameter

$$m = W^\top u_1, \quad [44]$$

and auxiliary fields

$$v_1^\mu = W^\top \eta^\mu, \qquad v_d^\mu = W^\top u_d^\mu, \quad d = 2, \ldots, N. \quad [45]$$

In terms of these fields, the energy becomes

$$\tilde{\Delta E}(v, m) = \frac{1}{2} \sum_{\mu=1}^{P} \left\langle \left( s_1^\mu \left(\sqrt{a}\, m + \sqrt{1-a}\, v_1^\mu - r^\mu\right) + \sum_{d=2}^{N} s_d^\mu v_d^\mu \right)^2 \right\rangle_\mu . \quad [46]$$

Carrying out the average over the manifold coordinates $s^\mu$, we obtain

$$\begin{aligned} \tilde{\Delta E}(v, m) = \frac{1}{2} \sum_{\mu=1}^{P} \Bigg[ & a\, \sigma_1^{\mu 2} m^2 + 2\sqrt{a(1-a)}\, \sigma_1^{\mu 2} m v_1^\mu + 2\sqrt{a} \sum_{d=2}^{N} \Sigma_{1d}^\mu m v_d^\mu - 2\sqrt{a}\, \sigma_1^{\mu 2} r^\mu m \\ & + (1-a)\sigma_1^{\mu 2} (v_1^\mu)^2 + 2\sqrt{1-a} \sum_{d=2}^{N} \Sigma_{1d}^\mu v_1^\mu v_d^\mu - 2\sqrt{1-a}\, \sigma_1^{\mu 2} r^\mu v_1^\mu + \sum_{d=2}^{N} \sigma_d^{\mu 2} (v_d^\mu)^2 \\ & - 2 r^\mu \sum_{d=2}^{N} \Sigma_{1d}^\mu v_d^\mu + \sigma_1^{\mu 2} (r^\mu)^2 \Bigg]. \end{aligned} \quad [47]$$

We next enforce the definitions of the auxiliary fields through Dirac delta representations. Introducing the conjugate fields $\hat{m}$, $\hat{v}_1^\mu$, and $\hat{v}_d^\mu$, the partition function can be written as

$$Z \propto \int DW \, dm \, d\hat{m} \, Dv \, D\hat{v} \, \exp\left(A(W, m, \hat{m}, v, \hat{v}) - \beta \tilde{\Delta E}(v, m)\right), \quad [48]$$

where

$$A(W,m,\hat{m},v,\hat{v}) = -i\hat{m}\,m + i\hat{m}\,W^\top u_1 - i\sum_{\mu=1}^{P}\hat{v}_1^\mu v_1^\mu + i\sum_{\mu=1}^{P}\hat{v}_1^\mu W^\top \eta^\mu - i\sum_{\mu=1}^{P}\sum_{d=2}^{N}\hat{v}_d^\mu v_d^\mu + i\sum_{\mu=1}^{P}\sum_{d=2}^{N}\hat{v}_d^\mu W^\top u_d^\mu. \quad [49]$$

Here and throughout, for any integration variable $z$ carrying one or more discrete indices, we denote by

$$Dz \equiv \prod_{\{\text{all indices of } z\}} dz \quad [50]$$

the corresponding integration measure. For example,

$$DW = \prod_{k=1}^{N} dW_k, \qquad Dv = \prod_{\mu=1}^{P}\prod_{d=1}^{N} dv_d^\mu, \qquad D\hat{v} = \prod_{\mu=1}^{P}\prod_{d=1}^{N} d\hat{v}_d^\mu. \quad [51]$$

***C.3. Replicas.*** To compute the disorder-averaged free energy, we use the replica identity

$$\langle \ln Z\rangle_C = \lim_{n\to 0}\frac{\langle Z^n\rangle_C - 1}{n}. \quad [52]$$

Replicating the partition function, we obtain

$$\langle Z^n\rangle_C \propto \int DW\,Dm\,D\hat{m}\,Dv\,D\hat{v}\,\left\langle \exp\left(A(\{W^\alpha, m^\alpha, \hat{m}^\alpha, v^\alpha, \hat{v}^\alpha\}) - \beta\sum_{\alpha=1}^{n}\tilde{\Delta E}(v^\alpha, m^\alpha)\right)\right\rangle_C, \quad [53]$$

with

$$A = -i\sum_{\alpha=1}^{n}\hat{m}^\alpha m^\alpha + i\sum_{\alpha=1}^{n}\hat{m}^\alpha W^{\alpha\top}u_1 - i\sum_{\alpha=1}^{n}\sum_{\mu=1}^{P}\hat{v}_1^{\mu\alpha}v_1^{\mu\alpha} + i\sum_{\alpha=1}^{n}\sum_{\mu=1}^{P}\hat{v}_1^{\mu\alpha}W^{\alpha\top}\eta^\mu$$
$$-i\sum_{\alpha=1}^{n}\sum_{\mu=1}^{P}\sum_{d=2}^{N}\hat{v}_d^{\mu\alpha}v_d^{\mu\alpha} + i\sum_{\alpha=1}^{n}\sum_{\mu=1}^{P}\sum_{d=2}^{N}\hat{v}_d^{\mu\alpha}W^{\alpha\top}u_d^\mu, \quad [54]$$

and

$$\tilde{\Delta E}(v^\alpha, m^\alpha) = \frac{1}{2}\sum_{\mu=1}^{P}\Bigg[a\,\sigma_1^{\mu 2}(m^\alpha)^2 + 2\sqrt{a(1-a)}\,\sigma_1^{\mu 2}m^\alpha v_1^{\mu\alpha} + 2\sqrt{a}\sum_{d=2}^{N}\Sigma_{1d}^\mu m^\alpha v_d^{\mu\alpha} - 2\sqrt{a}\,\sigma_1^{\mu 2}r^\mu m^\alpha$$
$$+(1-a)\sigma_1^{\mu 2}(v_1^{\mu\alpha})^2 + 2\sqrt{1-a}\sum_{d=2}^{N}\Sigma_{1d}^\mu v_1^{\mu\alpha}v_d^{\mu\alpha} - 2\sqrt{1-a}\,\sigma_1^{\mu 2}r^\mu v_1^{\mu\alpha} + \sum_{d=2}^{N}\sigma_d^{\mu 2}(v_d^{\mu\alpha})^2$$
$$-2r^\mu\sum_{d=2}^{N}\Sigma_{1d}^\mu v_d^{\mu\alpha} + \sigma_1^{\mu 2}(r^\mu)^2\Bigg]. \quad [55]$$

***C.4. Average over the quenched disorder.*** We now average over the quenched disorder, i.e. over the random manifold directions. Concretely, we average over the random vectors $\eta^\mu$ and $u_d^\mu$ for $d > 1$. This boils down to the calculation of standard Gaussian integrals, at the end of which we obtain

$$\langle Z^n\rangle_C \propto \int DQ\,D\hat{Q}\,DW\,Dm\,D\hat{m}\,Dv\,D\hat{v}\,\exp\left(A(Q,W,m,\hat{m},v,\hat{v}) + B(Q,\hat{Q},W) - \beta\sum_{\alpha=1}^{n}\tilde{\Delta E}(v^\alpha, m^\alpha)\right), \quad [56]$$

where

$$B(Q,\hat{Q},W) = \frac{N}{2}\mathrm{Tr}\,Q^\top\hat{Q} - \frac{1}{2}\sum_{\alpha,\beta=1}^{n}\hat{Q}^{\alpha\beta}W^{\alpha\top}W^\beta, \quad [57]$$

$$A = -i\sum_{\alpha=1}^{n}\hat{m}^\alpha m^\alpha + i\sum_{\alpha=1}^{n}\hat{m}^\alpha W^{\alpha\top}u_1 - i\sum_{\alpha=1}^{n}\sum_{\mu=1}^{P}\hat{v}_1^{\mu\alpha}v_1^{\mu\alpha} - i\sum_{\alpha=1}^{n}\sum_{\mu=1}^{P}\sum_{d=2}^{N}\hat{v}_d^{\mu\alpha}v_d^{\mu\alpha}$$
$$-\frac{1-a}{2}\sum_{\mu=1}^{P}\sum_{\alpha,\beta=1}^{n}\hat{v}_1^{\mu\alpha}Q^{\alpha\beta}\hat{v}_1^{\mu\beta} - \frac{1}{2}\sum_{\mu=1}^{P}\sum_{d=2}^{N}\sum_{\alpha,\beta=1}^{n}\hat{v}_d^{\mu\alpha}\left(Q^{\alpha\beta} - \frac{a}{N}m^\alpha m^\beta\right)\hat{v}_d^{\mu\beta}, \quad [58]$$

and, by means of Dirac delta representations, we introduced the overlap order parameters

$$Q^{\alpha\beta} = \frac{1}{N} W^{\alpha\top} W^{\beta}, \tag{59}$$

and their conjugates $\hat{Q}^{\alpha\beta}$.

***C.5. Gaussian integrals in the auxiliary fields.*** We now perform, in this order, the Gaussian integrals over $\hat{v}$, $W$, and $\hat{m}$. Neglecting additive constants independent of $\beta$, and dropping the subleading $\mathcal{O}(1)$ log-determinant contribution generated by the $\hat{m}$ integral, we obtain

$$\langle Z^n \rangle_C \propto \int DQ\, D\hat{Q}\, Dm\, Dv\, \exp\left( A(Q, \hat{Q}, m, v) + B(Q, \hat{Q}, m) - \beta \sum_{\alpha=1}^{n} \tilde{\Delta E}(v^\alpha, m^\alpha) \right), \tag{60}$$

with

$$A = -\frac{1}{2} \sum_{\alpha,\beta=1}^{n} m^\alpha \hat{Q}^{\alpha\beta} m^\beta - \frac{1}{2}(1-a)^{-1} \sum_{\mu=1}^{P} \sum_{\alpha,\beta=1}^{n} v_1^{\mu\alpha} \left(Q^{-1}\right)^{\alpha\beta} v_1^{\mu\beta}$$

$$-\frac{1}{2} \sum_{\mu=1}^{P} \sum_{d=2}^{N} \sum_{\alpha,\beta=1}^{n} v_d^{\mu\alpha} \left[ Q^{-1} + \frac{a}{N} \frac{Q^{-1} m m^\top Q^{-1}}{1 - \frac{a}{N} m^\top Q^{-1} m} \right]^{\alpha\beta} v_d^{\mu\beta}, \tag{61}$$

and

$$B = \frac{N}{2} \mathrm{Tr}\, Q^\top \hat{Q} - \frac{1}{2} PN \ln\det(Q) - \frac{1}{2} NP \ln\left(1 - \frac{a}{N} m^\top Q^{-1} m\right) - \frac{1}{2} N \ln\det(\hat{Q}), \tag{62}$$

where we used the identities

$$\left(Q - \frac{a}{N} m m^\top\right)^{-1} = Q^{-1} + \frac{a}{N} \frac{Q^{-1} m m^\top Q^{-1}}{1 - \frac{a}{N} m^\top Q^{-1} m}, \tag{63}$$

and

$$\det\left(Q - \frac{a}{N} m m^\top\right) = \left(1 - \frac{a}{N} m^\top Q^{-1} m\right) \det(Q). \tag{64}$$

***C.6. Replica symmetric ansatz.*** Assuming a replica-symmetric saddle point, we make the ansatz

$$Q = (q_1 - q_0)\, \mathbb{I} + q_0 \mathbf{1}, \qquad \hat{Q} = (\hat{q}_1 - \hat{q}_0)\, \mathbb{I} + \hat{q}_0 \mathbf{1}, \qquad m^\alpha = m, \tag{65}$$

Using standard identities for replica-symmetric matrices, we then obtain

$$A(Q, \hat{Q}, m, v) = -\frac{1}{2} n\, (\hat{q}_1 - \hat{q}_0)\, m^2 - \frac{1}{2} \frac{(1-a)^{-1}}{q_1 - q_0} \sum_{\mu=1}^{P} \sum_{\alpha=1}^{n} (v_1^{\mu\alpha})^2 + \frac{1}{2} \frac{(1-a)^{-1} q_0}{(q_1 - q_0)^2} \sum_{\mu=1}^{P} \left( \sum_{\alpha=1}^{n} v_1^{\mu\alpha} \right)^2$$

$$-\frac{1}{2} \frac{1}{q_1 - q_0} \sum_{\mu=1}^{P} \sum_{d=2}^{N} \sum_{\alpha=1}^{n} (v_d^{\mu\alpha})^2 + \frac{1}{2} \frac{q_0}{(q_1 - q_0)^2} \sum_{\mu=1}^{P} \sum_{d=2}^{N} \left( \sum_{\alpha=1}^{n} v_d^{\mu\alpha} \right)^2 - \frac{1}{2} \frac{a}{N} \frac{m^2}{(q_1 - q_0)^2} \sum_{\mu=1}^{P} \sum_{d=2}^{N} \left( \sum_{\alpha=1}^{n} v_d^{\mu\alpha} \right)^2, \tag{66}$$

and

$$B(Q, \hat{Q}, m) = \frac{Nn}{2} \left[ (q_1 - q_0)(\hat{q}_1 - \hat{q}_0) + (q_1 - q_0)\hat{q}_0 + (\hat{q}_1 - \hat{q}_0) q_0 \right]$$

$$-\frac{1}{2} PN\, n \left[ \ln(q_1 - q_0) + \frac{q_0}{q_1 - q_0} \right] - \frac{1}{2} N\, n \left[ \ln(\hat{q}_1 - \hat{q}_0) + \frac{\hat{q}_0}{\hat{q}_1 - \hat{q}_0} \right] + \frac{1}{2} NP \frac{a}{N}\, n\, \frac{m^2}{q_1 - q_0}. \tag{67}$$

where we retained only terms up to linear order in $n$.

***C.7. Remaining Gaussian integrals.*** Note that Eq. (66) contains replica-coupling terms of the form $(\sum_\alpha v_1^{\mu\alpha})^2$ and $(\sum_\alpha v_d^{\mu\alpha})^2$. We decouple these terms by standard Hubbard-Stratonovich transformations, introducing Gaussian auxiliary fields $t_1^\mu$ and $t_d^\mu$ for $d = 2, \ldots, N$. At fixed $t$, the integrals over the replicated variables factorize over $\alpha$. Since the replica limit $n \to 0$ is taken at the end, we retain only the linear term in $n$, using

$$\int Dt^\mu \left[Z_\mu(t^\mu)\right]^n = 1 + n \int Dt^\mu \ln Z_\mu(t^\mu) + \mathcal{O}(n^2), \tag{68}$$

where $Dt^\mu$ denotes the standard Gaussian measure over the auxiliary fields associated with manifold $\mu$. We then perform sequentially the Gaussian integrals over $v_d^\mu$ for $d = 2, \ldots, N$, over $v_1^\mu$, and finally over the auxiliary fields $t^\mu$. This yields

$$\langle Z^n \rangle_C \propto \int dm\, dq_0\, d\delta q\, d\hat{q}_0\, d\delta\hat{q}\, \exp\left(S_{\mathrm{RS}}(m, q_0, \delta q, \hat{q}_0, \delta\hat{q})\right), \tag{69}$$

where

$$\begin{aligned}
\frac{2}{Pn} S_{\mathrm{RS}}(m, q_0, \delta q, \hat{q}_0, \delta\hat{q}) &= -\beta \frac{1}{P} \sum_{\mu=1}^{P} \sigma_1^{\mu 2} r^{\mu 2} - \beta a \left\langle \sigma_1^{\mu 2} \right\rangle_C m^2 + 2\beta\sqrt{a} \frac{1}{P} \sum_{\mu=1}^{P} r^\mu \sigma_1^{\mu 2} m - \frac{\beta}{P} \delta\hat{q}\, m^2 \\
&+ \alpha^{-1} \left[\delta q\, \delta\hat{q} + \beta^{-1} \delta q\, \hat{q}_0 + \beta\, \delta\hat{q}\, q_0\right] - \alpha^{-1} \left[\ln(\beta\delta\hat{q}) + \frac{\hat{q}_0}{\beta\delta\hat{q}}\right] \\
&- N \left[\ln(\beta^{-1}\delta q) + \frac{\beta q_0}{\delta q}\right] + \frac{\beta a\, m^2}{\delta q} - \frac{1}{P} \sum_{\mu=1}^{P} \sum_{d=2}^{N} \ln \beta U_d^\mu + \frac{1}{P} \beta \sum_{\mu=1}^{P} \sum_{d=2}^{N} S_d^\mu\, (\delta_m^\mu)^2 + \frac{1}{P} \beta \sum_{\mu=1}^{P} \sum_{d=2}^{N} (U_d^\mu)^{-1} V_2 \\
&- \frac{1}{P} \sum_{\mu=1}^{P} \ln \left(\beta U_1^\mu - \beta \sum_{d=2}^{N} S_d^\mu\right) + \frac{1}{P} \beta \sum_{\mu=1}^{P} \left(U_1^\mu - \sum_{d=2}^{N} S_d^\mu\right)^{-1} \left(\sigma_1^{\mu 2} - \sum_{d=2}^{N} S_d^\mu\right)^2 (\delta_m^\mu)^2 \\
&+ \frac{1}{P} \beta \sum_{\mu=1}^{P} \left(U_1^\mu - \sum_{d=2}^{N} S_d^\mu\right)^{-1} V_1 + \frac{1}{P} \beta \sum_{\mu=1}^{P} \left(U_1^\mu - \sum_{d=2}^{N} S_d^\mu\right)^{-1} V_2 \sum_{d=2}^{N} \left((U_d^\mu)^{-1} \Sigma_{1d}^\mu\right)^2,
\end{aligned} \tag{70}$$

with

$$U_1^\mu = \sigma_1^{\mu 2} + \frac{(1-a)^{-1}}{\delta q}, \qquad U_d^\mu = \sigma_d^{\mu 2} + \frac{1}{\delta q}, \qquad d = 2, \ldots, N, \tag{71}$$

$$V_1 = \frac{(1-a)^{-1} q_0}{\delta q^2}, \qquad V_2 = \frac{q_0}{\delta q^2} - \frac{a}{N} \frac{m^2}{\delta q^2}, \tag{72}$$

$$S_d^\mu = (U_d^\mu)^{-1} (\Sigma_{1d}^\mu)^2, \qquad \delta_m^\mu = r^\mu - \sqrt{a}\, m. \tag{73}$$

Note that we performed the change of variables

$$\beta^{-1} \delta q = q_1 - q_0 \tag{74}$$

$$\beta \delta\hat{q} = \hat{q}_1 - \hat{q}_0 \tag{75}$$

while keeping $q_0$ and $\hat{q}_0$ unchanged.

***C.8. Saddle point.*** In the limit $N, P \to \infty$, with $\frac{P}{N} = \alpha \in [0, \infty]$, the integral in (69) can be solved via saddle point. We note that $S_{\mathrm{RS}}$ is linear in $q_0$ and $\hat{q}_0$, so we do not need to compute the saddle point solution for $q_0$ and $\hat{q}_0$ explicitly, as any term of $S_{\mathrm{RS}}$ that is linear in these variables vanishes under the saddle point condition. The saddle point equation w.r.t. $\hat{q}_0$ gives

$$\delta q = \delta\hat{q}^{-1} \tag{76}$$

The saddle point equation w.r.t. $q_0$ gives the self-consistent equation for $\delta\hat{q}$

$$\boxed{\delta\hat{q} \frac{1}{N} \frac{1}{P} \sum_{\mu} \left\{ \sum_{d=2}^{N} (U_d^\mu)^{-1} + (1-a)^{-1} \left[U_1^\mu - \sum_{d=2}^{N} S_d^\mu\right]^{-1} + \left[U_1^\mu - \sum_{d=2}^{N} S_d^\mu\right]^{-1} \sum_{d=2}^{N} \left((U_d^\mu)^{-1} \Sigma_{1d}^\mu\right)^2 \right\} = 1 - \frac{1}{N} \alpha^{-1}} \tag{77}$$

***C.9. Order parameter in the*** $\alpha \to \infty$ ***limit.*** It is useful to study the order parameter in the limit $\alpha \to \infty$. We rewrite the self-consistent equation Eq. (77)

$$\frac{1}{N}\frac{1}{P}\sum_{\mu}\left\{\sum_{d=2}^{N}\frac{\delta\hat{q}}{\sigma_d^{\mu 2}+\delta\hat{q}}+\frac{(1-a)^{-1}\delta\hat{q}}{\sigma_1^{\mu 2}+(1-a)^{-1}\delta\hat{q}-\sum_{d>1}\frac{(\Sigma_{1d}^{\mu})^2}{\sigma_d^{\mu 2}+\delta\hat{q}}}\right.$$
$$\left.+\frac{\delta\hat{q}}{\sigma_1^{\mu 2}+(1-a)^{-1}\delta\hat{q}-\sum_{d>1}\frac{(\Sigma_{1d}^{\mu})^2}{\sigma_d^{\mu 2}+\delta\hat{q}}}\sum_{d=2}^{N}\left(\frac{\Sigma_{1d}^{\mu}}{\sigma_d^{\mu 2}+\delta\hat{q}}\right)^2\right\}=1-\frac{1}{N}\alpha^{-1} \quad [78]$$

It is readily verified that $\delta\hat{q}\overset{\alpha\to\infty}{\to}\infty$. Indeed, taking this limit we get the identity

$$\frac{1}{P}\frac{1}{N}\sum_{\mu}\left\{\sum_{d=2}^{N}1+1+\mathcal{O}\left(\frac{1}{\delta\hat{q}}\right)\right\}=1+\mathcal{O}\left(\alpha^{-1}\right) \quad [79]$$

To determine the asymptotic behavior of $\delta\hat{q}$, we expand the l.h.s of Eq. (78) for $\delta\hat{q}\to\infty$. Term by term, we have

$$\frac{1}{\frac{\sigma_d^{\mu 2}}{\delta\hat{q}}+1}\sim 1-\frac{1}{\delta\hat{q}}\sigma_d^{\mu 2} \quad [80]$$

$$\frac{1}{1+\frac{\sigma_1^{\mu 2}}{(1-a)^{-1}\delta\hat{q}}+\mathcal{O}\left(\frac{1}{\delta\hat{q}^2}\right)}\sim 1-\frac{1}{\delta\hat{q}}(1-a)\,\sigma_1^{\mu 2} \quad [81]$$

while the last term is of subleading order $\mathcal{O}\left(\frac{1}{\delta\hat{q}^2}\right)$. Plugging this back into Eq. (78) we get

$$\delta\hat{q}\overset{\alpha\to\infty}{\sim}\alpha\left[(1-a)\left\langle\sigma_1^{\mu 2}\right\rangle_C+\sum_{d>1}\left\langle\sigma_d^{\mu 2}\right\rangle_C\right] \quad [82]$$

Note that, at least for large $\alpha$, the order parameter is proportional to $\sum_{d>1}\left\langle\sigma_d^{\mu 2}\right\rangle_C=\mathcal{O}(N)$. This will be useful to determine which terms can be neglected in the final expression for $\Delta E$, because of subleading order in $N$, and which should be kept.

***C.10. Local-global gap at the saddle point.*** At the saddle point, the local-global gap is given by

$$\Delta E=\min_m\left[\Delta E(m)\right] \quad [83]$$

where

$$\Delta E(m)=-\frac{1}{n}\frac{2}{P}\frac{1}{\beta}S_{\mathrm{RS}}=\frac{1}{P}\sum_{\mu=1}^{P}\sigma_1^{\mu 2}r^{\mu 2}+a\left\langle\sigma_1^{\mu 2}\right\rangle_C m^2-2\sqrt{a}m\,\frac{1}{P}\sum_{\mu=1}^{P}r^{\mu}\sigma_1^{\mu 2}+\frac{1}{P}\delta\hat{q}\,m^2 \quad [84]$$

$$-\delta\hat{q}\,a\,m^2-\frac{1}{P}\sum_{\mu=1}^{P}\sum_{d=2}^{N}S_d^{\mu}\left(r^{\mu}-\sqrt{a}\,m\right)^2+\frac{1}{N}\delta\hat{q}^2am^2\frac{1}{P}\sum_{\mu=1}^{P}\sum_{d=2}^{N}(U_d^{\mu})^{-1} \quad [85]$$

$$-\frac{1}{P}\sum_{\mu=1}^{P}\left(U_1^{\mu}-\sum_{d=2}^{N}S_d^{\mu}\right)^{-1}\left(\sigma_1^{\mu 2}-\sum_{d=2}^{N}S_d^{\mu}\right)^2\left(r^{\mu}-\sqrt{a}\,m\right)^2 \quad [86]$$

$\delta\hat{q}$ is determined by the self-consistent equation Eq. (77), and we have kept only those terms that are of leading order in both $\beta$ as $\beta\to\infty$, and in $N$ or $P$, in the limit $N,P\to\infty$, with $\frac{P}{N}=\alpha\in[0,\infty]$. Note that some terms in $\Delta E(m)$, such as $\frac{1}{P}\delta\hat{q}\,m^2$, may appear to be of subleading order in this limit, but are in fact to be kept because, at least for large $\alpha$, the order parameter $\delta\hat{q}$ is of $\mathcal{O}(N)$.

Observing that Eq. (86) is quadratic in $m$, its minimization is readily obtained as

$$\Delta E = -\frac{k_1^2}{4k_2} + k_0 \quad [87]$$

with

$$k_2 = a\left\langle \sigma_1^{\mu 2}\right\rangle_C + \frac{1}{P}\delta\hat{q} - a\delta\hat{q} - a\frac{1}{P}\sum_{\mu=1}^{P}\sum_{d=2}^{N} S_d^\mu + \frac{a}{N}(\delta\hat{q})^2\frac{1}{P}\sum_{\mu=1}^{P}\sum_{d=2}^{N}(U_d^\mu)^{-1}$$
$$- a\frac{1}{P}\sum_\mu\left(U_1^\mu - \sum_{d=2}^{N} S_d^\mu\right)^{-1}\left(\sigma_1^{\mu 2} - \sum_{d=2}^{N} S_d^\mu\right)^2 \quad [88]$$

$$k_1 = -2\sqrt{a}\frac{1}{P}\sum_{\mu=1}^{P} r^\mu\sigma_1^{\mu 2} + 2\sqrt{a}\frac{1}{P}\sum_{\mu=1}^{P}\sum_{d=2}^{N} S_d^\mu r^\mu + 2\sqrt{a}\frac{1}{P}\sum_\mu\left(U_1^\mu - \sum_{d=2}^{N} S_d^\mu\right)^{-1}\left(\sigma_1^{\mu 2} - \sum_{d=2}^{N} S_d^\mu\right)^2 r^\mu \quad [89]$$

$$k_0 = \frac{1}{P}\sum_{\mu=1}^{P}\sigma_1^{\mu 2} r^{\mu 2} - \frac{1}{P}\sum_{\mu=1}^{P}\sum_{d=2}^{N} S_d^\mu r^{\mu 2} - \frac{1}{P}\sum_{\mu=1}^{P}\left(U_1^\mu - \sum_{d=2}^{N} S_d^\mu\right)^{-1}\left(\sigma_1^{\mu 2} - \sum_{d=2}^{N} S_d^\mu\right)^2 r^{\mu 2} \quad [90]$$

***C.11. Local-global gap in the $\alpha \to \infty$ limit.*** It is instructive to check that our expression for the local-global gap in the case of infinite categories, i.e. $\alpha \to \infty$, is recovered. As $\delta\hat{q} \to \infty$, most terms in Eqs. (88-90) that depend on $\delta\hat{q}$ vanish, except for the following three terms in Eqs. (88)

$$+\frac{1}{P}\delta\hat{q} \sim \frac{\alpha}{P}\left[(1-a)\left\langle\sigma_1^{\mu 2}\right\rangle_C + \sum_{d>1}\left\langle\sigma_d^{\mu 2}\right\rangle_C\right] = \frac{1}{N}\left[(1-a)\left\langle\sigma_1^{\mu 2}\right\rangle_C + \sum_{d>1}\left\langle\sigma_d^{\mu 2}\right\rangle_C\right] \quad [91]$$

$$-a\delta\hat{q} \quad [92]$$

$$+\frac{a}{N}(\delta\hat{q})^2\frac{1}{P}\sum_\mu\sum_{d>1}(U_d^\mu)^{-1} = \frac{a}{N}\frac{1}{P}\sum_\mu\sum_{d>1}\delta\hat{q}\left[\frac{1}{1+\frac{\sigma_d^{\mu 2}}{\delta\hat{q}}}\right] \sim a\delta\hat{q} - \frac{a}{N}\sum_{d>1}\left\langle\sigma_d^{\mu 2}\right\rangle_C \quad [93]$$

The second term, which is diverging, cancels with part of the third, and we have (assuming $r^\mu$ and $\sigma_1^\mu$ to be uncorrelated as in our infinite-$P$ theory)

$$k_2 = a\left\langle\sigma_1^{\mu 2}\right\rangle_C + (1-a)\frac{1}{N}\sum_{d>1}\left\langle\sigma_d^{\mu 2}\right\rangle_C \quad [94]$$

$$k_1 = -2\sqrt{a}\left\langle r^\mu\right\rangle_C\left\langle\sigma_1^{\mu 2}\right\rangle_C \quad [95]$$

$$k_0 = \left\langle\sigma_1^{\mu 2}\right\rangle_C\left\langle r^{\mu 2}\right\rangle_C \quad [96]$$

which plugged into Eq. (87) give exactly our prediction for $\Delta E$ in the limit $\alpha \to \infty$, Eq. (23).

**D. Numerical validation.** In the main text, we tested the theory predictions against the *test* regression error measured from the empirical data and found good agreement. Here, instead, we validate the theory in the setting for which it is derived: the *training* regression error, for manifolds generated under the theory generative model. In this case, we expect virtually perfect agreement up to finite-size corrections, which is indeed what we observe (**Fig. S28**).

The generative model still leaves freedom in the choice of its parameters, including the alignment $a$, the local scales $r^\mu$, and the manifold covariances $\Sigma^\mu$. In particular, the entries $\sigma_1^{\mu 2}$ and $\sigma_{d>1}^{\mu 2}$ contribute to the SNR, while the off-diagonal terms $\Sigma_{1d}^\mu$ determine how the feature-encoding direction is embedded within each manifold. To test the theory in a realistic regime, we chose these parameters to match those measured from the category manifolds at the feature layer of network CR (and the corresponding local feature-encoding directions for a given bounding-box feature). Concretely, we sampled $u_1^\mu = \sqrt{a}\,u_1 + \sqrt{1-a}\,\eta^\mu$,with $u_1$ and $a$ estimated from the data. Similarly, we sampled $r^\mu \sim \mathcal{N}\big(\mathrm{Avg}_\mu[r^\mu], \mathrm{Var}_\mu[r^\mu]\big)$, and we took the covariance matrices $\Sigma^\mu$ directly from the empirical category manifolds. When $P$ exceeded the 265 category manifolds available from the data, we sampled $\Sigma^\mu$ uniformly from these 265 empirical covariances for categories with $\mu > 265$. **Fig. S28** shows the local-global error gap as measured from this set of category manifolds, compared against the infinite-$P$ and finite-$P$ theory.

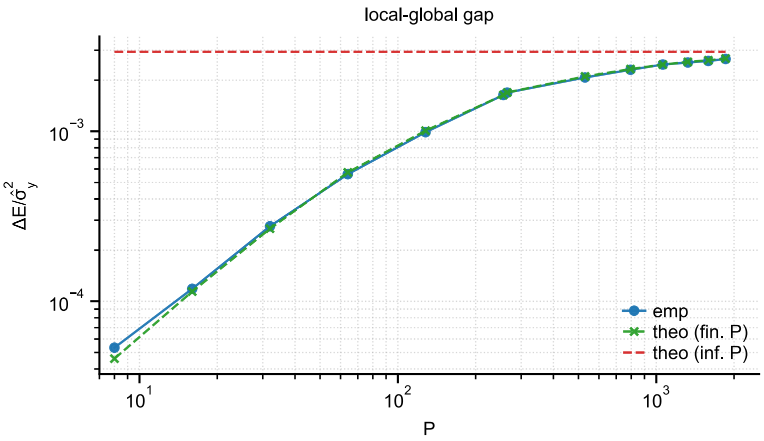

**Fig. S28.** Theory validation. Training local-global error gap for category manifolds generated with the methods described in section D, for bounding-box feature $L_h$. Blue: measured gap; green: finite-$P$ theory; red line: infinite-$P$ theory.
